# Perturbed pediatric serum metabolome in mild and severe dengue disease

**DOI:** 10.1101/2024.07.16.603788

**Authors:** Paul S. Soma, Rebekah C. Gullberg, Barbara Graham, M. Nurul Islam, Angel Balmaseda, Carol D. Blair, Barry J. Beaty, John T. Belisle, Eva Harris, Rushika Perera

## Abstract

Dengue viruses (DENVs) are the most prevalent arboviruses affecting humans. Four billion people are at risk of infection and this burden is rapidly increasing due to geographic expansion of the mosquito vector. Infection with any of the four serotypes of DENV can result in a self-limiting disease but debilitating febrile illness (DF), and some infections progress to severe disease with manifestations such as hemorrhage and shock. DENV infection drives the metabolic state of host cells for viral benefit and induces a host-immune response that has metabolic implications that link to disease. In this study, a dynamic metabolic response to DENV infection and disease was measured in 535 pediatric patient sera using liquid chromatography-mass spectrometry. The metabolome was interrogated to discover biochemical pathways and identify key metabolites perturbed in severe dengue disease. A biomarker panel of thirty-two perturbed metabolites was utilized to classify DF, and severe dengue hemorrhagic fever (DHF) and dengue shock syndrome (DSS) with high sensitivity and specificity equating to a balanced accuracy of 96.9%. Some metabolites that were structurally confirmed here belong to important biochemical pathways of omega-3 and omega-6 fatty acids, sphingolipids, purines, and tryptophan metabolism. A previously reported trend between serotonin and platelets in DHF patients has been expanded upon here to reveal a major depletion of serotonin, but not platelets, in DSS patients. This study differentiated and classified DF and DHF/DSS using a serum metabolic biomarker panel based on perturbed biochemical pathways that have potential implications for severe dengue disease.

**One sentence summary:** Metabolic biomarkers distinguish dengue hemorrhagic fever and dengue shock syndrome from dengue fever and lend insight to severe disease pathology

## INTRODUCTION

Dengue viruses (DENV) are mosquito-borne flaviviruses that place 3.97 billion people at risk of infection each year, with up to 390 million infections annually, rendering them the most prevalent arboviruses worldwide(*1–3*). In the first 5 months of 2024 there were 7.6 million reported dengue cases including 16,000 cases of severe disease and 3,000 deaths, quickly surpassing the previous annual high of 4.6 million dengue cases during 2023(*4*). There are four DENV serotypes (DENV1-4), and while infection with one serotype can cross-protect from infection with a heterologous serotype in the short term, it can lead to antibody-dependent enhancement of disease in the longer term(*5*). DENV is the etiologic agent of dengue fever (DF), which is an incapacitating but self-limited disease, but some cases progress to the potentially fatal dengue hemorrhagic fever (DHF) or dengue shock syndrome (DSS). Peak viremia and fever/symptom onset occur several days after the human host is bitten by an infected mosquito, the acute phase lasts 7 days and includes the ‘critical phase’ 4 to 7 days after fever onset(*6*). Most patients present with DF and defervesce during the critical phase. In a minority of cases, severe manifestations present during the critical phase with the hallmark vascular leakage that can lead to shock and lethal outcomes. Various proteins, peptides, and metabolites have been previously proposed as severe dengue biomarkers(*7–14*), but metabolite biomarkers hold notable advantages, and the study of metabolism presents an opportunity to help resolve challenges in biomarker discovery.

Metabolites within host cells are small molecule intermediates and products of biochemical reactions in the body. These metabolic reactions are driven by upstream gene expression (transcription and translation) and enzyme activity, as well as the current cellular environment such as active metabolic signaling and bystander effects(*15–17*). The metabolome is known as an effector of phenotype because it is dynamic and responds to external stimuli(*18*). Therefore, the metabolome may describe the current physiological state of the system under study with greater temporal resolution than upstream biomolecules. Perturbations in cellular metabolism due to viral infection and the host immune response are reflected in the host’s serum metabolite profile. Accordingly, measuring the serum metabolome of DENV infected patients can provide information about system wide metabolic shifts upon DENV infection and onset of disease.

Infection with DENV prompts perturbations in host cellular metabolism to facilitate various stages of the viral lifecycle and induces a host immune response that is also associated with metabolic alterations(*11–13, 19, 20*). Metabolomic measurements that lead to mapping the dysregulated pathways in DENV infection and disease could improve understanding of pathogenic processes, define enzyme drug targets that, when inhibited, interfere with the viral lifecycle, or identify biomarkers of severe dengue disease that aid in triage. In this study, the DENV-induced metabolic perturbations in pediatric patient sera were measured using liquid chromatography-mass spectrometry (LC-MS). The measured dysregulated metabolome was used for disease state classification, metabolic pathway analysis, and exploration of the biochemistry of dengue disease pathology.

## RESULTS

### Clinical samples

For this study 535 retrospective serum samples from individual pediatric patients enrolled in two well-established studies in Managua, Nicaragua (see Materials and Methods for details) were used. Selected patient demographics are summarized in Figure 1A-E. Statistical assessment of patient demographics between disease states (clinical diagnosis) is included in Table S1. Of the 535 suspected dengue cases, 251 were laboratory-confirmed as dengue-positive, and 284 were non-dengue (ND) febrile illnesses. Based the 1997 World Health Organization disease severity criteria(*21*), the dengue cases were classified as DF (185), DHF (44), and DSS (22). The combination of DHF and DSS patients will be referred to as ‘severe dengue disease’. The study included pediatric patients of ages 1 to 16 years (Figure 1A) and similar numbers of females (n = 257, 48%) and males (n = 278, 52%) (Figure 2B). All samples used for this study were collected at day 1 to 7 of illness (Figure 1C). Dengue patients were positive for DENV serotypes 1, 2 or 3, or an undetermined serotype (Figure 1D). Immune status (primary or secondary infection) was reported (Figure 1E).

**Figure 1.**
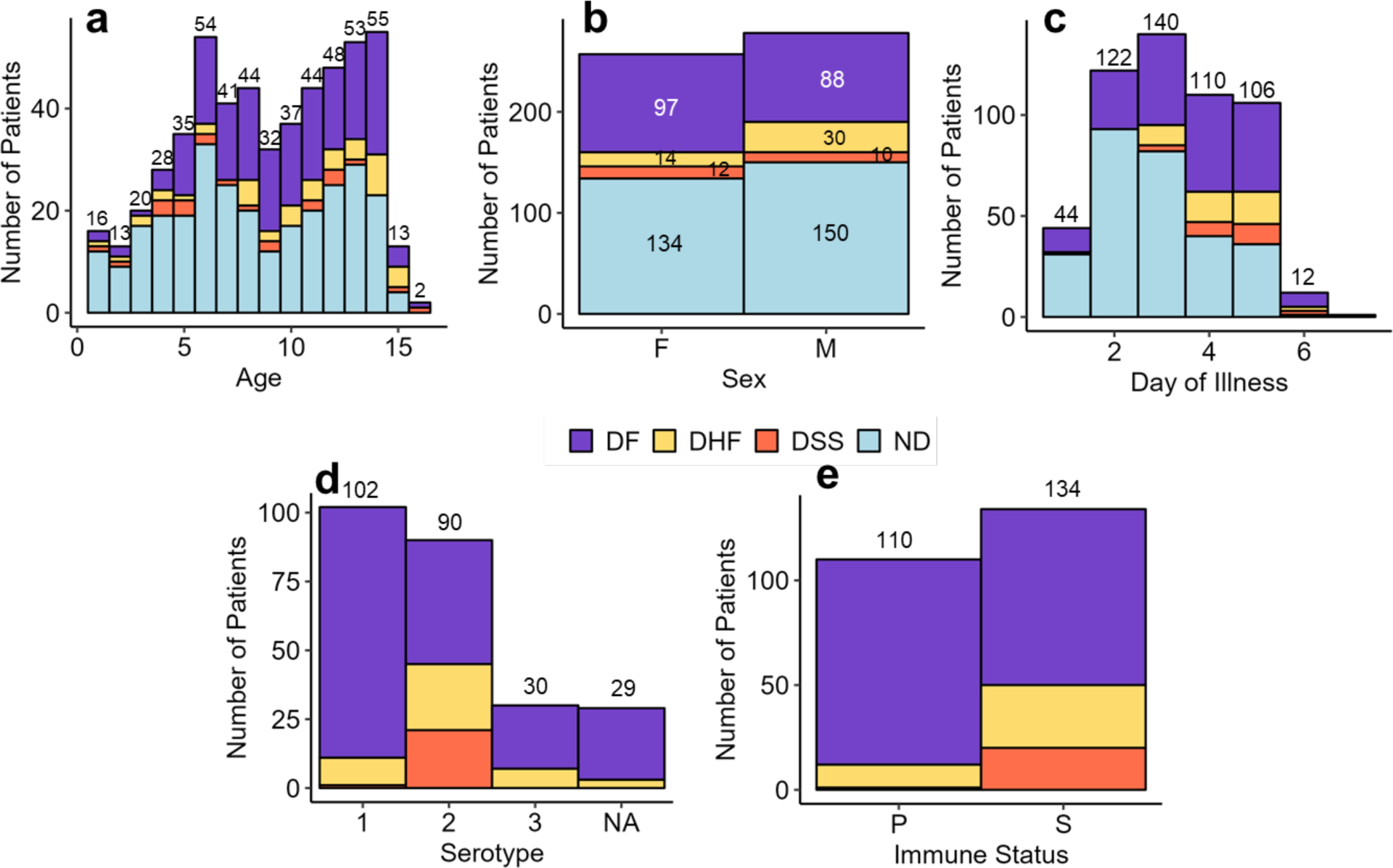
Summary of selected pediatric patient metadata. A) patient age, B) sex, C) day of illness, D) DENV serotype and E) immune status, primary (P), secondary (S). Colors represent disease outcomes: DF, DHF, DSS or ND. Numbers above the bars for age, day of illness, serotype, and infection history (primary or secondary infection) represent the total number of pediatric patients in each group. Numbers within each color bar for sex represent the number of male and female patients with the respective disease outcome.

**Figure 2.**
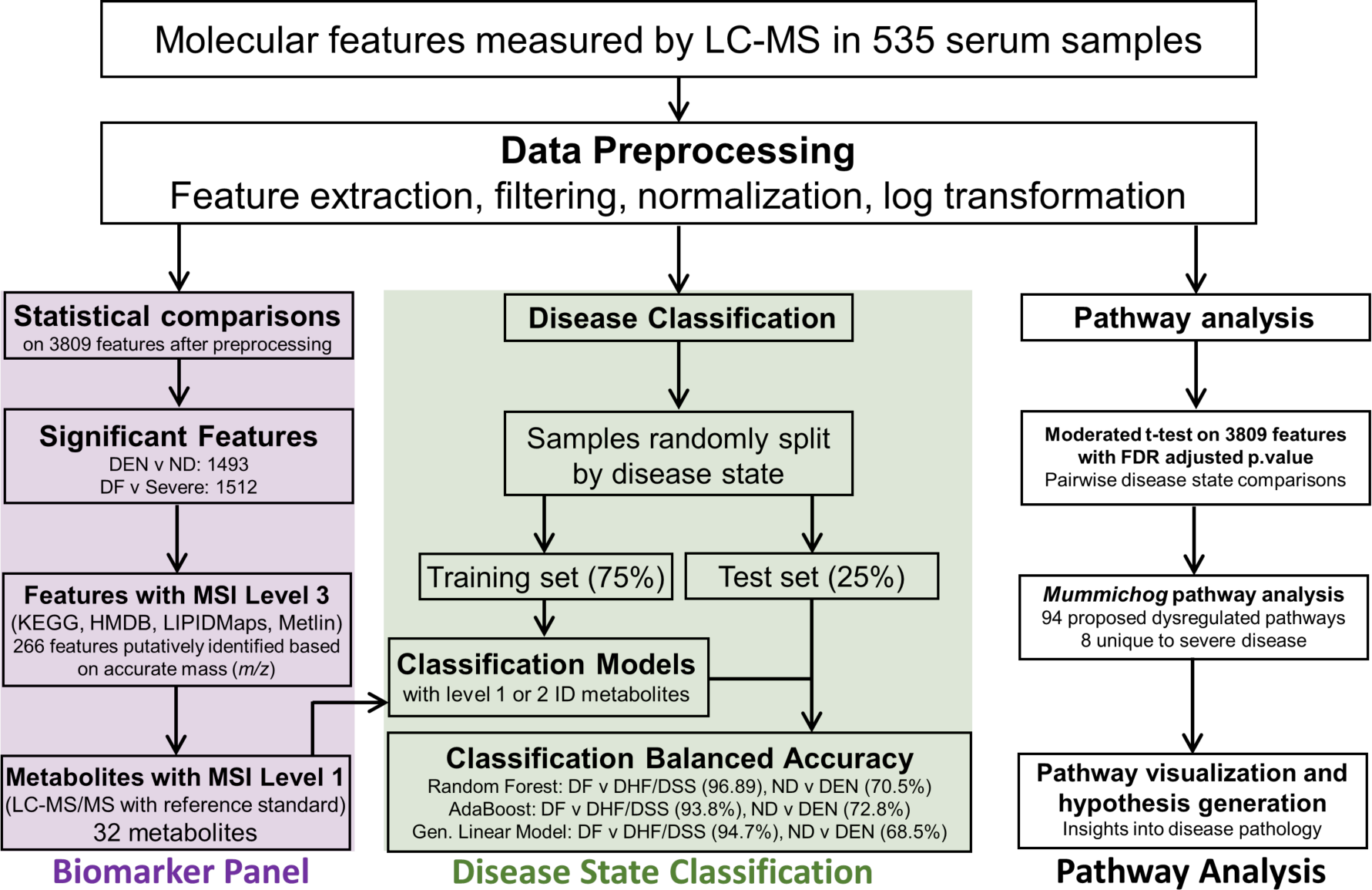
Data processing and analysis workflow, and summarized results for the serum biomarker panel, disease state classification and pathway analysis.

### Serum metabolomics

The serum metabolome of each patient was measured *via* untargeted LC-MS, and the workflow to identify molecular features of significant differential abundance across disease states is shown in Figure 2. A total of 1512 features revealed significant (*adjusted p-value < 0.05*) differential abundance between three pairwise disease state comparisons: DFvsDSS, DHFvsDSS and DFvsDHF. Volcano plots were used to visualize the differential abundance of features between disease states (Figure 3 and Figure S1), and histograms were used to visualize the frequency of features across log_2_FC values (Figure S2). In the pairwise comparison of DFvsDSS, 1117 and 160 features were identified as less abundant (*log_2_FC < −*1) and more abundant (log_2_FC > *1*), respectively, in DSS, and 235 features with significant p values presented a log_2_FC of < 1 or > −1 and were considered not changed. Comparing DHFvsDSS resulted in 961 and 320 features that were less or more abundant, respectively in DSS and 231 that were not changed. Comparing DFvsDHF resulted in 364 features that were less abundant in DHF, 31 features that were more abundant in DHF and 1117 that met the defined criteria for ‘not changed’.

**Figure 3.**
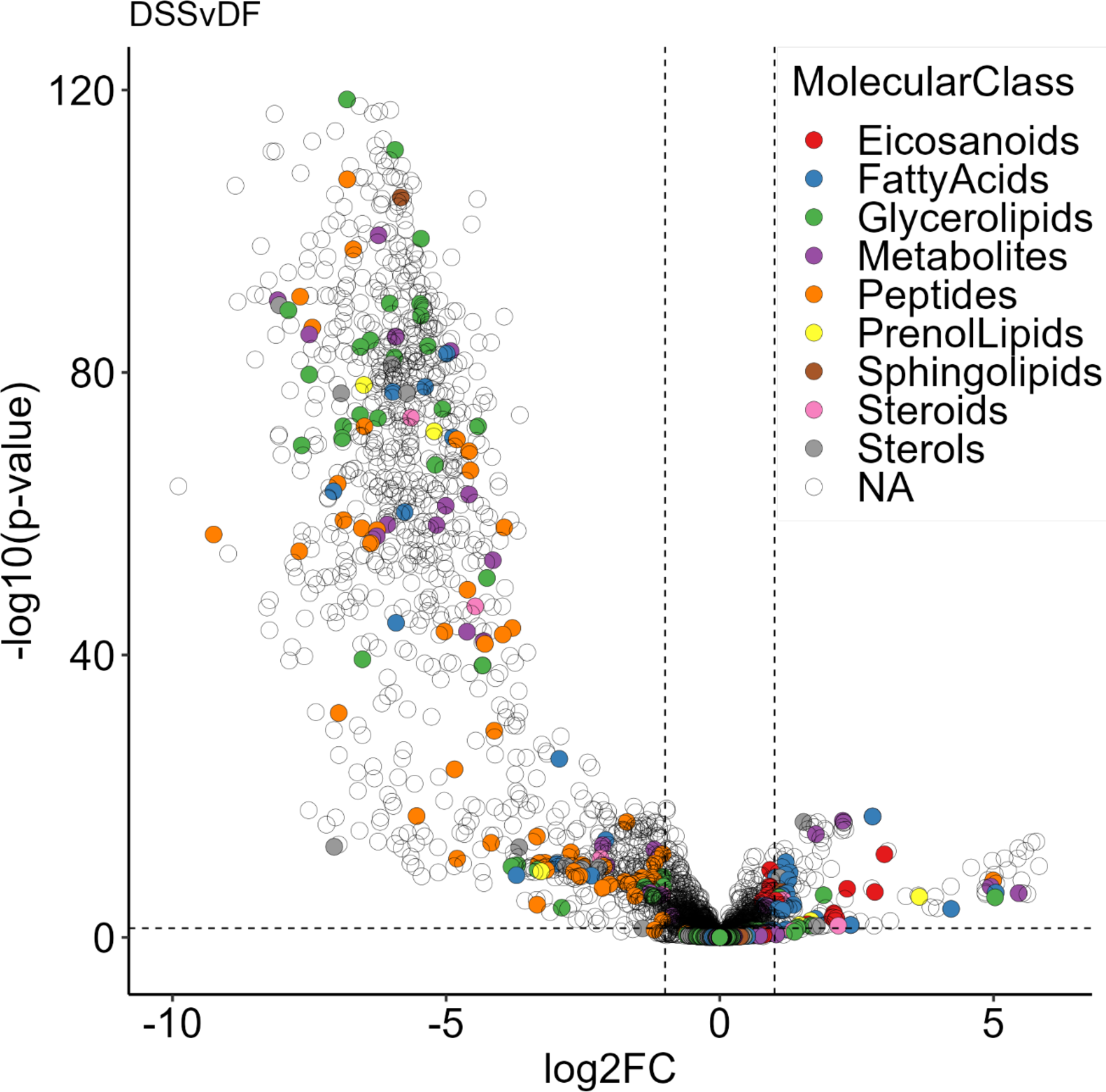
Volcano plot displaying differential abundance of features between DF and DSS. Negative log2FC values can be interpreted as either a lower feature abundance in DSS or greater abundance in DF. Positive log2FC values can be interpreted as either a greater feature abundance in DSS or lower abundance in DF. Colored circles represent the molecular classes of 266 tentatively identified molecules. Noncolored circles represent features that were either manually defined as not biologically relevant, or where tentative identification based on accurate mass was unsuccessful. The horizontal dotted line represents an adjusted p-value of 0.05 and the vertical dotted lines represent log2FC = ± 1.

A subset analysis for days 3 to 6 of illness was done to ensure that day of illness was not a confounding variable for the selected metabolites that were different between DF and DHF/DSS. The subset analysis did not reveal any difference for the selected metabolites relative to the analysis performed for days 1 to 6 of illness. Thus, despite the statistically different distribution for day of illness between DF and DHF/DSS patients, differences in metabolite abundance were due to disease severity and not day of illness.

Serum creatinine level was used as a quality control in this study. Serum creatinine is well-known to positively correlate with increasing age(*22, 23*), and this trend was recapitulated in the current dataset. Creatinine was identified in the current data set and a linear regression on creatinine abundance as a function of patient age produced a positive slope, independent of disease state and sex (Figure S4). Other metabolites identified between disease states did not correlate with age.

### Serum biomarker panel development and disease state classification

Capacity for disease state differentiation and classification using the serum metabolic profile was tested. The 1512 significant features possess information that can differentiate disease state but are not clinically useful without being identified. Therefore, to develop a tool with potential clinical utility, a subset of molecules was identified at confidence levels 1 and 2 using LC-tandem MS (MS/MS). These molecules were chosen based on pathway analysis of the metabolomics dataset (see below), stark trends between disease states, known relevance to dengue virus and disease, previously published literature as well as for quality control (*18*). Thirty-two metabolites were included in the biomarker panel and 12, 19 and 1 metabolites were identified at confidence levels 1, 2 or 3, respectively (*24*). Identification details and relevant pathways for all metabolites identified in this study are summarized in Table S2, and Table S3 summarizes the log2FC and p-values for the pairwise disease state comparisons of these metabolites. LC-MS/MS data validating metabolite identities at confidence level 1 are provided in Figure S3a-n.

Classification models were trained using 32 of the identified metabolites within the biomarker panel (Figure 2, Table S2 and S3). Performance of each model for either differentiating DF from DHF/DSS or ND from DF was tested, and each classification model ranked the 32 metabolites on their importance for the quality of disease state differentiation. The resulting metabolite importance plots for each model are displayed in Figure S5. PCA was employed using only the highest 4 ranked metabolites from rf and adaBoost models (Figure 4a-b). In Figure 4a, DSS samples are almost completely differentiated from DHF samples because serotonin was 1 of the top 4 ranked metabolites in the rf model (Figure S5). The adaBoost model did not rank serotonin in the top 4 metabolites, thus there is greater overlap of DHF and DSS in Figure 4b. Serotonin has great differentiating power for DFvsDSS but is not as great for DFvsDHF or the DFvsDHF/DSS combination. See the section on tryptophan metabolite for details on serotonin.

**Figure 4.**
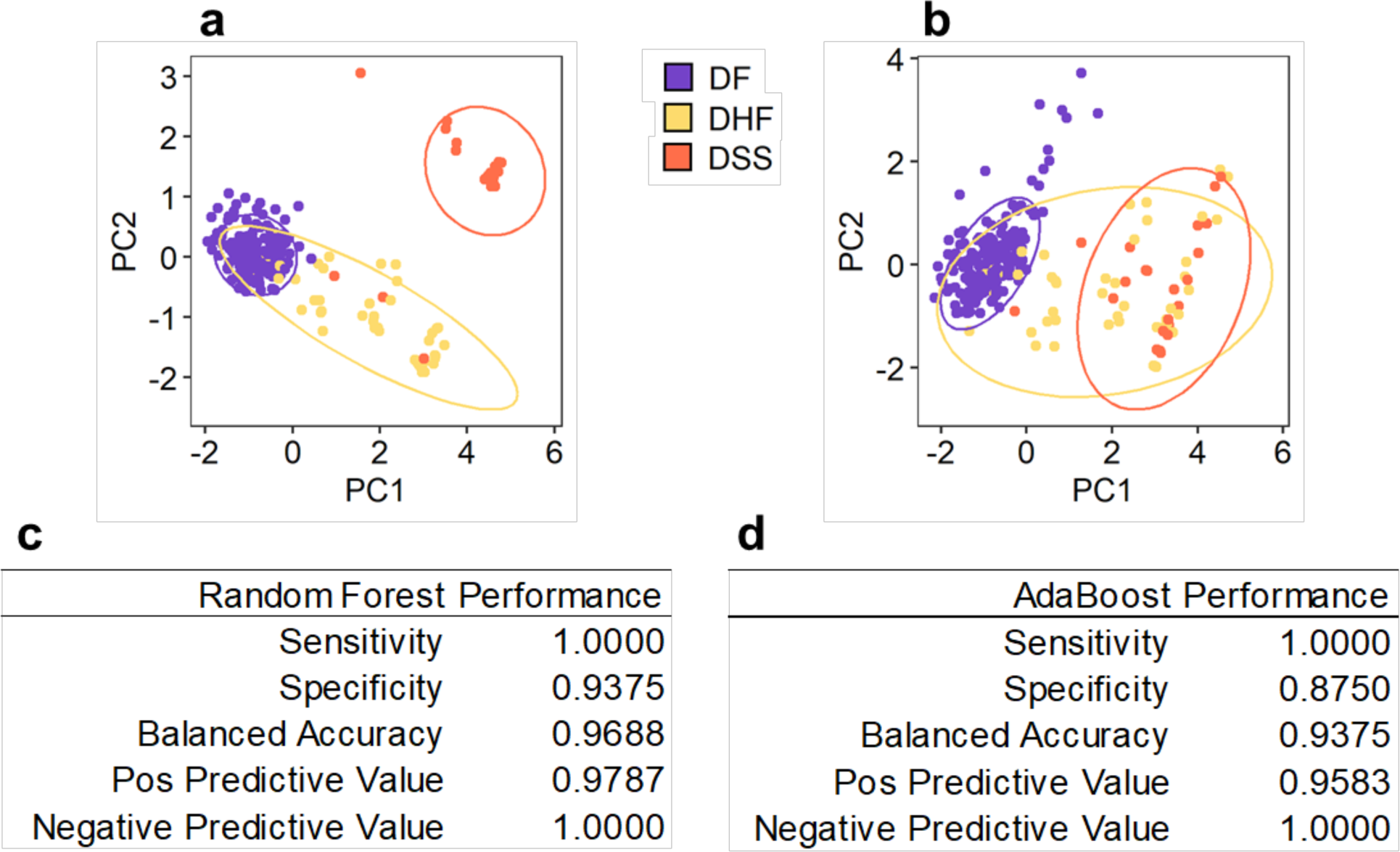
(**a,b**) PCA analyses using the top 4 importance ranked metabolites from the (a) random forest and (c) adaBoost models. (**c,d**) Performance statistics for the (c) random forest and (d) adaBoost models. Sensitivity is the proportion of correctly classified DF samples. Specificity is the proportion of correctly classified DHF/DSS samples. Balanced accuracy is the average of sensitivity and specificity. Positive predictive value is the proportion of samples classified as DF that are truly DF. Negative predictive value is the proportion of samples classified as DHF/DSS that are truly DHF/DSS.

Classification performance for DFvsDHF/DSS was satisfactory for two of three models (Figure 2). The rf and adaBoost models prospered in classification when using all 32 metabolites (Figure 4c-d). When challenged with classification of the 62 samples in the test set, rf misclassified 1 DF sample as DHF/DSS, resulting in a balanced accuracy of 96.9% (Figure 4c). The adaBoost model produced a balanced accuracy of 93.75% with two DF samples misclassified as DHF/DSS (Figure 4d). High classification accuracy and overprediction of severe dengue disease *via* the rf and adaBoost models was a satisfactory outcome. The glm unsatisfactorily misclassified two DHF/DSS cases as DF (underprediction of severe disease).

Classification performance was poorer for the NDvsDF comparison (Figure 2). When challenged with the test sample set, balanced accuracies for the rf and adaBoost models were 70.5 and 72.8%, respectively.

### Metabolic pathway analysis

To glean biological information from this pediatric serum metabolomics dataset, all molecular features (RT, *m/z*, p-value and t-score) were fed into the *mummichog* algorithm to be tentatively identified and mapped to known metabolic pathways. Pathway analysis using all molecular features tentatively proposed 94 enriched metabolic pathways, each pathway having its own magnitude and direction of dysregulation and statistical significance between three pairwise comparisons: DFvsDSS, DFvsDHF and DHFvsDSS. Of the 94 pathways, 86 were common among the three comparisons, 3 were unique to DHFvsDF and 5 were unique to severe disease. Though 86 pathways were identified to be altered in all disease states, the extent of perturbation between comparisons was different. Median t-scores and Fischer’s exact p-values for all pathways and comparisons are shown in Table S4.

Twenty-five perturbed pathways with known biological relevance to dengue and other pathological phenotypes are shown in Figure 5. The 25 pathways are related the metabolism of lipids (e.g., fatty acids, phospholipids, eicosanoids, and sphingolipids), bile acids, amino acids, purines, sugars, and other cellular energy-related small molecules. The TCA cycle, which drives cellular energy generation, was upregulated in DSS, and pathways related to amino acids and purines were downregulated. Dysregulation of lipids, notably upregulation of fatty acid (FA) synthesis, during DENV infection align with our previous *in vitro* and *in vivo* studies(*10, 20*). Metabolism of bile acids and retinoic acid were upregulated in DSS, which may relate to liver damage observed in DHF/DSS(*25, 26*). Regarding sugars, hyaluronan metabolism, sialic acid metabolism and heparan sulfate degradation were all downregulated in DSS, which may relate to the endothelial glycocalyx and its role in viral entry(*27, 28*) or its breakdown in the loss of vascular integrity in severe dengue disease(*29–32*).

**Figure 5.**
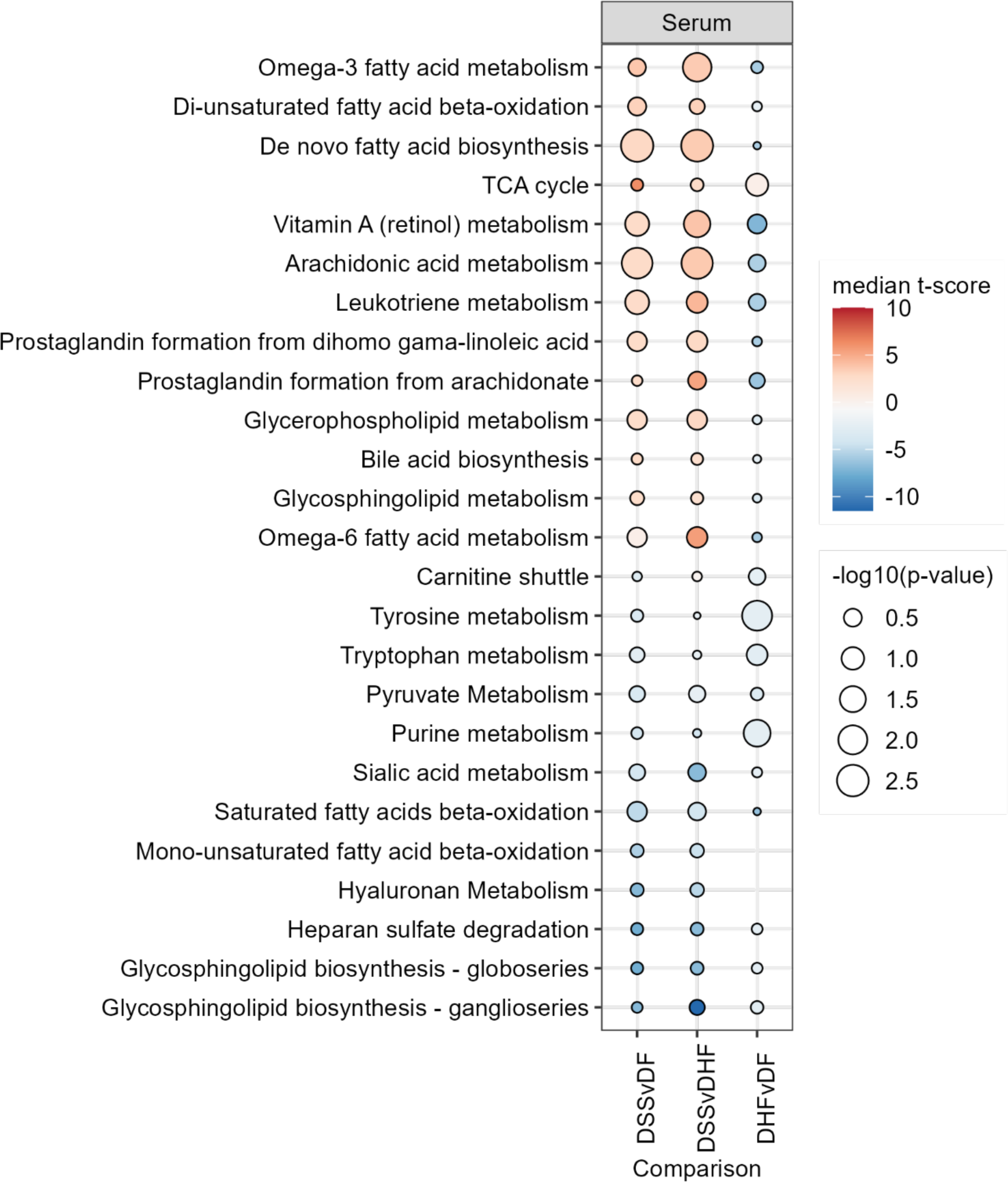
Direction, magnitude, and statistical significance of selected pathway dysregulation upon comparison of DFvsDSS. Red indicates the pathway is upregulated in DSS, and blue indicates the pathway is downregulated in DSS. The direction and magnitude of dysregulation was based on the median t-score of all metabolites that are dysregulated in the indicated pathway. Statistical significance is represented by the −log_10_(p-value) originating from Fischer’s exact test.

### Tryptophan metabolism

Tryptophan metabolism has been implicated in many disease conditions, including viral infection. In this study, tryptophan and seven of its metabolites were identified in patient serum. Metabolite abundance in each patient sample was stratified by disease state and visualized using boxplots, and tryptophan metabolic pathways are illustrated in Figure 6. Additionally, the differential abundance of each metabolite is described by log_2_FC and adjusted p-value from a moderated t-test implemented in the *limma* R package (Table S3). For a subset of 144 patients, sample matched clinical measurements of serum albumin and platelet levels were correlated to tryptophan and serotonin levels, respectively, or correlated to disease state (Figure 7).

**Figure 6.**
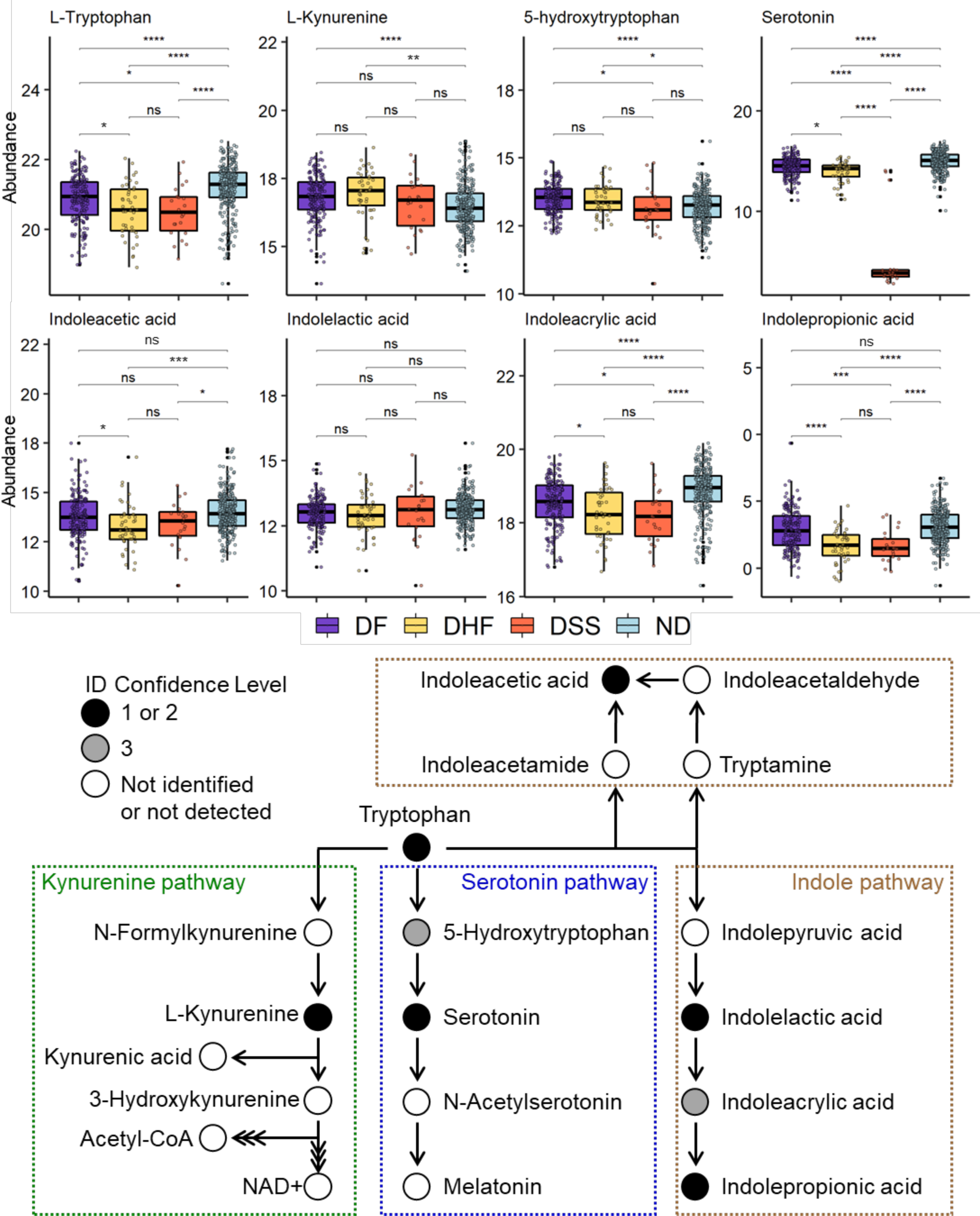
Top) Abundance of tryptophan metabolites in the sera of ND and dengue patients. Abundance is represented as the normalized and log2-transformed LC-MS peak area. Benjamini-Hochberg adjusted p-values generated using a moderated t-test were used to define statistical significance level; p < 0.0001, 0.001, 0.01, 0.05 were each represented by ****, ***, **, or *, respectively. Bottom: Eight metabolites from three tryptophan metabolic pathways were identified in this dataset. Triple arrows between metabolites indicate multiple metabolic steps are required.

**Figure 7.**
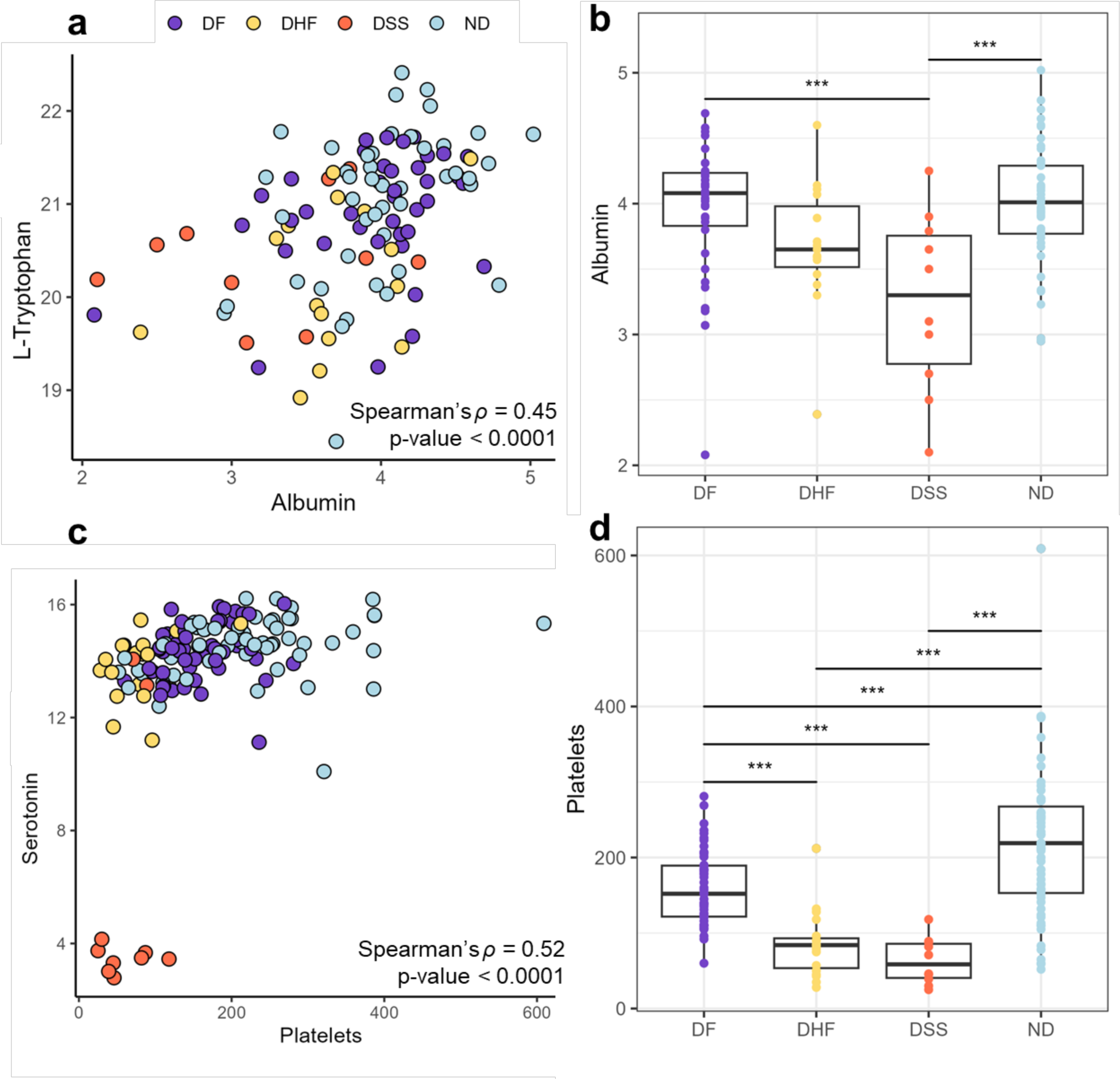
Correlation of serum albumin levels (g/dL) with a) tryptophan serum levels and b) disease state. Correlation of serum platelet levels (platelets/µL) with c) serotonin serum levels and d) disease state. Benjamini-Hochberg adjusted p-values generated using a moderated t-test were used to define statistical significance level; p < 0.0001, 0.001, 0.01, 0.05 were each represented by ****, ***, **, or *, respectively.

Tryptophan abundance was significantly lower in the sera of DF patients compared to ND. Regarding dengue disease severity, tryptophan abundance was increasingly lower in sera of patients with DHF and DSS. Serum tryptophan and albumin levels had a positive Spearman’s ρ correlation coefficient of 0.45 (p-value < 0.0001) (Figure 7a). Additionally, albumin levels in DSS (p-value < 0.0003) and DHF (not significant) were lower on average than in ND or DF patients (Figure 7b).

Kynurenine, a metabolite from one of the two host-driven tryptophan metabolic pathways, was significantly elevated in DF and DHF compared to ND. Kynurenine was not significantly elevated in DSS compared to ND, and there were no observed significant differences in kynurenine between dengue disease states. The first metabolic product in the serotonin pathway, 5-hydroxytryptophan, was significantly elevated in DF and DHF, but not significantly different in DSS, compared to ND. Accordingly, 5-hydroxytryptophan was significantly lower in DSS compared to DF.

The serum abundance of serotonin, the downstream metabolite 5-hydroxytryptophan, was significantly decreased in all dengue disease states. In DF and DHF patients, serotonin abundance was moderately decreased relative to ND. Strikingly, serotonin abundance was massively decreased in DSS patients. Serum serotonin and platelet levels had a positive Spearman correlation coefficient of 0.52 (p-value < 0.0001) (Figure 7c). Additionally, platelet levels were significantly decreased in all dengue disease states compared to ND, but platelet levels were not significantly different between DHF and DSS patients (Figure 7d).

A trend in serotonin abundance was observed when stratified by day of illness, and a similar trend was observed for platelets for a subset of patients (Figure S6). In DF patients, serotonin abundance began ‘normal’ or relatively high on day 1 of illness, then decreased until days 5 or 6, where the rate of decrease (slope) levels off. In DHF, serum serotonin abundance does not appear to reach a minimum at day 6 (Figure S6), although only two DHF samples were collected on day 6 of illness. In DSS patients, serotonin levels were massively decreased on all days (days 3 through 6 measured for DSS). The trend in ND patients was like that in DF patients, potentially indicating similarities between the non-severe febrile DF and other undifferentiated non-dengue febrile illnesses.

Four metabolic products from the indole metabolic pathway of tryptophan via the gut microbiota were measured: indoleacetic acid (IAA), indolelactic acid (ILA), indoleacrylic acid (IAcrA) and indolepropionic acid (IPA). The serum abundances of IAA and IPA were significantly decreased in DHF/DSS, and the abundance of IAcrA was significantly decreased in all dengue disease states (Figure 6, Table S3). ILA abundance was not observed to differ significantly among disease states.

### Omega-3 and omega-6 fatty acid metabolism

The serum abundances of omega-3 (n-3) and omega-6 (n-6) fatty acids (FA) (Figure 8, Table S3) and their downstream bioactive lipids were assessed (Figure S7 and Table S3). Linoleic acid (18:2 n-6), a dietary FA, and three downstream FAs within the desaturation and elongation pathway were identified. Three bioactive eicosanoids within different pathways of the n-6 arachidonic acid cascade were tentatively identified. Regarding the n-3 FAs, α-linolenic acid, and docosahexaenoic acid were identified. Three other n-3 FAs downstream of α-linolenic acid were tentatively identified. All the lipids assessed within n-6 and n-3 FA metabolism were observed to be more abundant in the serum of DSS patients when compared to all other disease states.

**Figure 8.**
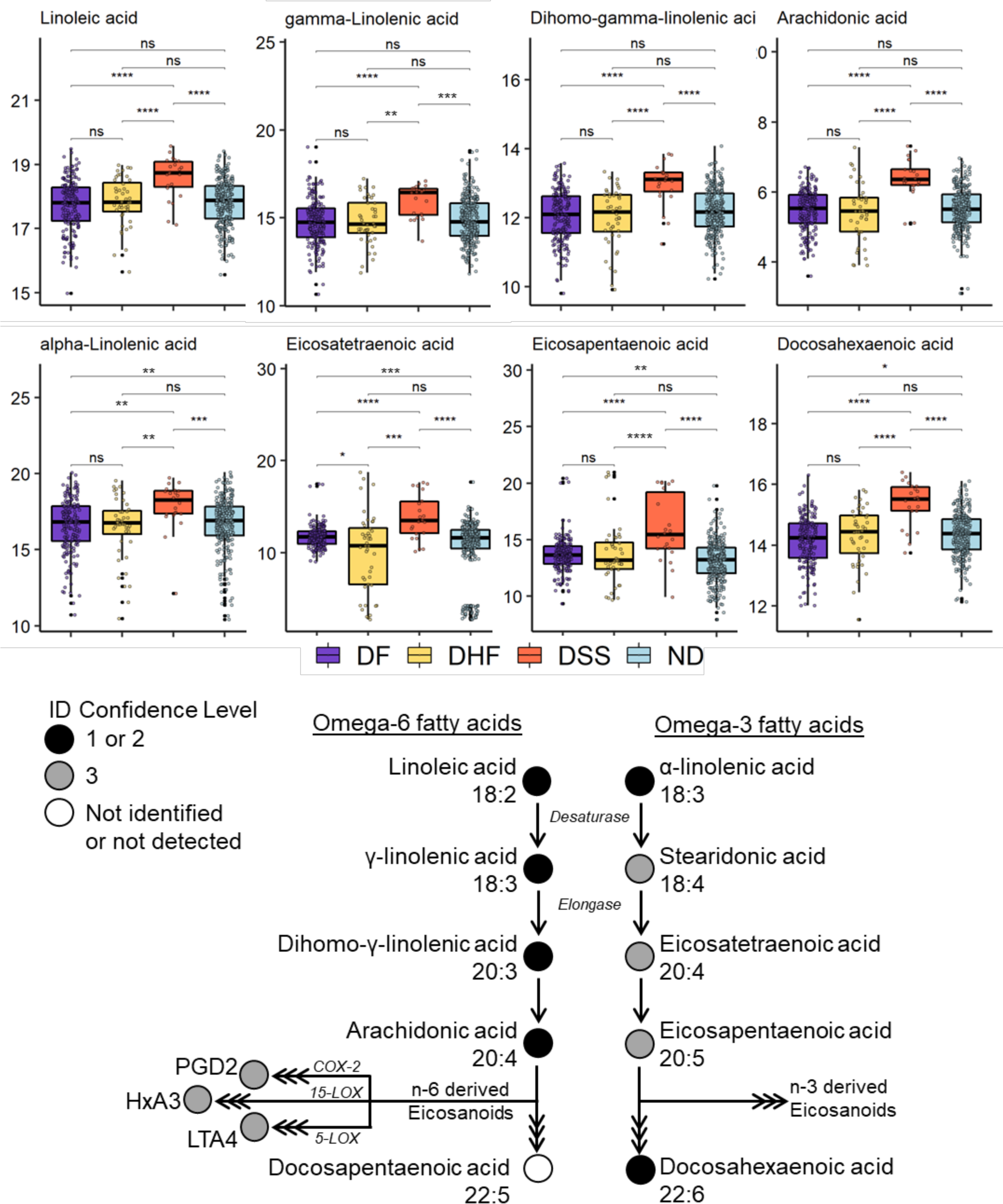
Top) Abundance of n-6 and n-3 FAs in sera of ND and dengue patients. Abundance is represented as the normalized and log2-transformed LC-MS peak area. Benjamini-Hochberg adjusted p-values generated using a moderated t-test were used to define statistical significance level; p < 0.0001, 0.001, 0.01, 0.05 were each represented by ****, ***, **, or *, respectively. Bottom: Fatty acids identified in patient serum. Triple arrows between metabolites indicate multiple metabolic steps are required.

### Sphingolipids

Four interesting sphingolipids were observed in the patient serum samples: sphingosine (d18:1), sphingosine-1-phosphate, sphinganine (d18:0) and sphinganine-1-phosphate (Figure S7). Elevated levels of sphingosine (d18:1) and sphinganine (d18:0) were measured in the sera of DSS patients compared to all other disease conditions. Decreased levels of sphingosine-1-phosphate and sphinganine-1-phosphate were measured in the sera of DHF and DSS patients in a severity dependent manner.

### Other metabolites of interest

Related to purine metabolism, hypoxanthine and inosine were significantly lower in abundance in severe disease (Figure S7). Amino acids, dipeptides, carnitines, and others are visualized in Figure S8. Glycerophospholipids and glycerolipids are visualized in Figure S9. Other identified metabolites of interest are described in Tables S1 and S2.

## DISCUSSION

### Serum metabolite biosignature development

To characterize the serum metabolic changes that correlate with disease state, serum metabolites were measured using untargeted LC-MS. We observed trends in this pediatric serum metabolomics dataset that matched well-established trends in the field. For instance, creatinine levels in serum or urine are commonly used as a measure of renal health. Kidneys filter creatinine out of the blood, such that an increase in serum creatinine levels may be a sign of liver dysfunction. It is also well-known that creatinine levels in serum increase as a function of age in children(*23*) and adults(*22*), independent of sex. In our dataset, the positive correlation between creatinine and patient age recapitulated these well-established trends (Figure S4).

A disparity in the number of unchanged features was observed between disease state comparisons. More unchanged features for DFvsDHF than DFvsDSS indicated that more serum metabolic changes were measured in DSS. Regarding the direction of differential abundance, most features were less abundant in more severe forms of dengue disease, especially DSS. The direction of change in disease states may have implications as correlates of protection (higher in DF, lower in DSS) or as correlates of severe disease pathogenesis (lower in DF, higher in DSS). Thus, interesting features were structurally confirmed for use as biomarkers and so that inferences may be made about their biological roles.

### Disease state classification

Multiple metabolomics studies involving DENV-infected patients have been conducted, all of which have contributed potential biomarkers from diverse sample backgrounds(*10–14*). This study complements the search for distinguishing biomarkers of severe dengue disease, especially due to the large 535 sample set that included 22 DSS patients. Thirty-two of the identified molecules were used as a serum biomarker panel to classify disease state of the pediatric patient samples.

Classification success for DFvsDHF/DSS implied that phenotype differences were reflected in the measured human serum metabolome. The 1 or 2 misclassified samples were due to overpredicting severe disease, a preferred outcome. In this scenario, the misclassified DF patient could be precautionarily monitored for signs of severe disease. Furthermore, poor classification performance for NDvsDF was likely due to fewer phenotype differences between self-limiting, febrile DF and other febrile illnesses (ND). Classification using the 32-metabolite biomarker panel revealed potential for prediction of severe dengue disease, as well as supported the hypothesis that the host serum metabolome contains disease phenotype information. The biomarker panel presented here should be further validated in diverse sample sets representing different geographic locations, genetic backgrounds, and age ranges.

### Tryptophan metabolism

Tryptophan can be metabolized into bioactive molecules that function in inflammation and ageing, gut-brain homeostasis, immune regulation, cardiovascular diseases, and endothelial dysfunction(*33–39*). It is also implicated in infectious diseases(*40, 41*), including dengue virus infection and disease(*10–12, 42*). Importantly, dysregulation of tryptophan metabolism observed within our dataset is consistent with, and contributes to, the knowledge gained from previous findings(*12*).

Hydroxylation of tryptophan via tryptophan hydroxylase (TPH) leads to the formation of bioactive serotonin and melatonin. Serotonin, and its precursor 5-hydroxytryptophan were identified in this study. Serotonin and platelet levels were highest in ND patients, lower in DF patients and drastically depleted with increasing dengue disease severity.

Additionally, serotonin serum abundance in DF patients was observed to trend with day of illness (Figure S6), which matched the trend in platelet counts as non-severe dengue disease progressed. Platelet counts were previously measured from day 1 to 10 by Yasuda *et al*.(*43*), and were observed to hit a low on day 6, then increase and return to normal by day 10 of illness. A similar platelet trend for day 1 to 6 was observed in this study. Because serotonin abundance in DF trended similarly to platelets from day 1 through 6, it is hypothesized that serotonin abundance would also recover to a ‘normal’ level by day 10 in DF patients. Testing this hypothesis would require collection and analysis of patient samples up to at least day 10 of illness.

Serotonin can be either protective or pathogenic in mediation of shock, and vascular endothelial dysfunction, and leak(*44, 45*). In a pathogenic role, serotonin has been shown to induce local vasodilation when released at the vascular endothelium, affecting endothelial function, and contributing to vascular leakage and shock. Systemic shock can be induced by circulating antibody-antigen immune complexes via the platelet Fcγ receptor RIIA(*45*) and serotonin, within this immune complex-induced mechanism, is pathogenic and is required for vasodilation. Serotonin can also be derived from mast cells (MC) and induce significant platelet activation, leading to thrombocytopenia via platelet aggregation and increased splenic uptake followed by phagocytosis(*46*). However, platelet activation is also necessary for hemostasis and vascular wall maintenance and thus serotonin can also play a protective role.

The reason for circulating serotonin depletion in dengue disease is still unclear. It could be hypothesized that thrombocytopenia is a cause of decreased serotonin. However, this current study revealed a massive decrease in serotonin, and not platelets, in DSS relative to DHF suggesting other, or additional, mechanisms of serotonin depletion from serum (Figure 7). One potential serotonin sink in dengue disease could be the liver. Liver damage is a clinical presentation of severe dengue disease, indicated by increased circulation of liver transaminases and bilirubin. Multiple studies involving DENV infected adults and children reported that greater than 90% of patients had elevated circulating liver aminotransferases(*47*). Through altered expression of serotonin receptor subtypes in the liver (e.g., 5-HT_2A_ and 5-HT_2B_ serotonin receptors), serotonin mediates hepatocyte proliferation and restoration of hepatic mass upon injury(*48–51*). Therefore, the uptake and usage of serotonin by hepatocytes to counteract dengue-induced liver damage may, in part, account for decreased serum serotonin levels.

5-Hydroxytryptophan is the direct precursor of serotonin. The elevated serum 5-hydroxytryptophan levels in DF and DHF compared to ND patients could be a protective metabolic response, in that host metabolism shifts to replenish serotonin levels. In DSS patients, 5-hydroxtryptophan serum levels are not significantly different from ND patients and are significantly lower than DF patients. The lack of elevated 5-hydroxtryptophan levels in DSS may indicate that patient metabolism did not effectively shift to supplement depleted serotonin levels. It is also important to note that 5-hydroxytryptophan did not show the same decrease in DSS that was observed for serotonin. Therefore, as hypothesized in a previous study(*12*), normal 5-hydroxytryptophan levels in DSS patients indicate that serotonin synthesis is not the major perturbation, rather serotonin release or uptake is highly perturbed.

Decreased production of albumin by the liver (e.g., liver damage) or increased escape of circulating albumin from the vascular space (increased vascular permeability) can lead to hypoalbuminemia(*52*). Two studies involving DENV infected children reported that 60-80% of children present with hypoalbuminemia(*47*). In blood, most tryptophan (90%) is bound to albumin, and 10% of tryptophan is unbound (free) and available for tissue uptake. Therefore, decreased tryptophan levels detected in the sera of dengue patients may be related to decreased albumin, which was supported by the positive correlation observed between the two measures.

Also, rapid equilibration between albumin-bound and free tryptophan in blood paired with sustained tissue uptake results in depleted tryptophan blood levels(*53*). Tissue uptake and subsequent usage of tryptophan through one of its four metabolic pathways, without sufficient replenishment of tryptophan, could be another reason for decreased serum tryptophan in dengue disease.

IPA was reported to have immunomodulatory, anti-inflammatory and antioxidant effects, and various protective functions in mammals(*37, 54*). IPA abundance has a positive correlation with factors that promote cardiovascular health, and IPA levels decrease in scenarios where cardiovascular health is diminished(*55*). Decreased IPA levels in DHF and DSS patients could be related to diminished cardiovascular health.

### Omega-3 and omega-6 fatty acid metabolism, and eicosanoids

Lipid metabolism is dysregulated in all disease states in our dataset; specifically, FA metabolism and biosynthesis, as well as FA beta-oxidation. FAs and other cellular lipids are important structural, signaling and energy-yielding molecules for viral entry, replication, assembly, and release. Previous work has shown that fatty acid biosynthesis is actively dysregulated during viral infection to benefit assembly and function of viral replication factories(*56–59*). FA oxidation pathways and the citric acid cycle (TCA) were observed to be altered across disease states. DENV utilizes FA beta-oxidation to fuel the mitochondrial TCA cycle, which in turn provides adenosine-5’-triphosphate (ATP) as energy for the viral lifecycle. Cellular lipids can also mediate the inflammatory response associated with disease. Increased linolenic acid (n-3) and linoleic acid (n-6, precursor to arachidonic acid) metabolism lead to formation of the pro- and anti-inflammatory and bioactive lipid mediators, such as eicosanoids (leukotrienes, prostaglandins, thromboxanes)(*60–62*).

Linoleic acid (18:2 n-6) is an essential n-6 FA that is metabolized to arachidonic acid or other very-long chain FAs. Enhanced levels of linoleic acid and metabolites serve to increase pools of arachidonic acid and its downstream bioactive lipids. Three eicosanoids were tentatively identified in this study and were elevated in the sera of DSS patients (Figure S7): leukotriene A_4_ (LTA_4_), prostaglandin E_2_ (PGE_2_), and hepoxilin A_3_ (HxA_3_). These three molecules may have implications for increased vascular permeability observed in severe dengue disease.

Leukotrienes are produced by the enzyme 5-lipoxygenase (5-LOX). Initially, 5-LOX generates 5-hydroperoxyeicosatatraenoic acid, which is unstable and is rapidly converted to 5-hydroxyeicosatetraenoic acid or to LTA_4_, which is converted to LTB_4_ or LTC_4_. LTC_4_ serves to increase vascular permeability and plasma leakage (*63, 64*), both of which are symptoms observed in DHF and DSS.

Prostaglandins are produced via arachidonic acid through cyclooxygenase (COX) isoenzymes. PGE_2_ is significantly increased during inflammation where it causes increased microvascular permeability(*61, 65*). Another study demonstrated involvement of COX-2 in DENV replication in cell culture(*66*), potentially implying a virally induced mechanism for increased PGE_2_. HxA_3_ is a non-canonical eicosanoid that is produced when arachidonic acid is converted to 12S-HpETE via the 12S-LOX enzyme, which is then converted to HxA_3_. HxA_3_ induces neutrophil chemotaxis, stimulates release of arachidonic acid and diacylglycerol and increases vascular permeability in rat skin(*67, 68*). Additionally, elevated circulating arachidonic acid levels could, in part, be related to HxA_3_ activity.

### Sphingolipids

Sphingosine is the common backbone of the diverse class of sphingolipid molecules, which can have signaling and structural roles. Sphingosine, or the closely related sphinganine, can be phosphorylated to generate a potent signaling molecule. Based on ELISA measurements of dengue patient sera from Colombo, Sri Lanka, sphingosine-1-phosphate decreased in a severity dependent manner and when compared to healthy controls(*69*). This current mass spectrometry-based study of patients from Nicaragua recapitulated the disease severity dependent decrease in serum sphingosine-1-phosphate, and revealed this trend extends to DSS patients. Like serotonin, sphingosine-1-phosphate is stored in and released by platelets(*70*), thus these trends may be associated with thrombocytopenia.

Sphingosine-1-phosphate and its various receptors (S1PR1 through S1PR5) have been implicated in both protection and disruption of the endothelial barrier. Upon binding of sphingosine-1-phosphate, S1PR1 promotes regulation of endothelial cell (EC) function via downstream signaling, and, contrarily, S1PR2 induces vascular permeability via the Rho-ROCK-PTEN signaling cascade that facilitates phosphorylation and loss of VE-Cadherin at adherens junctions(*71–73*). Modak *et al.* hypothesized that low serum sphingosine-1-phosphate and DENV-induced upregulation of the high-affinity S1PR2 in ECs results in preferential activation of disruptive signaling pathways that lead to vascular permeability(*73*).

Sphinganine-1-phosphate was progressively decreased in the serum of DF, DHF and DSS patients in this study. With its fully saturated alkyl chain, sphinganine-1-phosphate lacks a double bond, thus, is closely related to sphingosine-1-phosphate and can bind S1P receptors. Sphinganine-1-phosphate was reportedly depleted in mice after hepatic ischemia reperfusion, and exogenous replenishment of sphinganine-1-phosphate protected the mice against liver and kidney injury and improved EC integrity and vascular function(*74*). The protective effect of sphinganine-1-phosphate on EC was found to be related to S1PR1(*75*). Thus, depleted sphinganine-1-phosphate in the sera of DENV infected patients may be related to vascular integrity.

### Inosine and hypoxanthine

Decreased levels of inosine and its downstream product hypoxanthine (Figure S7) in dengue disease, especially severe disease, could be related to higher levels of adenosine (inosine precursor) and lower levels of xanthine and uric acid (hypoxanthine catabolites). Adenosine is a vasodilator and platelet aggregation inhibitor(*76–79*). Circulating adenosine was reported to inhibit polymorphonuclear leukocyte function, resulting in decreased synthesis of specialized pro-resolving lipid mediators during coagulation that drive resolution of inflammation(*80*). Additionally, circulating adenosine deaminase abundance and activity (the enzyme that converts adenosine to inosine) are reportedly altered in various diseases(*81–83*).

## STUDY LIMITATIONS

Biomarkers with potential for triaging severe disease must be perturbed and measurable within the first 3 days of the acute phase of dengue disease. This study includes samples that were collected from day 0 to 6 of illness, thus, the perturbed metabolites described here are only associated with DHF/DSS and cannot be used to predict progression to DHF/DSS. The next step towards biomarkers for early triage will require analysis of the biomarkers presented here within samples from patients on day 3 of illness or earlier, who then later progress to severe disease. Additionally, further expansion of these studies to other geographical regions, genetic backgrounds, age ranges and longitudinal sample collections will help to fortify such biomarker panels.

This study did not include healthy controls, but using the ND control and various dengue disease severities this study was able to assess the serum metabolic changes specific to DF febrile illness, and metabolic changes between DF and DHF/DSS.

Serum sample preparation involves platelet coagulation, which should be considered when interpreting the relationship between platelets and serum metabolites. Additionally, coagulation can induce production of arachidonic acid-derived lipid mediators. But the metabolic associations between disease states remain valid because this study included only serum samples, but expansion to other biofluids (e.g., plasma that is not coagulated upon collection) may be useful.

Like all metabolomics studies, this study had an inherent measurement bias based on molecular polarity. The extraction protocol and LC-MS/MS analyses were optimized for mid-polar metabolites and lipids, so additional workflows would be necessary if comprehensive analyses of highly polar metabolites, sugars, or nonpolar lipids were desired.

## CONCLUSIONS

In this study we demonstrate that the human serum metabolome is dynamically altered in response to DENV infection and is associated with disease severity. The serum metabolome was used for successful disease state classification and to explore the biochemistry of severe dengue disease pathogenesis. Unique to this study, was the identification of the metabolic biosignatures of DSS. Biomarkers of DHF were identified that were either novel or that recapitulated those reported in studies from other geographical regions, supporting the hypothesis that the metabolome could provide information that is independent of genetic backgrounds. These studies could be critical in assembling biomarkers for early triaging of severe dengue disease.

## MATERIALS AND METHODS

### Study Design

For this study 535 serum samples from 535 individuals were retrospectively obtained from two different well-established studies in Managua, Nicaragua. A set of 122 well-characterized samples collected between 2012 and 2013 were sourced from the Pediatric Dengue Cohort Study (PDCS), following over 3,800 children between the ages of 2 and 14 years old since 2004 (Figure 1A). Cohort patients were enrolled as healthy volunteers, followed for all medical episodes, and monitored for suspected arboviral diseases; those who met the case definition of dengue, or undifferentiated febrile illnesses, were worked up for laboratory confirmation using molecular biological, virological, and/or serological methods. Another 413 serum samples were obtained from patients 1 to 16 years old (Figure 1A) in the Pediatric Dengue Hospital-based Study (PDHS) who presented at the Hospital Infantil Manuel de Jesús Rivera, the National Pediatric Reference Hospital in Nicaragua, between 2005 and 2015 with a fever or history of fever for <7 days and one or more of the following signs and symptoms: headache, arthralgia, myalgia, retro-orbital pain, positive tourniquet test, petechiae, or signs of bleeding. Cases were laboratory-confirmed for DENV infection by detection of DENV RNA by RT-PCR, isolation of DENV, seroconversion of DENV-specific IgM antibody titers observed by MAC-ELISA in paired acute- and convalescent-phase samples, and/or a ≥4-fold increase in anti-DENV antibody titer measured using inhibition ELISA in paired acute and convalescent samples(*84–86*). Immune status was determined using Inhibition ELISA in early convalescent samples (14 or more days post-onset of symptoms; <2,560 was considered primary infection and ≥2,560 was considered secondary infection(*85, 87*). Disease severity (DF, DHF, or DSS) was diagnosed by computerized algorithms based on the 1997 WHO schema were used to classify cases(*21*). Samples that were negative for DENV infection were classified as ND. Metadata recorded included patient sex, age, disease outcome (disease state), infection history, DENV serotype, day of illness (number of days of fever at time of sample collection) and date of sample collection, as well as detailed clinical data across disease evolution.

### Statistical analyses

Univariate analysis of features was implemented in the R package *limma* version 3.32.10, in which linear models with empirical Bayes statistics were applied feature-wise to generate pairwise comparison of feature abundances across disease states(*88–90*). The models provided estimated log2 FCs, moderated t-statistics, and false discovery rate adjusted p-values for each feature. Significant features were defined by log_2_FC ≥ 1 and p-value < 0.05 after adjustment for false discovery rate (*91*).

To develop classification models, the 535 samples were randomly divided within disease states into a training set (75% of samples within each disease state) and a test set (remaining 25% of samples), where disease states were ND, DF and severe disease (DHF/DSS). DHF and DSS were combined into the severe disease category to alleviate the small sample size for each disease state. Training sets were used to fit models. Test sets were used only to test predictions from fitted models.

Using the training sample set, classification models (random forest, adaBoost, generalized linear model) were constructed for two separate classification problems: NDvsDF and DFvsDHF/DSS(*92–94*). The R “caret” package verison 6.0.93 was used to build several classification models(*95*), also including support vector machines, linear and quadratic discriminate analysis, and k-nearest-neighbors. Parameters were chosen by 5-fold cross-validation; peak areas were scaled and centered. The generalized linear model, random forest, and adaBoost.M1 gave perfect classification on the training set and were therefore chosen for evaluation with the test set.

## LIST OF SUPPLEMENTARY MATERIALS

Materials and Methods

Chemicals and Reagents
Sample Preparation and Extraction
Untargeted liquid chromatography-mass spectrometry
Metabolite identification via liquid chromatography-tandem mass spectrometry
Data Processing
Metabolic pathway analysis
Table S1. Summarized patient metadata and statistical assessment
Figure S1a-b. Volcano plots displaying metabolite differential abundance between disease states
Figure S2. Frequency distribution of log_2_FC values for all LC-MS features
Table S2. Metabolites of interest LC-MS metrics
Table S3. Metabolites of interest log_2_FC and adjusted p-values
Figure S3a-n. LC-MS/MS validation of metabolite identities at confidence level 1
Figure S4. Creatinine abundance correlations with patient age
Figure S5. Metabolite importance plots for the random forest and adaBoost classification models
Figure S6. Serotonin and platelet trends as a function of patient day of illness
Figure S7. Metabolite abundance boxplots in each disease state for eicosanoids, purines and sphingolipids
Figure S8. Metabolite abundance boxplots in each disease state for amino acids, dipeptides, carnitines, and other metabolites.
Figure S9. Metabolite and abundance boxplots in each disease state for glycerophospholipids and glycerolipids
Table S4. Median t-scores and Fischer’s exact test p-values for all metabolic pathways and disease state comparisons

