## Supplemental file for "Perturbed pediatric serum metabolome in mild and severe dengue disease"

**1** Department of Microbiology, Immunology, and Pathology, Colorado State University, Fort Collins, Colorado, United States of America, **2** Sustainable Sciences Institute, Managua, Nicaragua, **3** Laboratorio Nacional de Virología, Centro Nacional de Diagnóstico y Referencia, Ministry of Health, Managua, Nicaragua, **4** Division of Infectious Diseases and Vaccinology, School of Public Health, University of California, Berkeley, Berkeley, California, United States of America

### LIST OF SUPPLEMENTARY MATERIALS

Materials and Methods

- Chemicals and Reagents
- Sample Preparation and Extraction
- Untargeted liquid chromatography-mass spectrometry
- Metabolite identification via liquid chromatography-tandem mass spectrometry
- Data Processing
- Metabolic pathway analysis

Table S1. Summarized patient metadata and statistical assessment

Figure S1a-b. Volcano plots displaying metabolite differential abundance between disease states

Figure S2. Frequency distribution of $log_2$FC values for all LC-MS features

Table S2. Metabolites of interest LC-MS metrics

Table S3. Metabolites of interest $log_2$FC and adjusted p-values

Figure S3a-n. LC-MS/MS validation of metabolite identities at confidence level 1

Figure S4. Creatinine abundance correlations with patient age

Figure S5. Metabolite importance plots for the random forest and adaBoost classification models

Figure S6. Serotonin and platelet trends as a function of patient days of illness

Figure S7. Metabolite abundance boxplots for eicosanoids, purines and sphingolipids

Figure S8. Metabolite abundance boxplots for amino acids, dipeptides, carnitines and other metabolites

Figure S9. Metabolite abundance boxplots for glycerophospholipids and glycerolipids

Table S4. Median t-scores and Fischer's exact test p-values for all metabolic pathways and disease state comparisons

### MATERIALS AND METHODS

#### Chemicals and Reagents

LC-MS grade water and methanol were purchased from Honeywell (Charlotte, NC). LC-MS grade acetonitrile was purchased from Fischer Scientific (Hampton, NH). Analytical grade 5-hydroxy-L-tryptophan, arachidonic acid, arachidonic acid-$d_8$, creatinine, creatinine-$d_3$, dihomo-gamma-linolenic acid, dihomo-gamma-linolenic acid-$d_6$, gamma-linolenic acid, indole-3-acetic acid, indole-3-lactic acid, indole-3-propionic acid, linoleic acid, linoleic acid-$d_4$, L-kynurenine, L-tryptophan, serotonin hydrochloride, and serotonin-$d_4$ hydrochloride were purchased from Cayman Chemical (Ann Arbor, MI). Analytical grade carnitine, hypoxanthine and inosine were purchased from Sigma Aldrich (St. Louis, MO).

#### Sample preparation and metabolite extraction

Twenty microliters of each serum sample were aliquoted into individual microcentrifuge tubes, and an additional 20 µL of each serum sample were combined to generate the pooled quality control (QC). For metabolite extraction, patient serum and pooled QC samples were randomized. Five microliters of L-tryptophan-$d_5$ (80 ng/mL) heavy-isotope labeled internal standard was added to 20 µL of patient serum (or pooled QC aliquot) in a microcentrifuge tube. To precipitate proteins, 100 µL of cold methanol was added to the serum and samples were incubated for 12 hours at -80 °C. Samples were then centrifuged at 4 °C for 15 min at 18,000 *x g* to pellet proteins. Supernatant was then transferred to a new microfuge tube and dried under nitrogen. Samples were reconstituted in 25 µL of methanol/water (50/50), let stand at room temperature for 15 minutes, vortexed for 20 seconds and centrifuged to pellet insoluble debris. Sample supernatants were then transferred to autosampler vials fitted with low-volume inserts and immediately submitted for LC-MS analysis. Serum samples were prepared and analyzed in 6 randomized batches. A pooled quality control sample was run after every 5 experimental samples and a solvent blank was run after every 10 samples.

**Untargeted liquid chromatography-mass spectrometry**

Sample order for injection was randomized. Each sample was injected (7.5 μL) onto an Agilent 1290 HPLC system where metabolites were separated on an XBridge BEH C18 column (2.5 μm particle size, 2.1x100mm, Waters Millford, MA, USA). The total mobile phase flow rate was 0.250 mL/min, made up of water + 0.1% formic acid (mobile phase A) and 95/5 acetonitrile/water + 0.1% formic acid (mobile phase B). For metabolite separation, the mobile phase composition began at 5% B and held until 0.5 minutes, then from 0.5 to 14 minutes the mobile phase composition was adjusted in a linear fashion to 98% B. From 14.5 to 15 minutes the mobile phase composition was returned to starting gradient conditions of 5% B. From 15 to 19.5 minutes the starting gradient conditions were held to equilibrate the LC column for the subsequent sample injection and metabolite separation. The LC column outlet was coupled to the electrospray ionization (ESI) source of an Agilent 6224 time-of-flight mass spectrometry system. The mass spectrometry system was operated in positive ionization mode. The ESI source was electrically grounded, and the MS-inlet capillary was held at -4000 V to generate positive ions from the metabolites that eluted from the LC column. The ESI nebulizer nitrogen gas was set to 45 psi, and the heated counter flow of nitrogen ('dry gas' used to aid in droplet desolvation) was flowed at 10 liters/minute and held at 310 °C. Beyond the MS-inlet capillary, the fragmentor voltage was set to 120 V to aid in ion desolvation and transmission, and the skimmer voltage was set to 50 V. The peak-to-peak voltage of the ion transfer octopole was set to 750 V. The time-of-flight mass analyzer was set to scan between *m/z* 70 to 1700 and collected full scan (MS1) spectra at a rate of 1.66 spectra/sec.

**Metabolite identification via liquid chromatography-tandem mass spectrometry**

To confirm metabolite identity, RT, *m/z*, and the collision-induced dissociation (CID) product ion spectra were collected for each metabolite. For level 1 identification, RT, *m/z,* and CID product ion spectrum from a pure synthetic standard were compared to that of the molecule originating from serum. For level 2 identification, RT, *m/z* and CID product ion spectrum for the molecule in serum were compared to literature, databases, or known dissociation patterns*(1–3)*. Confirmation of metabolite identities was performed on an Agilent 1290 HPLC system coupled to an Agilent 6546 quadrupole time-of-flight (QTOF) mass spectrometry system. LC conditions, ionization polarity and ion transfer optics in the MS were identical to conditions used for the untargeted LC-MS experiment. To acquire CID product ion spectra for metabolites of interest, the quadrupole mass analyzer was set to pass the precursor *m/z* of interest with an isolation width of 1.3 Da. Precursor ions were transmitted through the quadrupole mass analyzer and accelerated into the collision cell filled with $N_2$ gas (24 psi). Within the collision cell, precursor ions underwent energetic collisions with $N_2$ gas molecules. Product ion spectra were collected at collision energies of 10 and 40 arbitrary units. After the collision cell, precursor and product ions were pulsed into the TOF for mass analysis and subsequent detection.

**Data processing**

*Molecular feature extraction*. The Agilent data files were converted to from .d to .mzML format using ProteoWizard MS Convert version 3.0.6478 64 bit. Peak picking, retention time correction, chromatogram alignment and gap filling were performed using XCMS software version 1.46 in R version 3.2.2 *(4)*. The R package IPO was used to optimize XCMS parameters on the dataset*(5)* and CAMERA was used for deisotoping*(6)*

*Molecular feature description.* Within each data file, the abundance and tentative identity of many potential metabolites (molecular features) are embedded. Two LC-MS metrics that define a molecular feature are the RT of its LC peak and the *m/z* of the ion that correlates to the chromatographic peak. Enabled by the accurate mass measurement (*m/z*) capability of the time-of-flight mass analyzer, each feature was tentatively identified by searching its measured accurate mass (within ± 20 parts-per-million mass error) against the Human Metabolome Database

(HMDB), LipidMaps, Metlin and Kegg databases using CEU mass mediator*(7)*. Additionally, the area under the chromatographic peak of a molecular feature represents its abundance in each serum sample.

*Preprocessing*. All data preprocessing steps were conducted in R version 3.4.2 *(8)*. Features were first filtered to remove any that failed to meet the following criteria: 1) present in at least 20% of all samples across all analysis batches; 2) present in at least 75% of all pooled QC samples; 3) present in at least 70% of samples from at least one disease group (ND, DF, DHF, DSS).

Normalization was conducted stepwise. First, within each batch separately, features that were not present in at least 80% of pooled QC samples in that batch were removed. Features were then normalized using a tobit regression (left-censored at the minimum value for each batch) fitted to the pooled QC samples, implemented in the R package 'AER' version 1.2.5*(9, 10)*. Batches were then combined, retaining only those features present in all batches, and normalized feature-wise over all batches by the ratio of pooled QC mean batch abundance to pooled QC overall mean abundance.

The year of sample collection influenced the presence or abundance of some features, suggesting that length of storage and/or methods of collection and handling may be influencing the results. Therefore, features were removed if found to be present (in ≥50% of all samples, irrespective of group) only in samples collected in the 2005-2009 seasons, or only in samples collected in the 2001-2015 seasons. However, if the feature appeared to be specific to a samples infected with a single serotype (i.e. found in >50% of samples of that serotype, and <50% of samples from other serotype), then the feature was retained. Finally, features that had a coefficient of variation >30% in pooled QC samples after combining and normalizing batches were deemed unreliable and removed. Abundances were $\log_2$-transformed. Missing values were imputed using a random forest algorithm implemented in the R package missForest version 1.4

*(11)*. Abundance variance was calculated for each feature across all samples, and features in the lowest quartile were excluded from analysis *(12)*.

**Metabolic pathway analysis**

Differential abundance and statistics (t-score and p-value) were calculated feature-wise via *limma* for each pairwise comparison of dengue disease severity. Pathway analysis was performed on results from each pairwise disease state comparison. Feature *m/z*, RT, p-value and t-score were used as inputs for the *mummichog* metabolic pathway analysis algorithm*(13)*. For *mummichog* analysis, primary ion types were forced. The metabolite p-value cutoff was 0.05 and a minimum of 3 metabolites were required to flag a pathway. A 20-ppm mass error tolerance was set for tentative metabolite annotation.

**Table S1.** Summarized patient metadata and statistical assessment.

| | Clinical Diagnosis | | | p-value | | |
|---|---|---|---|---|---|---|
| | **ND** | **DF** | **DHF/DSS** | **ND vs. DF** | **ND vs. DHF/DSS** | **DF vs. DHF/DSS** |
| **Number of Patients** | 284 | 185 | 66 | | | |
| **Age (mean (range))** | 8.22 (0.6 to 14.6) | 9.44 (0.5 to 15.9) | 9.71 (0.6 to 15.6) | < 0.0001 | 0.2 | 1 |
| **Sex** | | | | 1 | 1 | 1 |
| Female (%) | 134 (47.2%) | 97 (52.4%) | 26 (39.4%) | | | |
| Male (%) | 150 (52.8%) | 88 (47.6%) | 40 (60.6%) | | | |
| **Days of Illness (mean(range))** | 2.9 (1 to 7) | 3.6 (1 to 6) | 4.3 (1 to 6) | < 0.0001 | | < 0.0001 |
| **DENV Serotype** | | | | | | < 0.0001 |
| DENV1 (%) | | 91 (49.2%) | 11 (16.7%) | | | |
| DENV2 (%) | | 45 (24.3%) | 45 (68.2%) | | | |
| DENV3 (%) | | 23 (12.4%) | 7 (10.6%) | | | |
| Unknown (%) | | 26 (14.1%) | 3 (4.5%) | | | |
| **Infection History** | | | | | | < 0.0001 |
| Primary (%) | | 98 (53.0%) | 12 (18.2%) | | | |
| Secondary (%) | | 84 (45.4%) | 50 (75.8%) | | | |
| Unknown (%) | | 3 (1.6%) | 4 (6.0%) | | | |

Each variable is presented as either number of patients (percentage of patients), or mean value (range of values). Statistical assessment of demographics between disease states; (sex) indicated no statistically significant differences in sex distribution (Chi-squared test, $p = 1$); (age) indicated a significantly different distribution between ND and DF (Welch’s t-test, $p < 0.0001$), but age distribution was not found to be statistically different between DF and DHF/DSS (Welch's t-test, $p = 1$) or ND and DHF/DSS (Welch's t-test, $p = 0.20$), (days of illness) indicated significantly different distributions between ND and DF (Welch’s t-test, $p < 0.0001$) and between DF and DHF/DSS (Welch’s t-test, $p < 0.0001$), see discussion on days of illness subset analysis; (infection history) indicated DHF/DSS has a higher percentage of secondary infections (Fischer Exact Test, $p < 0.0001$); (serotype) indicated DF has a greater percentage of serotype 1 and DHF/DSS has a greater percentage of serotype 2 (Chi-squared test, $p < 0.0001$). Reported p-values were Bonferroni adjusted for multiple comparison.

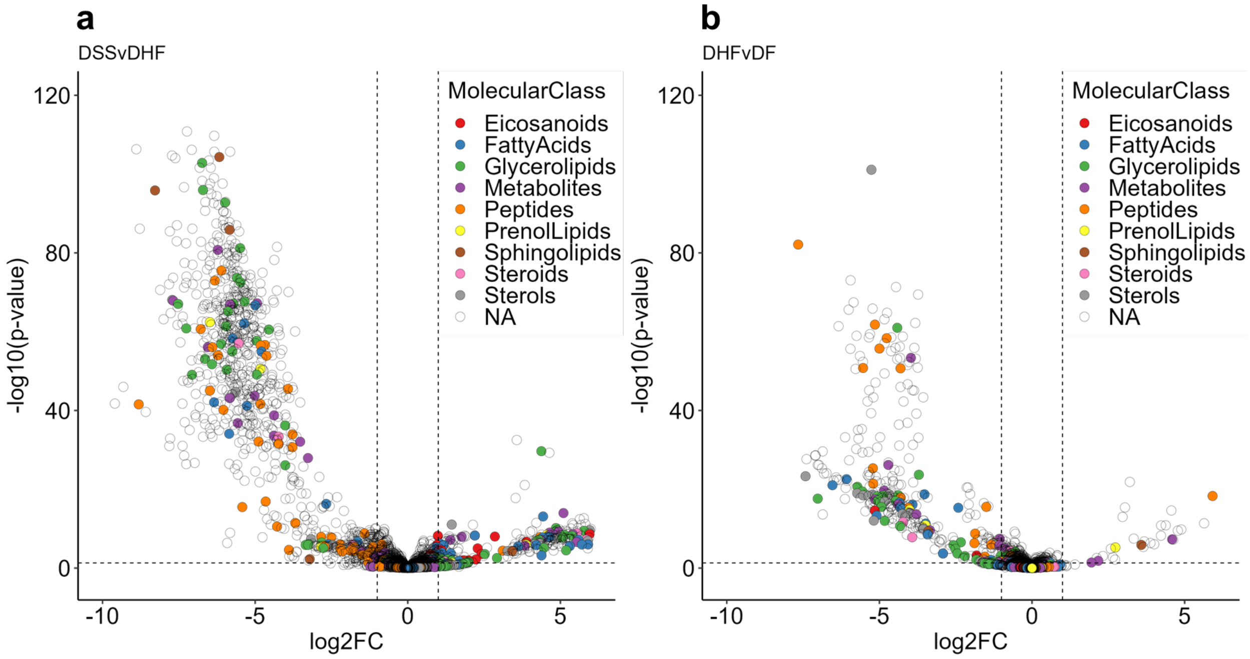


**Figure S1.** Volcano plots displaying differential abundance of features between two pairwise disease state comparisons. (**a**) Negative $log_2FC$ values can be interpreted as either a lower feature abundance in DSS or a greater abundance in DHF. Positive $log_2FC$ values can be interpreted as either a greater feature abundance in DSS or a lower abundance in DHF. (**b**) Negative $log_2FC$ values can be interpreted as either a lower feature abundance in DHF or a greater abundance in DF. Positive $log_2FC$ values can be interpreted as either a greater feature abundance in DHF or a lower abundance in DF. Colored points represent a manually curated list of 266 tentatively identified (based on accurate mass) potentially biologically relevant molecules, where color represents the tentative molecular class. Noncolored points represent features that were either manually defined as not biologically relevant, or tentative identification based on accurate mass was unsuccessful. The horizontal dotted line represents an adjusted p-value of 0.05 and the vertical dotted lines represent $log_2FC = \pm 1$.

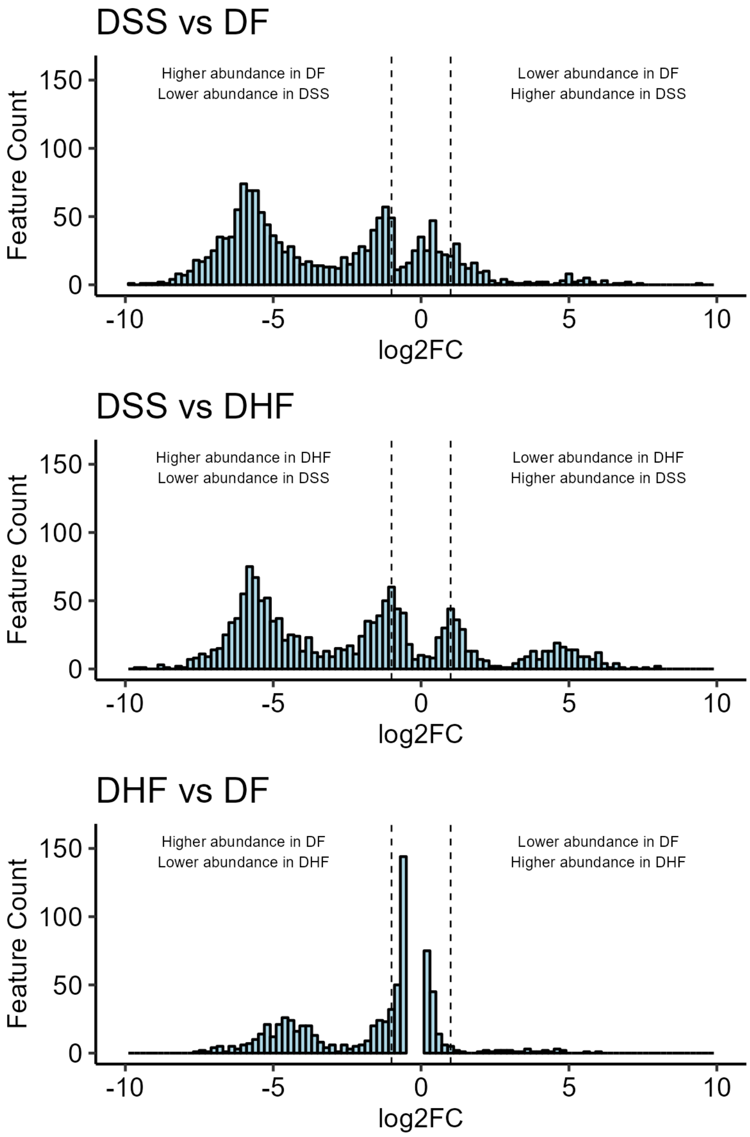


**Figure S2.** Frequency distribution of log2 FCs for the 1512 features that had significant differential abundance (adjusted p-value < 0.05) for three pairwise comparisons (top: DSS vs DF, middle: DSS vs DHF, bottom: DHF vs DF). $Log_2FC$ bin width was set to 0.2. Vertical lines represent $log_2FC = \pm 1$. Log2 FCs were calculated with the R package *limma.*

**Table S2.** Metabolites of interest LC-MS metrics, 32 were included in the biomarker classification panel.

| KEGG ID | Feature | Metabolite | Neutral Chemical Formula | Ion Type | Exact Mass (*m/z*) | Accurate Mass (*m/z*) | Mass Error (ppm) | Retention Time (min) | Relevance | Conf Level | Panel |
|---|---|---|---|---|---|---|---|---|---|---|---|
| C05464 | MZ450.3226_8.73 | Glycodeoxycholic acid | C26H43NO5 | [M+H]+ | 450.3214 | 450.3226 | 2.67 | 8.73 | Bile Acid | 2 | Yes |
| C02838 | MZ286.2017_5.87 | 2-Octenoylcarnitine | C15H27NO4 | [M+H]+ | 286.2013 | 286.2017 | 1.45 | 5.87 | Carnitine | 2 | Yes |
| C00318 | MZ162.1126_1.07 | Carnitine | C7H15NO3 | [M]+ | 162.1125 | 162.1126 | 0.62 | 1.07 | Carnitine | 1 | No |
| C03017 | MZ218.1387_1.69 | Propionylcarnitine | C10H19NO4 | [M+H]+ | 218.1387 | 218.1387 | 0.07 | 1.69 | Carnitine | 2 | Yes |
| | MZ255.2323_13.93 | Palmitoleic acid | C16H30O2 | [M+H]+ | 255.2319 | 255.2323 | 1.74 | 13.93 | Fatty Acid Biosynthesis | 2 | Yes |
| | MZ331.2846_14.28 | MG(16:0) | C19H38O4 | [M+H]+ | 331.2843 | 331.2846 | 0.91 | 14.28 | Glycerolipid | 2 | No |
| | MZ496.3422_11.66 | LPC(16:0) sn1 | C24H50NO7P | [M+H]+ | 496.3398 | 496.3422 | 4.91 | 11.59 | Glycerophospholipid | 2 | Yes |
| | MZ496.3422_11.38 | LPC(16:0) sn2 | C24H50NO7P | [M+H]+ | 496.3398 | 496.3422 | 4.91 | 11.38 | Glycerophospholipid | 2 | Yes |
| | MZ522.3579_11.99 | LPC(18:1) | C26H52NO7P | [M+H]+ | 522.3554 | 522.3579 | 4.76 | 11.99 | Glycerophospholipid | 2 | Yes |
| | MZ518.3249_8.69 | LPC(18:3) | C26H48NO7P | [M+H]+ | 518.3241 | 518.3249 | 1.54 | 8.69 | Glycerophospholipid | 2 | No |
| | MZ524.3735_13.07 | LysoPC(18:0) sn1 | C26H54NO7P | [M+H]+ | 524.3711 | 524.3735 | 4.64 | 13.07 | Glycerophospholipid | 2 | Yes |
| | MZ524.3736_12.78 | LysoPC(18:0) sn2 | C26H54NO7P | [M+H]+ | 524.3711 | 524.3736 | 4.83 | 12.78 | Glycerophospholipid | 2 | Yes |
| | MZ542.3242_8.83 | LysoPC(20:5) | C28H48NO7P | [M+H]+ | 542.3241 | 542.3242 | 0.11 | 8.83 | Glycerophospholipid | 2 | No |
| | MZ568.3387_11.59 | LysoPC(22:6) | C30H50NO7P | [M+H]+ | 568.3398 | 568.3387 | -1.87 | 11.66 | Glycerophospholipid | 2 | Yes |
| | MZ526.2942_10.98 | LysoPE(22:6) | C27H44NO7P | [M+H]+ | 526.2928 | 526.2942 | 2.63 | 10.98 | Glycerophospholipid | 2 | Yes |
| C06427 | MZ279.2319_13.45 | alpha-Linolenic acid | C18H30O2 | [M+H]+ | 279.2319 | 279.2319 | 0.16 | 13.45 | n-3 Fatty Acid Metabolism | 1 | Yes |
| C06429 | MZ329.2493_13.79 | Docosahexaenoic acid | C22H32O2 | [M+H]+ | 329.2475 | 329.2493 | 5.45 | 13.79 | n-3 Fatty Acid Metabolism | 2 | Yes |
| | MZ303.2321_11.77 | Eicosapentaenoic acid | C20H30O2 | [M+H]+ | 303.2319 | 303.2321 | 0.80 | 11.77 | n-3 Fatty Acid Metabolism | 3 | No |
| | MZ305.2446_11.85 | Eicosatetraenoic acid | C20H32O2 | [M+H]+ | 305.2475 | 305.2446 | -9.50 | 11.85 | n-3 Fatty Acid Metabolism | 3 | No |
| | MZ277.216_11.03 | Stearidonic acid | C18H28O2 | [M+H]+ | 277.2162 | 277.2160 | -0.74 | 11.03 | n-3 Fatty Acid Metabolism | 3 | No |
| C00219 | MZ305.2481_14.01 | Arachidonic acid | C20H32O2 | [M+H]+ | 305.2475 | 305.2481 | 1.95 | 14.5 | n-6 Fatty Acid Metabolism | 1 | Yes |
| C03242 | MZ307.2632_14.48 | dihomo-gamma-linolenic acid | C20H34O2 | [M+H]+ | 307.2632 | 307.2632 | 0.14 | 14.9 | n-6 Fatty Acid Metabolism | 1 | Yes |
| C06426 | MZ279.2319_13.57 | Gamma-Linolenic acid | C18H30O2 | [M+H]+ | 279.2319 | 279.2319 | 0.16 | 13.57 | n-6 Fatty Acid Metabolism | 1 | Yes |
| | MZ359.2217_9.34 | Hepoxilin A3 | C20H32O4 | [M+Na]+ | 359.2193 | 359.2217 | 6.68 | 9.34 | n-6 Fatty Acid Metabolism | 3 | No |
| | MZ319.2269_9.87 | Leukotriene A4 | C20H30O3 | [M+H]+ | 319.2273 | 319.2269 | -1.25 | 9.87 | n-6 Fatty Acid Metabolism | 3 | No |
| C01595 | MZ281.2482_14.2 | Linoleic acid | C18H32O2 | [M+H]+ | 281.2475 | 281.2482 | 2.47 | 14.7 | n-6 Fatty Acid Metabolism | 1 | Yes |
| | MZ353.2315_8.32 | Prostaglandin D2 | C20H32O5 | [M+H]+ | 353.2323 | 353.2315 | -2.26 | 8.32 | n-6 Fatty Acid Metabolism | 3 | No |
| C00791 | MZ114.0664_0.98 | Creatinine | C4H7N3O | [M+H]+ | 114.0662 | 114.0664 | 1.75 | 0.98 | Other | 1 | No |
| C01586 | MZ180.0661_3.89 | Hippuric acid | C9H9NO3 | [M+H]+ | 180.0655 | 180.0661 | 3.23 | 3.89 | Other | 2 | Yes |
| | MZ203.1365_1.84 | Leucyl-alanine | C9H18N2O3 | [M+H]+ | 203.1390 | 203.1365 | -12.40 | 1.84 | Peptide | 2 | Yes |
| C00079 | MZ166.0865_2.27 | L-Phenylalanine | C9H11NO2 | [M+H]+ | 166.0863 | 166.0865 | 1.48 | 2.27 | Peptide | 2 | Yes |
| | MZ213.1229_1.38 | L-prolyl-proline | C10H16N2O3 | [M+H]+ | 213.1234 | 213.1229 | -2.35 | 1.38 | Peptide | 2 | Yes |
| | MZ313.1558_4.82 | Phenylalanylphenylalanine | C18H20N2O3 | [M+H]+ | 313.1547 | 313.1558 | 3.61 | 4.82 | Peptide | 2 | Yes |
| C00148 | MZ116.0708_1.09 | Proline | C5H9NO2 | [M+H]+ | 116.0706 | 116.0708 | 1.68 | 1.09 | Peptide | 2 | Yes |
| | MZ231.1697_2.89 | Valyl-Leucine | C11H22N2O3 | [M+H]+ | 231.1703 | 231.1697 | -2.60 | 2.89 | Peptide | 2 | Yes |
| C00262 | MZ137.0465_1.35 | Hypoxanthine | C5H4N4O | [M+H]+ | 137.0463 | 137.0465 | 1.46 | 1.32 | Purine Metabolism | 1 | Yes |
| C00294 | MZ269.09_1.39 | Inosine | C10H12N4O5 | [M+H]+ | 269.0886 | 269.09 | 5.20 | 1.39 | Purine Metabolism | 1 | No |
| C01017 | MZ221.0917_1.67 | 5-hydroxytryptophan | C11H12N2O3 | [M+H]+ | 221.0921 | 221.0915 | -2.57 | 1.59 | Tryptophan Metabolism | 3 | Yes |
| C00954 | MZ176.0711_5.67 | Indoleacetic acid | C10H9NO2 | [M+H]+ | 176.0706 | 176.0711 | 2.84 | 5.6 | Tryptophan Metabolism | 1 | Yes |
| | MZ188.0707_3.09 | Indoleacrylic acid | C11H9NO2 | [M+H]+ | 188.0706 | 188.0707 | 0.51 | 3.09 | Tryptophan Metabolism | 3 | No |
| C02043 | MZ206.0825_5.04 | Indolelactic acid | C11H11NO3 | [M+H]+ | 206.0812 | 206.0825 | 6.46 | 5.1 | Tryptophan Metabolism | 1 | Yes |
| | MZ190.0866_6.5 | Indolepropionic acid | C11H11NO2 | [M+H]+ | 190.0863 | 190.0863 | 0.00 | 6.5 | Tryptophan Metabolism | 1 | Yes |
| C00328 | MZ209.0922_2.23 | L-Kynurenine | C10H12N2O3 | [M+H]+ | 209.0921 | 209.0922 | 0.63 | 1.98 | Tryptophan Metabolism | 1 | Yes |
| C00078 | MZ205.0977_3.09 | L-Tryptophan | C11H12N2O2 | [M+H]+ | 205.0972 | 205.0977 | 2.66 | 3.0 | Tryptophan Metabolism | 1 | Yes |
| C00780 | MZ177.1011_1.7 | Serotonin | C10H12N2O | [M+H]+ | 177.1022 | 177.1011 | -6.43 | 1.5 | Tryptophan Metabolism | 1 | Yes |
| C06124 | MZ380.257_10.14 | Sphingosine-1-phosphate | C18H38NO5P | [M+H]+ | 380.256 | 380.257 | 2.63 | 10.14 | Sphingolipid | 3 | No |
| C01120 | MZ382.2723_10.45 | Sphinganine-1-phosphate | C18H40NO5P | [M+H]+ | 382.2717 | 382.2723 | 1.57 | 10.45 | Sphingolipid | 3 | No |
| C00319 | MZ322.2746_14.2 | Sphingosine (d18:1) | C18H37NO2 | [M+Na]+ | 322.2717 | 322.2746 | 9 | 14.2 | Sphingolipid | 3 | No |
| C00836 | MZ324.2905_14.98 | Sphinganine (d18:0) | C18H39NO2 | [M+Na]+ | 324.2873 | 324.2905 | 9.87 | 14.98 | Sphingolipid | 3 | No |

**Table S3.** Metabolites of interest and their $log_2FC$ and adjusted p-values for each pairwise disease state comparison.

| KEGG ID | Feature | Metabolite | DSS.DF $log_2FC$ | DSS.DF pval | DHF.DF $log_2FC$ | DHF.DF pval | DSS.DHF $log_2FC$ | DSS.DHF pval | DSS.ND $log_2FC$ | DSS.ND pval | DHF.ND $log_2FC$ | DHF.ND pval | DF.ND $log_2FC$ | DF.ND pval | Relevance |
|---|---|---|---|---|---|---|---|---|---|---|---|---|---|---|---|
| C05464 | MZ450.3226_8.73 | Glycodeoxycholic acid | 0.795 | 1.23E-02 | 0.187 | 5.81E-01 | 0.608 | 1.23E-01 | 0.974 | 1.29E-03 | 0.366 | 1.75E-01 | 0.179 | 3.02E-01 | Bile Acid |
| C02838 | MZ286.2017_5.87 | 2-Octenoylcarnitine | 0.856 | 1.48E-03 | -0.006 | 9.85E-01 | 0.862 | 7.56E-03 | 0.781 | 2.97E-03 | -0.081 | 7.74E-01 | -0.075 | 6.73E-01 | Carnitine |
| C00318 | MZ162.1126_1.07 | Carnitine | -1.198 | 3.70E-13 | -0.443 | 1.35E-03 | -0.754 | 1.16E-04 | -1.269 | 4.65E-15 | -0.515 | 4.40E-05 | -0.071 | 4.86E-01 | Carnitine |
| C03017 | MZ218.1387_1.69 | Propionylcarnitine | -0.615 | 3.13E-03 | -0.339 | 7.20E-02 | -0.276 | 3.12E-01 | -0.534 | 8.62E-03 | -0.258 | 1.46E-01 | 0.081 | 5.25E-01 | Carnitine |
| | MZ255.2323_13.93 | Palmitoleic acid | 1.143 | 1.53E-04 | 0.358 | 2.25E-01 | 0.785 | 3.70E-02 | 1.134 | 1.15E-04 | 0.35 | 1.89E-01 | -0.008 | 9.72E-01 | Fatty Acid Biosynthesis |
| | MZ331.2846_14.28 | MG(16:0) | 1.086 | 2.21E-06 | 0.229 | 3.32E-01 | 0.857 | 1.95E-03 | 1.432 | 1.60E-10 | 0.574 | 1.33E-03 | 0.346 | 9.63E-04 | Glycerolipid |
| | MZ496.3422_11.66 | LPC(16:0) sn1 | 0.013 | 8.93E-01 | -0.046 | 5.50E-01 | 0.06 | 5.49E-01 | -0.078 | 2.91E-01 | -0.138 | 1.57E-02 | -0.092 | 5.30E-03 | Glycerophospholipid |
| | MZ496.3422_11.38 | LPC(16:0) sn2 | -0.011 | 9.44E-01 | -0.092 | 4.29E-01 | 0.081 | 6.03E-01 | -0.12 | 3.00E-01 | -0.201 | 2.57E-02 | -0.109 | 4.46E-02 | Glycerophospholipid |
| | MZ524.3735_13.07 | LPC(18:0) sn1 | -0.025 | 8.43E-01 | -0.251 | 1.48E-03 | 0.226 | 6.01E-02 | -0.112 | 2.58E-01 | -0.337 | 2.06E-06 | -0.087 | 6.84E-02 | Glycerophospholipid |
| | MZ524.3736_12.78 | LPC(18:0) sn2 | -0.048 | 7.44E-01 | -0.175 | 9.84E-02 | 0.128 | 4.08E-01 | -0.222 | 5.21E-02 | -0.35 | 4.43E-05 | -0.175 | 6.45E-04 | Glycerophospholipid |
| | MZ522.3579_11.99 | LPC(18:1) | -0.156 | 8.21E-02 | -0.07 | 4.26E-01 | -0.086 | 4.52E-01 | -0.27 | 1.06E-03 | -0.184 | 5.30E-03 | -0.114 | 2.96E-03 | Glycerophospholipid |
| | MZ518.3249_8.69 | LPC(18:3) | -0.372 | 6.39E-01 | -1.25 | 2.69E-02 | 0.878 | 3.02E-01 | 1.253 | 5.12E-02 | 0.376 | 5.47E-01 | 1.626 | 6.99E-09 | Glycerophospholipid |
| | MZ542.3242_8.83 | LPC(20:5) | -0.179 | 8.45E-01 | -0.582 | 4.17E-01 | 0.404 | 6.80E-01 | 1.336 | 5.32E-02 | 0.932 | 1.14E-01 | 1.515 | 5.96E-07 | Glycerophospholipid |
| | MZ568.3387_11.59 | LPC(22:6) | -0.658 | 5.96E-07 | -0.33 | 3.28E-03 | -0.328 | 4.96E-02 | -0.862 | 2.15E-11 | -0.534 | 6.40E-08 | -0.205 | 6.63E-04 | Glycerophospholipid |
| | MZ526.2942_10.98 | LPE(22:6) | 0.363 | 7.86E-02 | 0.396 | 1.96E-02 | -0.033 | 9.07E-01 | 0.433 | 2.53E-02 | 0.466 | 1.89E-03 | 0.069 | 5.71E-01 | Glycerophospholipid |
| C06427 | MZ279.2319_13.45 | alpha-Linolenic acid | 1.259 | 3.46E-03 | -0.164 | 7.37E-01 | 1.423 | 5.35E-03 | 1.161 | 5.65E-03 | -0.263 | 5.23E-01 | -0.098 | 7.33E-01 | n-3 Fatty Acid Metabolism |
| C06429 | MZ329.2493_13.79 | Docosahexaenoic acid | 1.205 | 2.24E-11 | 0.153 | 4.24E-01 | 1.052 | 6.42E-07 | 1.003 | 1.35E-08 | -0.05 | 8.03E-01 | -0.203 | 1.97E-02 | n-3 Fatty Acid Metabolism |
| | MZ303.2321_11.77 | Eicosapentaenoic acid | 2.327 | 1.31E-07 | 0.037 | 9.50E-01 | 2.29 | 1.01E-05 | 2.897 | 1.91E-11 | 0.607 | 1.14E-01 | 0.57 | 5.64E-03 | n-3 Fatty Acid Metabolism |
| | MZ305.2446_11.85 | Eicosatetraenoic acid | 2.072 | 1.60E-03 | -1.817 | 6.20E-04 | 3.888 | 2.03E-07 | 3.141 | 5.51E-07 | -0.747 | 1.92E-01 | 1.069 | 2.32E-04 | n-3 Fatty Acid Metabolism |
| | MZ277.216_11.03 | Stearidonic acid | 1.023 | 6.21E-06 | -0.083 | 7.56E-01 | 1.107 | 3.18E-05 | 1.228 | 2.55E-08 | 0.122 | 5.95E-01 | 0.205 | 7.29E-02 | n-3 Fatty Acid Metabolism |
| C00219 | MZ305.2481_14.01 | Arachidonic acid | 0.918 | 2.76E-10 | -0.069 | 6.76E-01 | 0.987 | 6.41E-09 | 0.868 | 1.18E-09 | -0.119 | 3.87E-01 | -0.05 | 5.95E-01 | n-6 Fatty Acid Metabolism |
| C03242 | MZ307.2632_14.48 | Dihomo-gamma-linolenic acid | 0.914 | 6.13E-08 | -0.048 | 8.19E-01 | 0.962 | 1.18E-06 | 0.791 | 1.82E-06 | -0.171 | 2.70E-01 | -0.123 | 1.78E-01 | n-6 Fatty Acid Metabolism |
| C06426 | MZ279.2319_13.57 | gamma-Linolenic acid | 1.218 | 8.15E-05 | 0.048 | 9.05E-01 | 1.17 | 1.51E-03 | 1.068 | 4.27E-04 | -0.101 | 7.56E-01 | -0.15 | 4.16E-01 | n-6 Fatty Acid Metabolism |
| | MZ359.2217_9.34 | Hepoxilin A3 | 2.568 | 6.13E-03 | -4.102 | 3.59E-09 | 6.67 | 2.27E-10 | 8.443 | 3.33E-21 | 1.774 | 1.54E-02 | 5.876 | 1.93E-46 | n-6 Fatty Acid Metabolism |
| | MZ319.2269_9.87 | Leukotriene A4 | 2.053 | 1.53E-03 | 0.06 | 9.44E-01 | 1.993 | 1.06E-02 | 3.345 | 6.32E-08 | 1.352 | 7.60E-03 | 1.292 | 5.21E-06 | n-6 Fatty Acid Metabolism |
| C01595 | MZ281.2482_14.2 | Linoleic acid | 0.866 | 1.28E-06 | 0.106 | 5.95E-01 | 0.76 | 3.59E-04 | 0.784 | 7.79E-06 | 0.023 | 9.13E-01 | -0.083 | 4.45E-01 | n-6 Fatty Acid Metabolism |
| | MZ353.2315_8.32 | Prostaglandin D2 | 2.791 | 7.54E-18 | 0.593 | 4.39E-02 | 2.197 | 5.47E-09 | 3.046 | 1.61E-21 | 0.848 | 7.41E-04 | 0.255 | 1.36E-01 | n-6 Fatty Acid Metabolism |
| C00791 | MZ114.0664_0.98 | Creatinine | -0.509 | 3.53E-04 | 0.082 | 5.97E-01 | -0.592 | 4.09E-04 | -0.404 | 4.05E-03 | 0.188 | 1.25E-01 | 0.106 | 1.62E-01 | Other |
| C01586 | MZ180.0661_3.89 | Hippuric acid | -1.034 | 2.32E-03 | -0.855 | 2.21E-03 | -0.179 | 7.11E-01 | -1.221 | 1.87E-04 | -1.042 | 3.62E-05 | -0.187 | 3.28E-01 | Other |
| | MZ203.1365_1.84 | Leucyl-alanine | -11.345 | 1.27E-96 | -7.66 | 7.75E-83 | -3.684 | 3.71E-12 | -12.054 | 3.77E-107 | -8.37 | 1.82E-98 | -0.71 | 6.54E-04 | Peptide |
| C00079 | MZ166.0865_2.27 | L-Phenylalanine | 0.057 | 7.60E-01 | -0.143 | 3.33E-01 | 0.2 | 3.00E-01 | 0.107 | 4.91E-01 | -0.094 | 5.07E-01 | 0.05 | 5.93E-01 | Peptide |
| | MZ213.1229_1.38 | L-prolyl-proline | -6.261 | 2.05E-58 | -4.762 | 4.58E-59 | -1.498 | 4.65E-04 | -6.013 | 1.59E-56 | -4.515 | 1.48E-57 | 0.248 | 2.07E-01 | Peptide |
| | MZ313.1558_4.82 | Phenylalanylphenylalanine | -3.189 | 2.78E-11 | -1.887 | 4.74E-07 | -1.303 | 3.11E-02 | -4.024 | 1.62E-17 | -2.722 | 2.15E-14 | -0.835 | 1.19E-04 | Peptide |
| C00148 | MZ116.0708_1.09 | Proline | -0.428 | 2.31E-03 | -0.478 | 1.05E-05 | 0.05 | 8.05E-01 | -0.424 | 1.89E-03 | -0.474 | 4.70E-06 | 0.004 | 9.71E-01 | Peptide |
| | MZ231.1697_2.89 | Valyl-Leucine | -7.684 | 2.06E-55 | -5.535 | 1.66E-51 | -2.149 | 6.37E-05 | -8.199 | 4.89E-63 | -6.05 | 2.12E-62 | -0.516 | 1.81E-02 | Peptide |
| C00262 | MZ137.0465_1.35 | Hypoxanthine | -2.139 | 8.64E-14 | -1.076 | 1.77E-06 | -1.063 | 2.22E-03 | -2.37 | 4.74E-17 | -1.307 | 1.00E-09 | -0.231 | 1.28E-01 | Purine Metabolism |
| C00294 | MZ269.09_1.39 | Inosine | -6.266 | 1.52E-57 | -0.696 | 3.33E-02 | -5.57 | 1.63E-37 | -6.51 | 8.34E-63 | -0.939 | 8.61E-04 | -0.243 | 2.24E-01 | Purine Metabolism |
| C01017 | MZ221.0917_1.67 | 5-Hydroxytryptophan | -0.378 | 1.31E-02 | -0.033 | 8.61E-01 | -0.345 | 6.35E-02 | -0.082 | 6.10E-01 | 0.264 | 2.82E-02 | 0.297 | 5.70E-06 | Tryptophan Metabolism |
| C00954 | MZ176.0711_5.67 | Indoleacetic acid | -0.387 | 1.68E-01 | -0.486 | 3.25E-02 | 0.099 | 7.90E-01 | -0.6 | 1.84E-02 | -0.699 | 3.29E-04 | -0.213 | 1.02E-01 | Tryptophan Metabolism |
| | MZ188.0707_3.09 | Indoleacrylic acid | -0.345 | 3.84E-02 | -0.3 | 3.45E-02 | -0.046 | 8.46E-01 | -0.672 | 1.30E-05 | -0.626 | 1.05E-07 | -0.327 | 3.37E-06 | Tryptophan Metabolism |
| C02043 | MZ206.0825_5.04 | Indolelactic acid | -0.026 | 9.00E-01 | -0.096 | 5.55E-01 | 0.07 | 7.48E-01 | -0.143 | 3.61E-01 | -0.213 | 9.43E-02 | -0.117 | 1.34E-01 | Tryptophan Metabolism |
| | MZ190.0866_6.5 | Indolepropionic acid | -1.222 | 1.83E-04 | -1.218 | 1.41E-06 | -0.004 | 9.93E-01 | -1.477 | 2.96E-06 | -1.473 | 8.49E-10 | -0.255 | 1.35E-01 | Tryptophan Metabolism |
| C00328 | MZ209.0922_2.23 | L-Kynurenine | -0.231 | 3.18E-01 | 0.096 | 6.68E-01 | -0.327 | 2.15E-01 | 0.145 | 5.02E-01 | 0.472 | 2.60E-03 | 0.376 | 2.68E-05 | Tryptophan Metabolism |
| C00078 | MZ205.0977_3.09 | L-Tryptophan | -0.361 | 3.69E-02 | -0.33 | 2.30E-02 | -0.03 | 9.00E-01 | -0.694 | 1.41E-05 | -0.664 | 5.36E-08 | -0.334 | 4.80E-06 | Tryptophan Metabolism |
| C00780 | MZ177.1011_1.7 | Serotonin | -9.442 | 2.43E-138 | -0.549 | 3.12E-02 | -8.893 | 4.97E-107 | -9.938 | 9.41E-150 | -1.045 | 9.85E-07 | -0.495 | 1.06E-04 | Tryptophan Metabolism |
| C06124 | MZ380.257_10.14 | Sphingosine-1-phosphate | -0.73 | 6.53E-10 | -0.546 | 2.00E-09 | -0.184 | 2.46E-01 | -0.778 | 1.91E-11 | -0.594 | 1.46E-11 | -0.049 | 5.11E-01 | Sphingolipid |
| C01120 | MZ382.2723_10.45 | Sphinganine-1-phosphate | -0.957 | 6.59E-12 | -0.554 | 3.66E-07 | -0.403 | 2.07E-02 | -0.75 | 3.94E-08 | -0.347 | 1.52E-03 | 0.207 | 1.25E-03 | Sphingolipid |
| C00319 | MZ322.2746_14.2 | Sphingosine (d18:1) | 0.925 | 9.24E-08 | 0.09 | 6.41E-01 | 0.835 | 4.55E-05 | 0.867 | 3.25E-07 | 0.032 | 8.76E-01 | -0.058 | 6.04E-01 | Sphingolipid |
| C00836 | MZ324.2905_14.98 | Sphinganine (d18:0) | 0.507 | 1.12E-03 | 0.183 | 2.24E-01 | 0.324 | 9.96E-02 | 0.482 | 1.50E-03 | 0.158 | 2.56E-01 | -0.025 | 8.14E-01 | Sphingolipid |

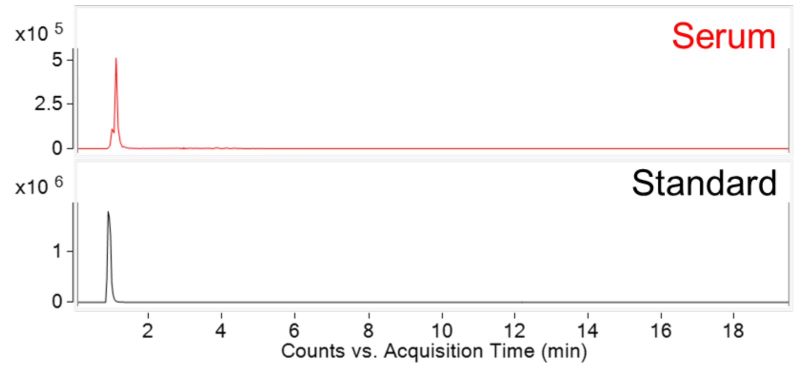


**Figure S3a**. Carnitine detected as $[M+H]^+$ at *m/z* 162.1125. **Top**) Chromatographic retention time was matched between synthetic standard and the endogenous molecule detected in serum. **Bottom**) Collision-induced dissociation product ion spectra comparison for synthetic and endogenous molecules at collision energy values of 10 and 40.

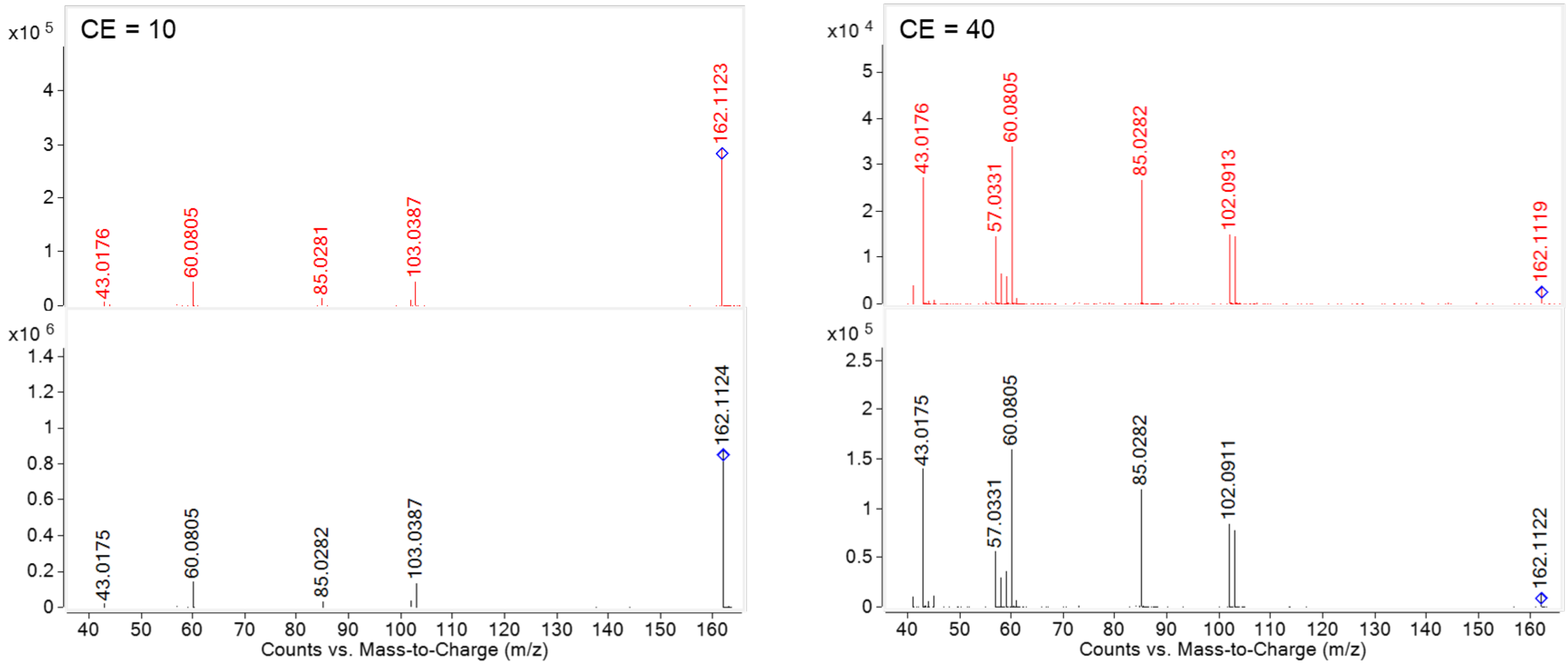

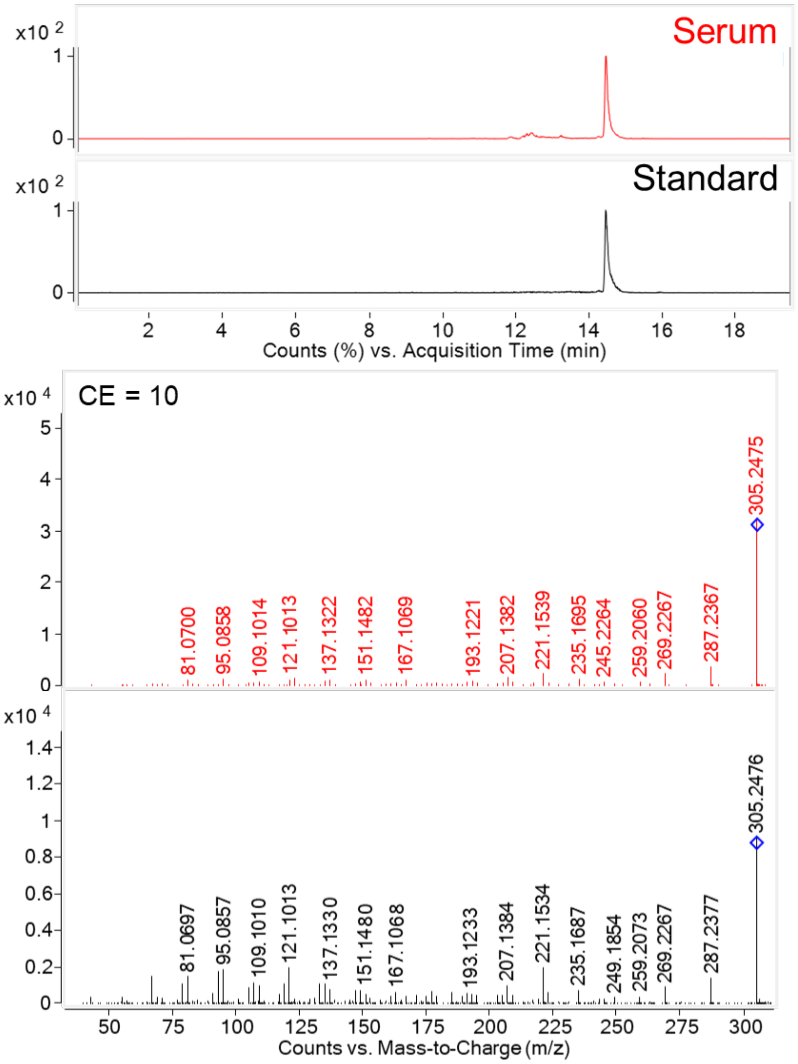


**Figure S3b**. Arachidonic acid detected as $[M+H]^+$ at *m/z* 305.2475. **Top**) Chromatographic retention time was matched between synthetic standard and the endogenous molecule detected in serum. **Bottom**) Collision-induced dissociation product ion spectra comparison for synthetic and endogenous molecules at collision energy values of 10 and 40.

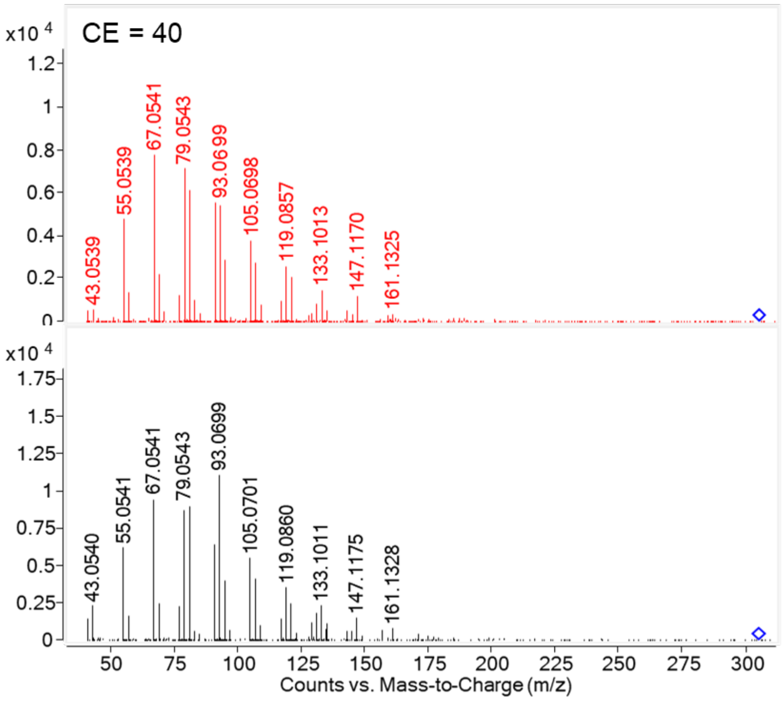

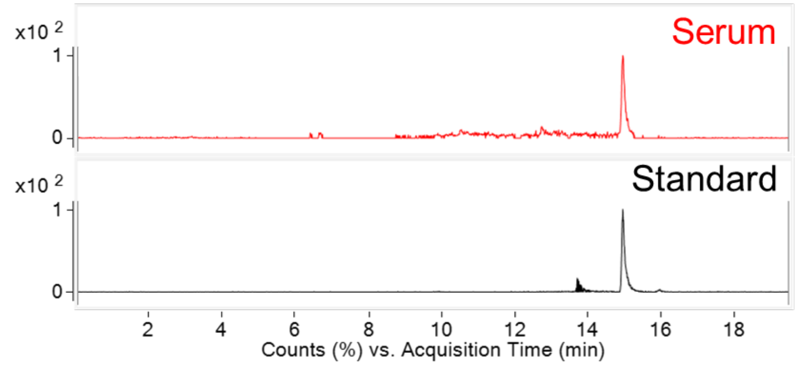


**Figure S3c**. Dihomo-γ-linolenic acid detected as $[M+H]^+$ at *m/z* 307.2632. **Top**) Chromatographic retention time was matched between synthetic standard and the endogenous molecule detected in serum. **Bottom**) Collision-induced dissociation product ion spectra comparison for synthetic and endogenous molecules at collision energy values of 10 and 40.

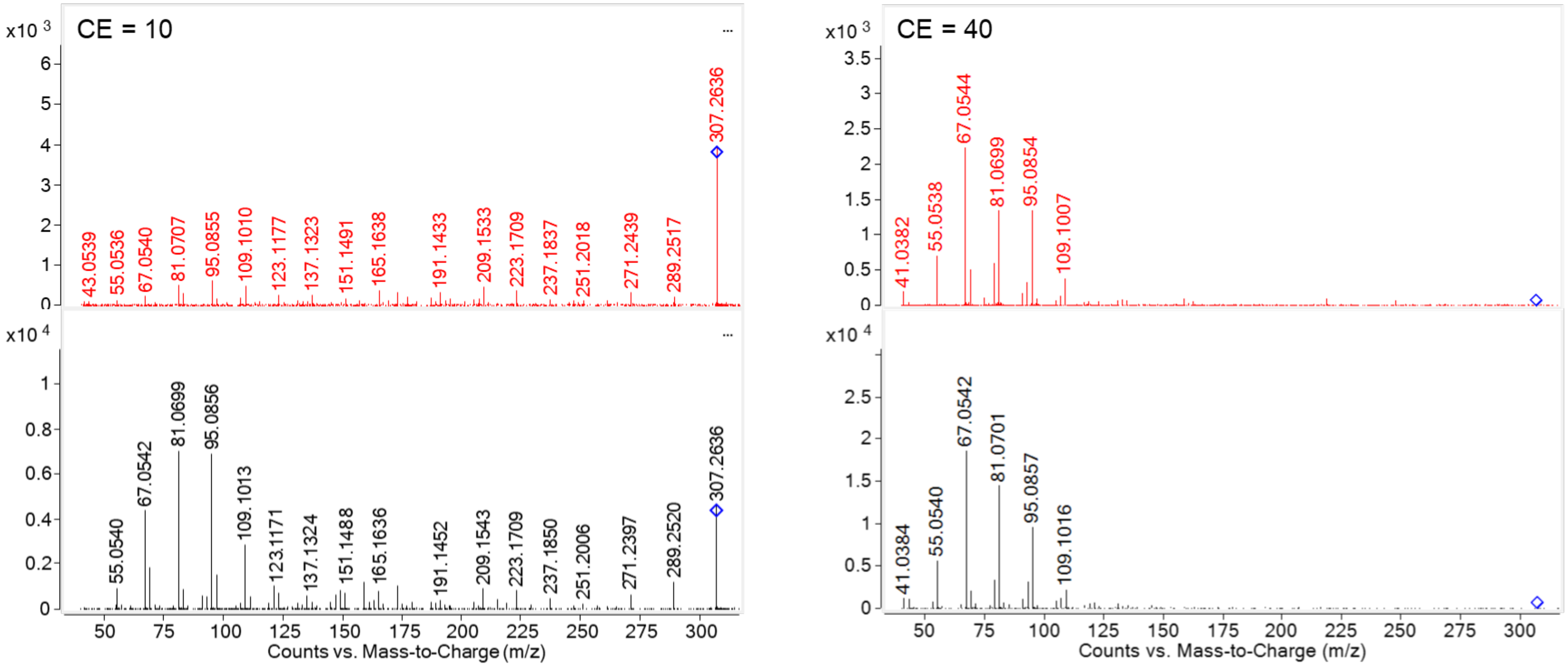

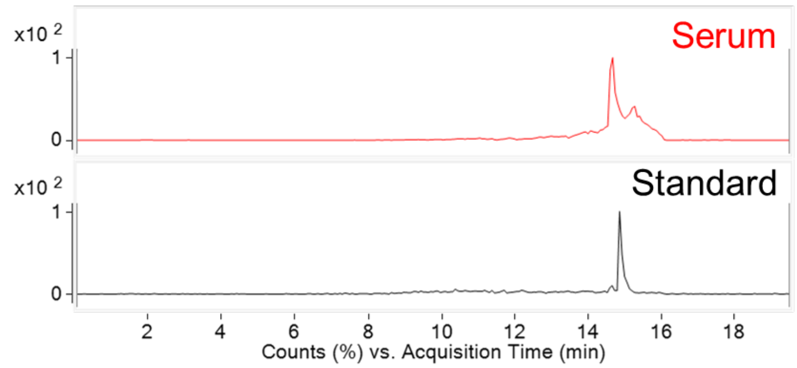


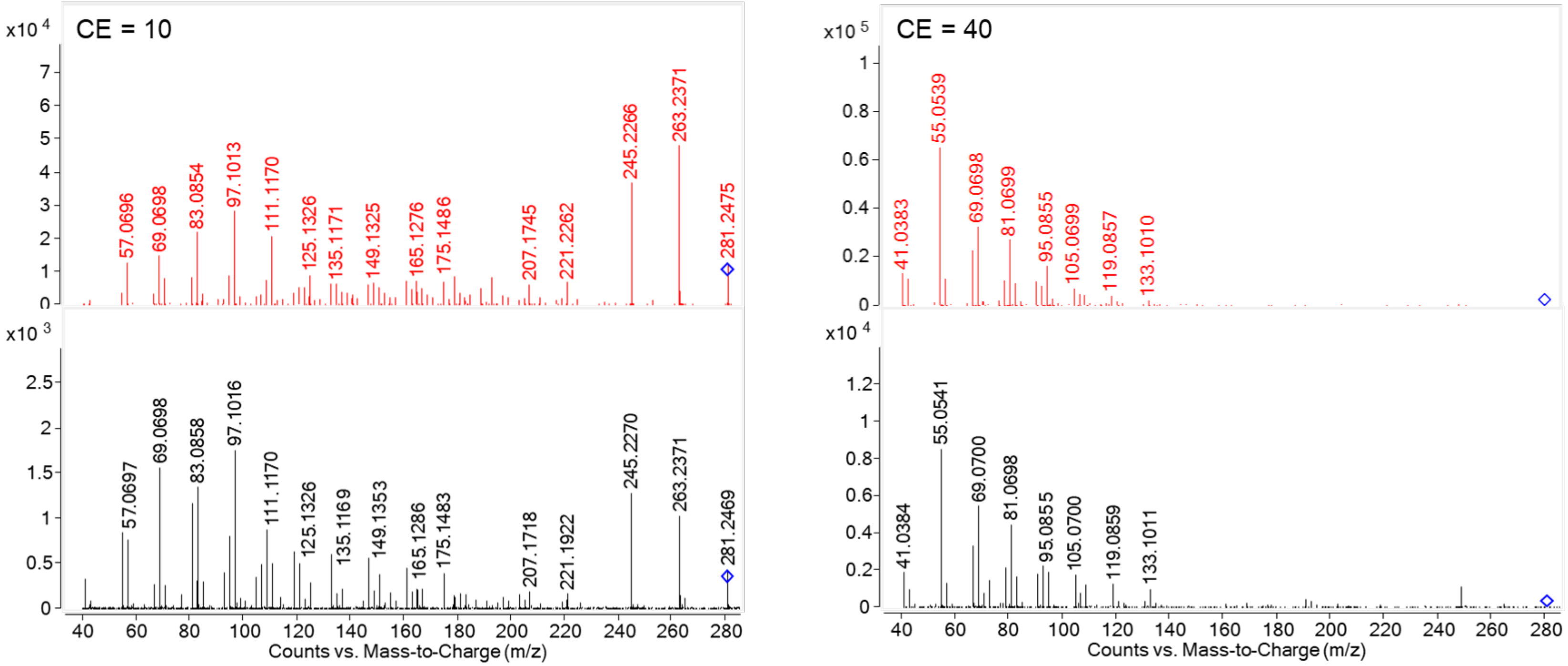


**Figure S3d**. Linoleic acid detected as $[M+H]^+$ at *m/z* 281.2475. **Top**) Chromatographic retention time was matched between synthetic standard and the endogenous molecule detected in serum. **Bottom**) Collision-induced dissociation product ion spectra comparison for synthetic and endogenous molecules at collision energy values of 10 and 40.

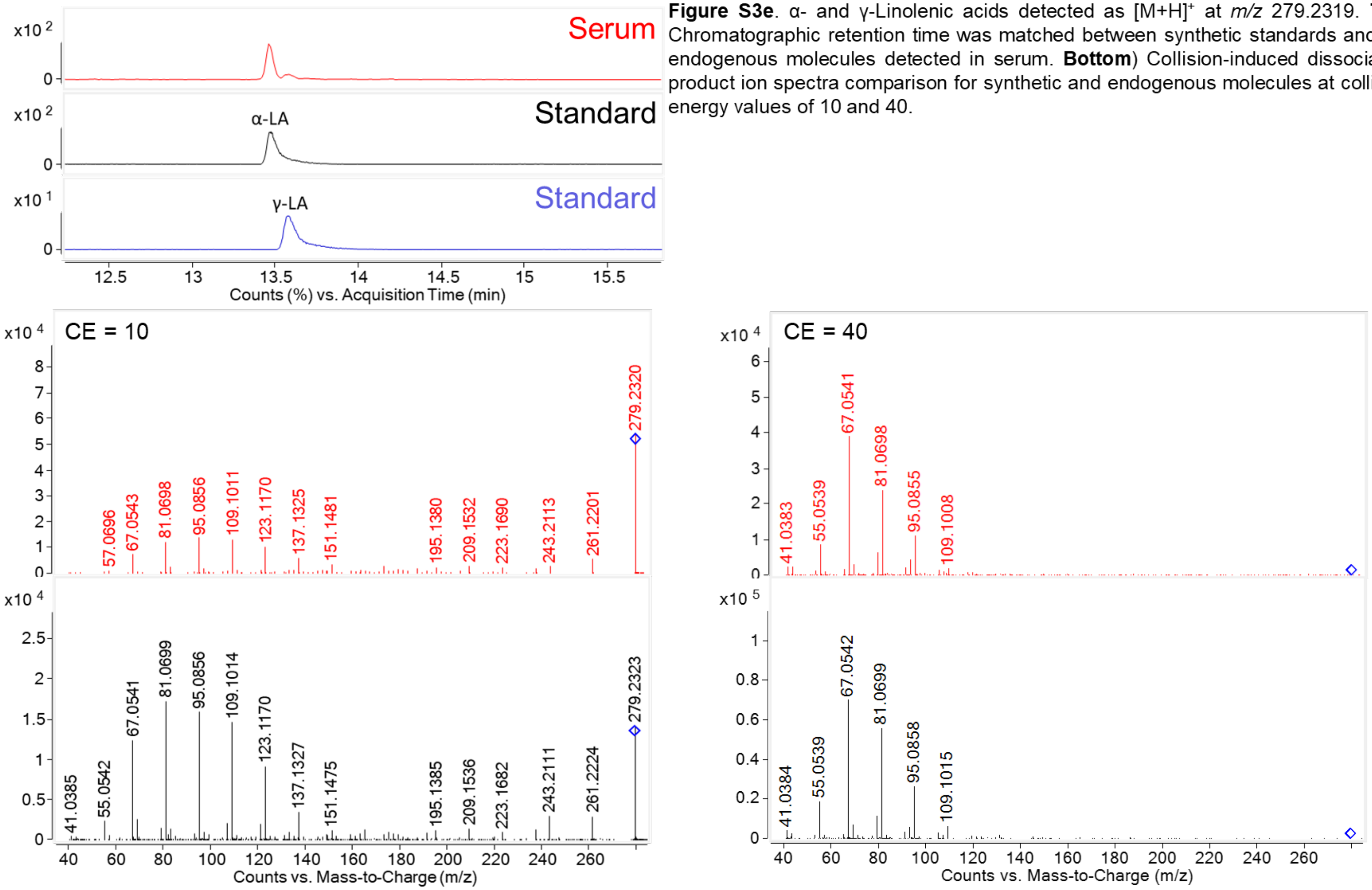


**Figure S3e**. α- and γ-Linolenic acids detected as $[M+H]^+$ at *m/z* 279.2319. **Top**) Chromatographic retention time was matched between synthetic standards and the endogenous molecules detected in serum. **Bottom**) Collision-induced dissociation product ion spectra comparison for synthetic and endogenous molecules at collision energy values of 10 and 40.

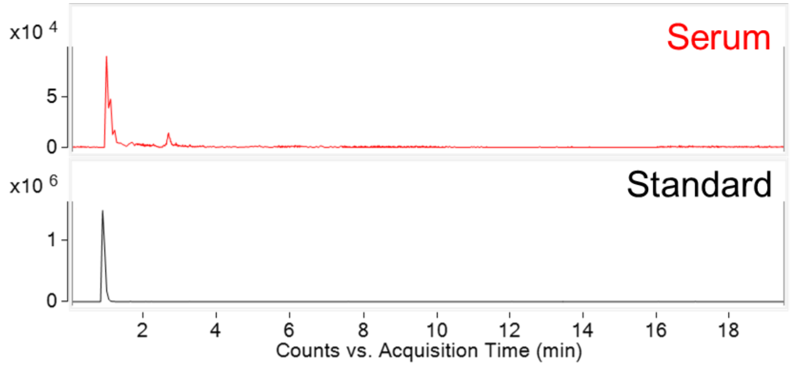


**Figure S3f**. Creatinine detected as $[M+H]^+$ at *m/z* 114.0662. **Top**) Chromatographic retention time was matched between synthetic standard and the endogenous molecule detected in serum. **Bottom**) Collision-induced dissociation product ion spectra comparison for synthetic and endogenous molecules at collision energy values of 10 and 40.

RT also matched creatinine-$d_3$ heavy-isotope labeled internal standard

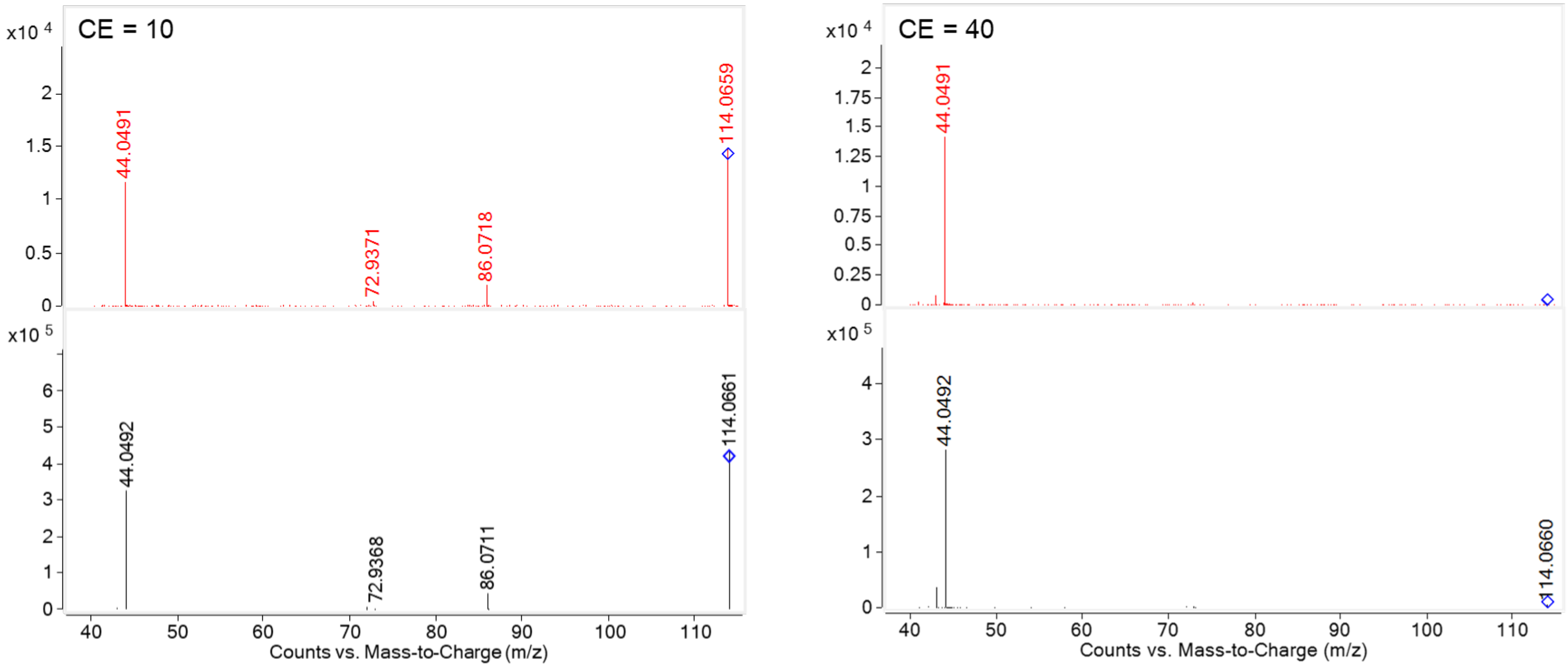

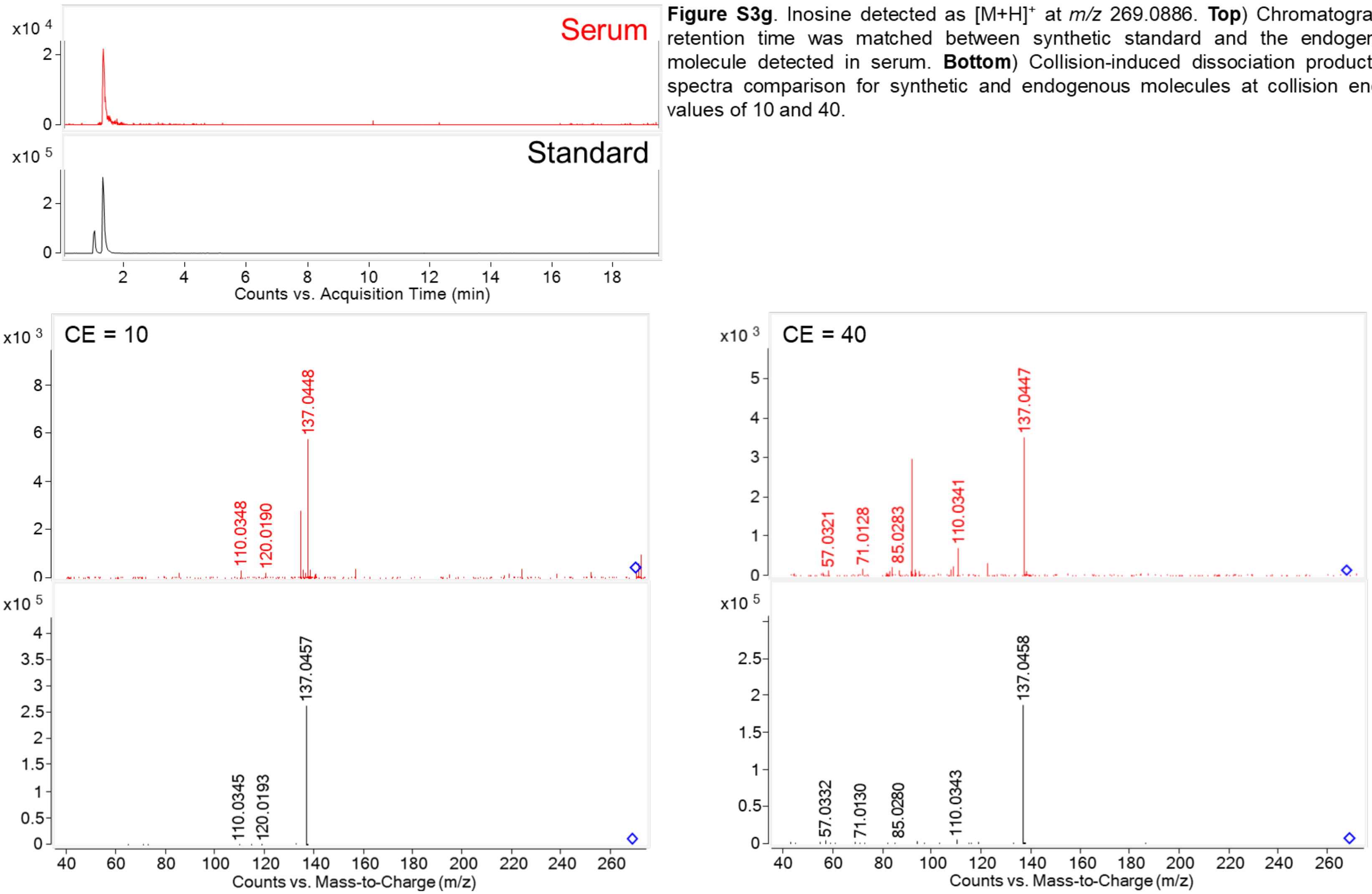


**Figure S3g**. Inosine detected as $[M+H]^+$ at *m/z* 269.0886. **Top**) Chromatographic retention time was matched between synthetic standard and the endogenous molecule detected in serum. **Bottom**) Collision-induced dissociation product ion spectra comparison for synthetic and endogenous molecules at collision energy values of 10 and 40.

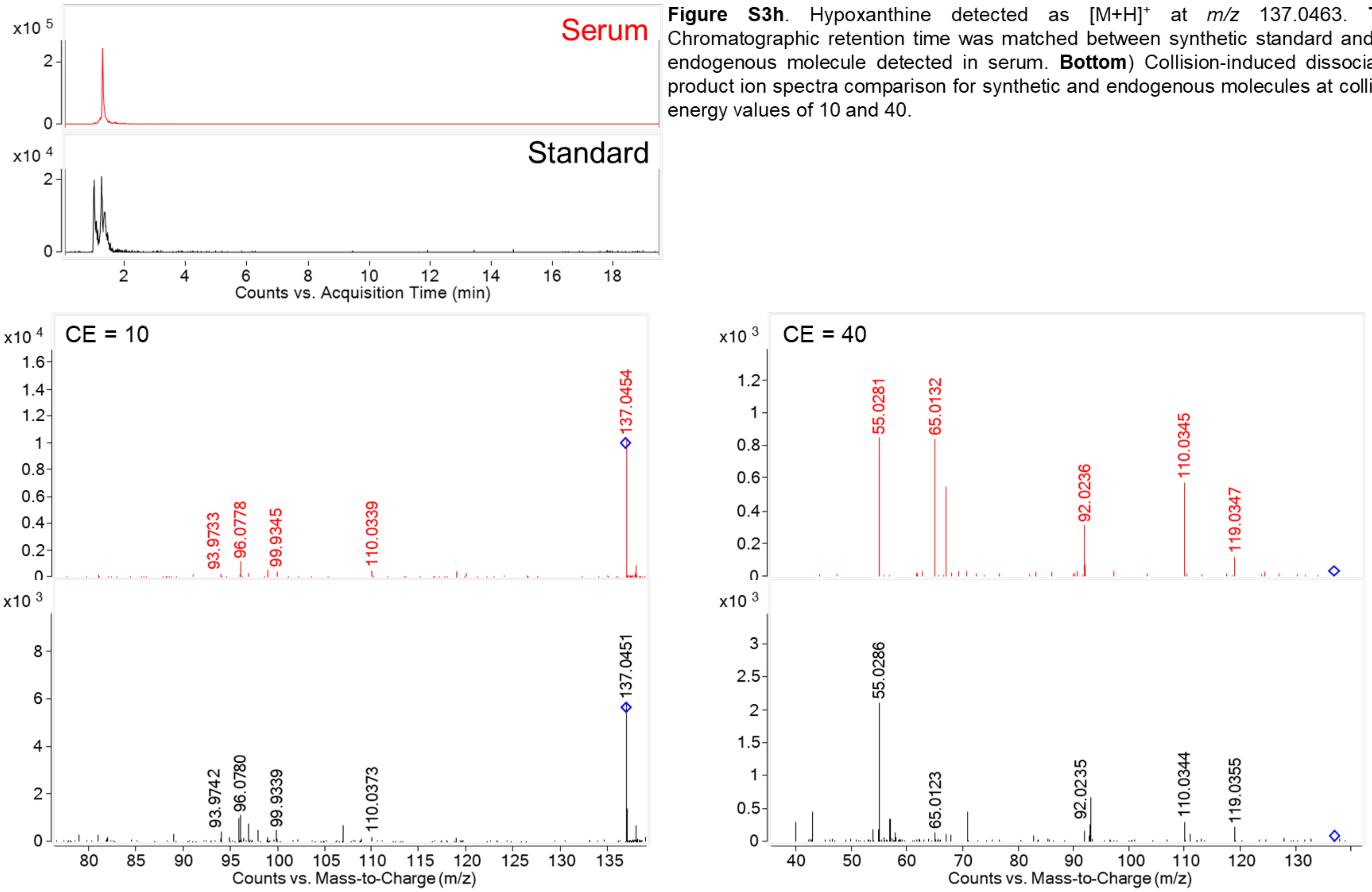


**Figure S3h**. Hypoxanthine detected as $[M+H]^+$ at *m/z* 137.0463. **Top**) Chromatographic retention time was matched between synthetic standard and the endogenous molecule detected in serum. **Bottom**) Collision-induced dissociation product ion spectra comparison for synthetic and endogenous molecules at collision energy values of 10 and 40.

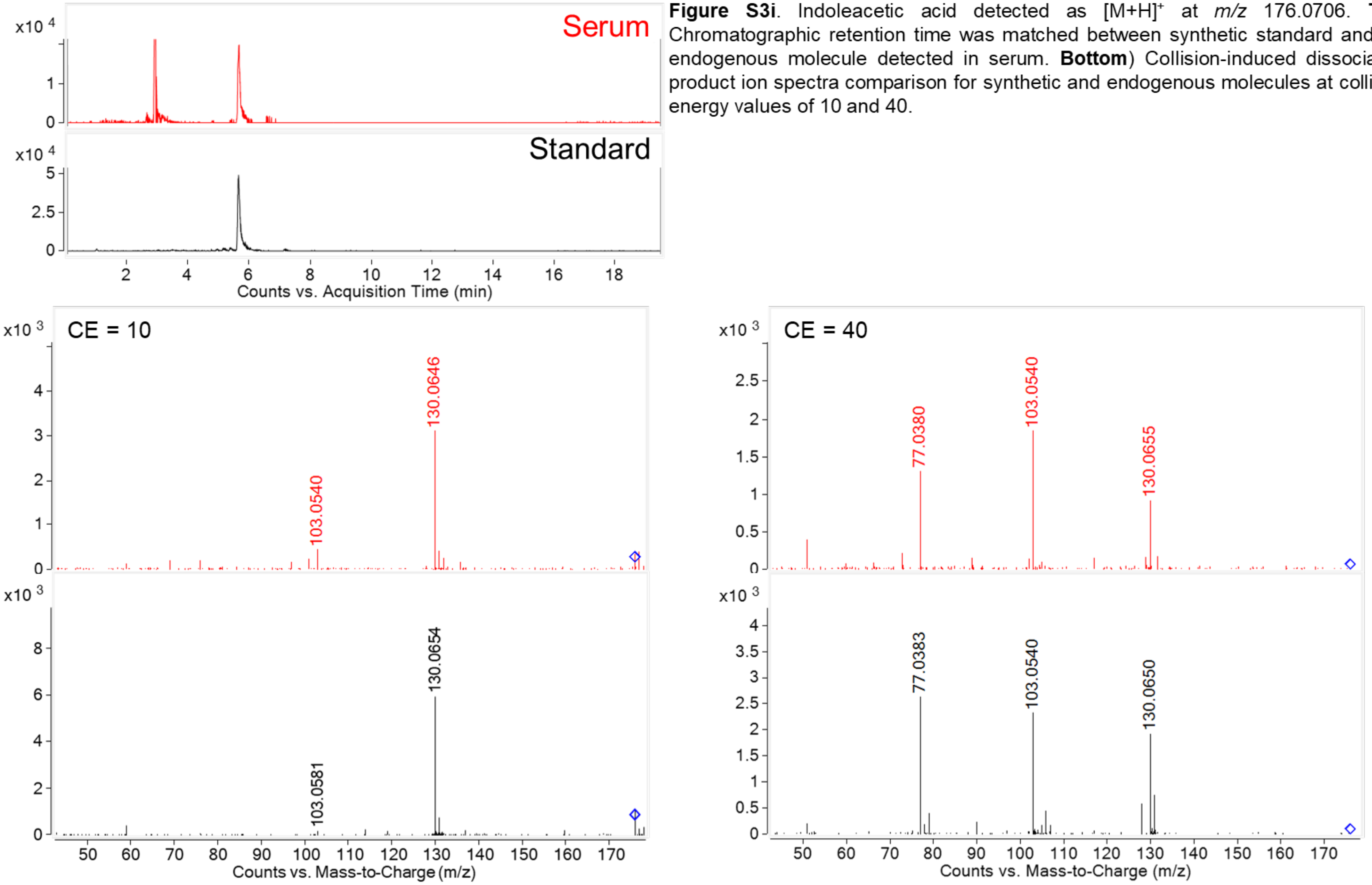


**Figure S3i**. Indoleacetic acid detected as $[M+H]^+$ at *m/z* 176.0706. **Top**) Chromatographic retention time was matched between synthetic standard and the endogenous molecule detected in serum. **Bottom**) Collision-induced dissociation product ion spectra comparison for synthetic and endogenous molecules at collision energy values of 10 and 40.

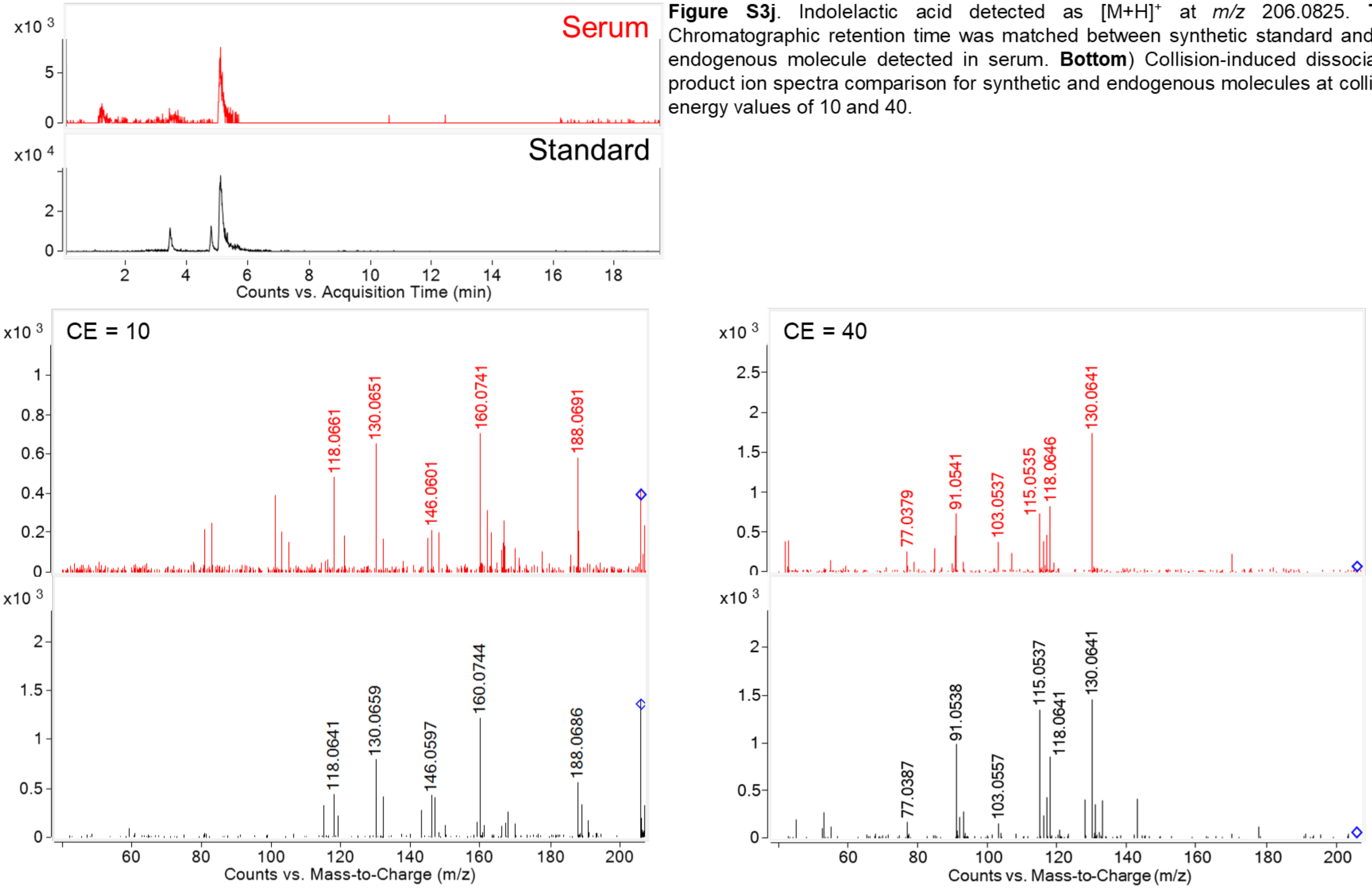


**Figure S3j**. Indolelactic acid detected as $[M+H]^+$ at *m/z* 206.0825. **Top**) Chromatographic retention time was matched between synthetic standard and the endogenous molecule detected in serum. **Bottom**) Collision-induced dissociation product ion spectra comparison for synthetic and endogenous molecules at collision energy values of 10 and 40.

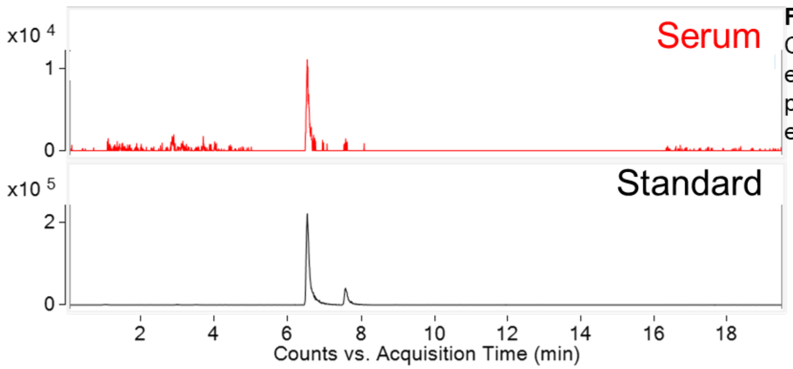


**Figure S3k**. Indolepropionic acid detected as $[M+H]^+$ at *m/z* 190.0863. **Top**) Chromatographic retention time was matched between synthetic standard and the endogenous molecule detected in serum. **Bottom**) Collision-induced dissociation product ion spectra comparison for synthetic and endogenous molecules at collision energy values of 10 and 40.

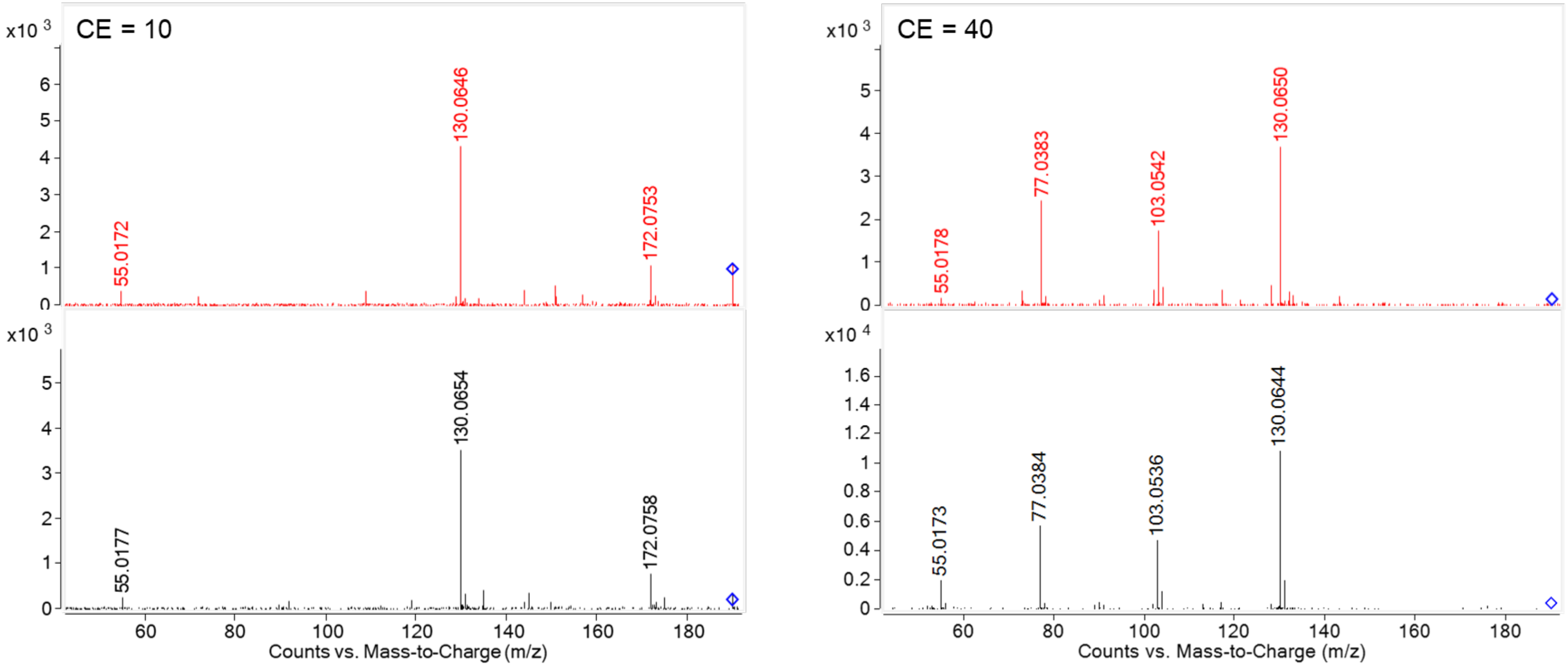

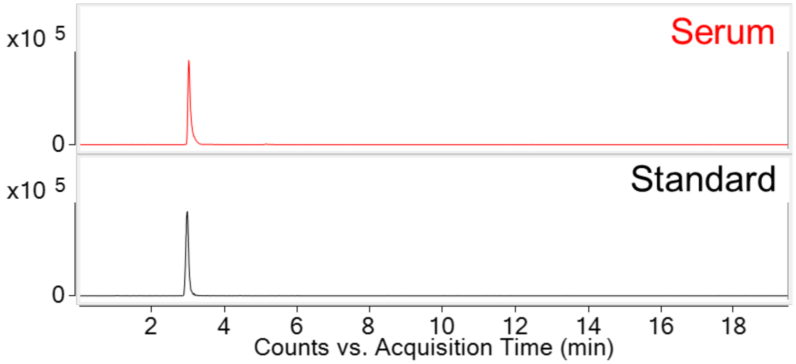


**Figure S3I**. Tryptophan detected as $[M+H]^+$ at *m/z* 205.0972. **Top**) Chromatographic retention time was matched between synthetic standard and the endogenous molecule detected in serum. **Bottom**) Collision-induced dissociation product ion spectra comparison for synthetic and endogenous molecules at collision energy values of 10 and 40.

RT also matched tryptophan-$d_5$ heavy-isotope labeled internal standard

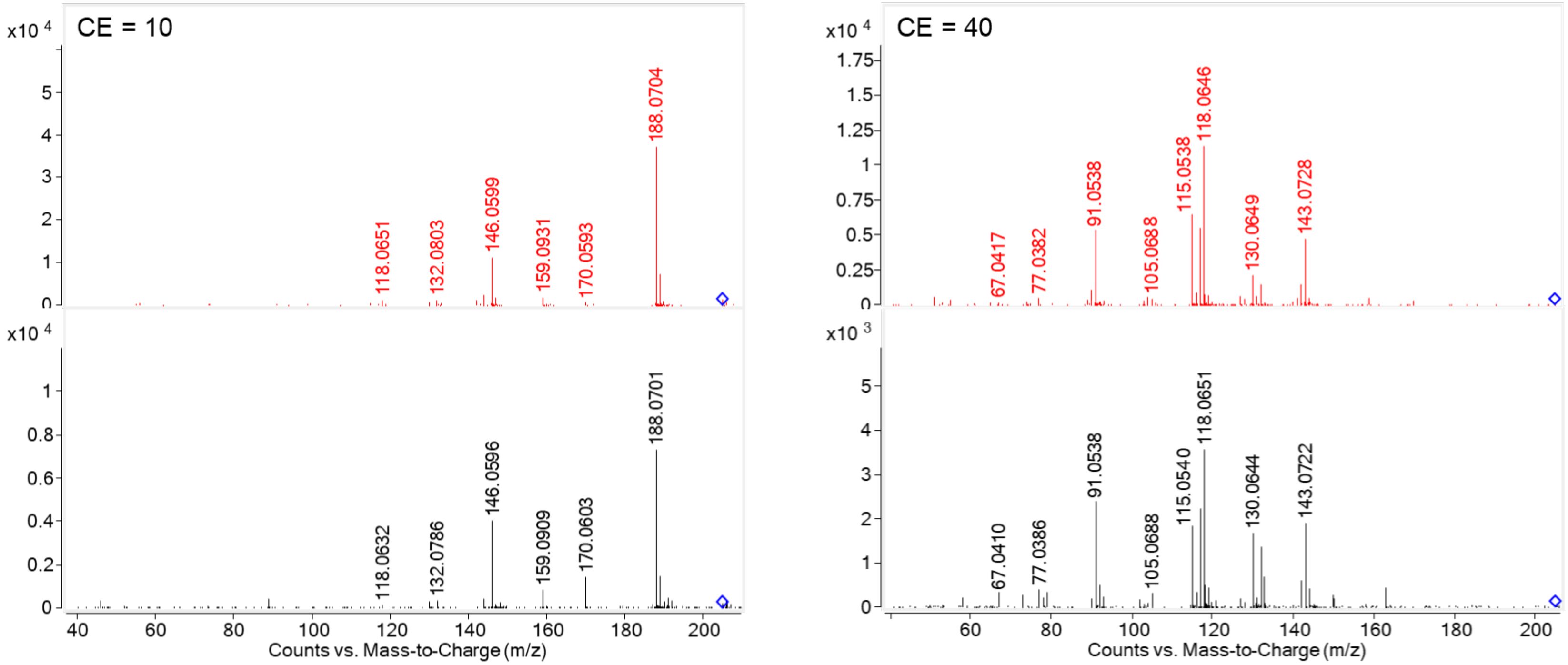

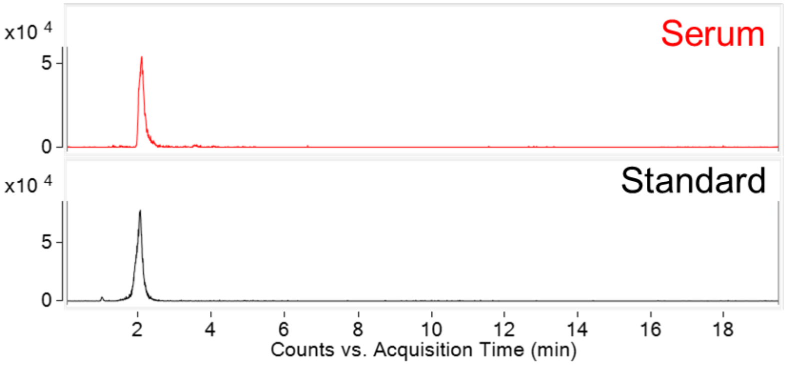


**Figure S3m**. Kynurenine detected as $[M+H]^+$ at *m/z* 209.0921. **Top**) Chromatographic retention time was matched between synthetic standard and the endogenous molecule detected in serum. **Bottom**) Collision-induced dissociation product ion spectra comparison for synthetic and endogenous molecules at collision energy values of 10 and 40.

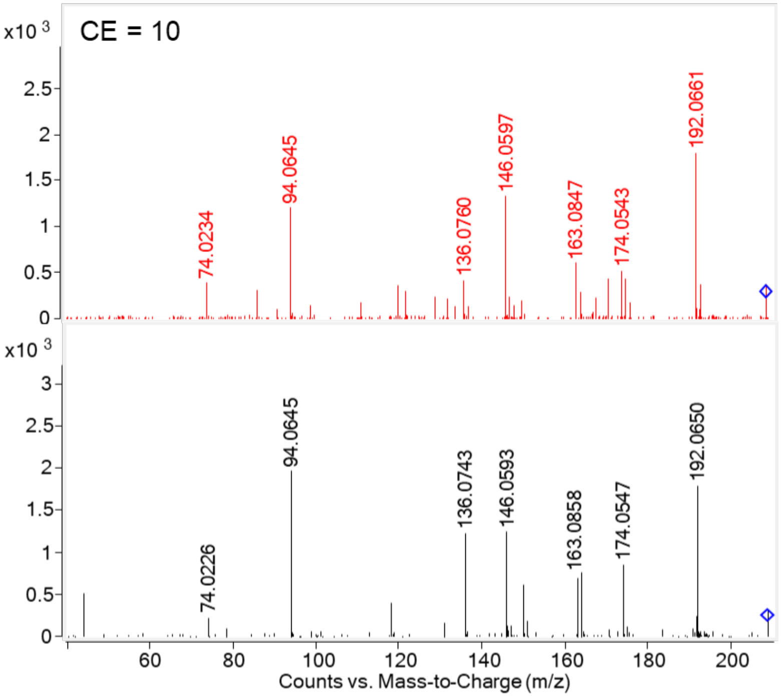


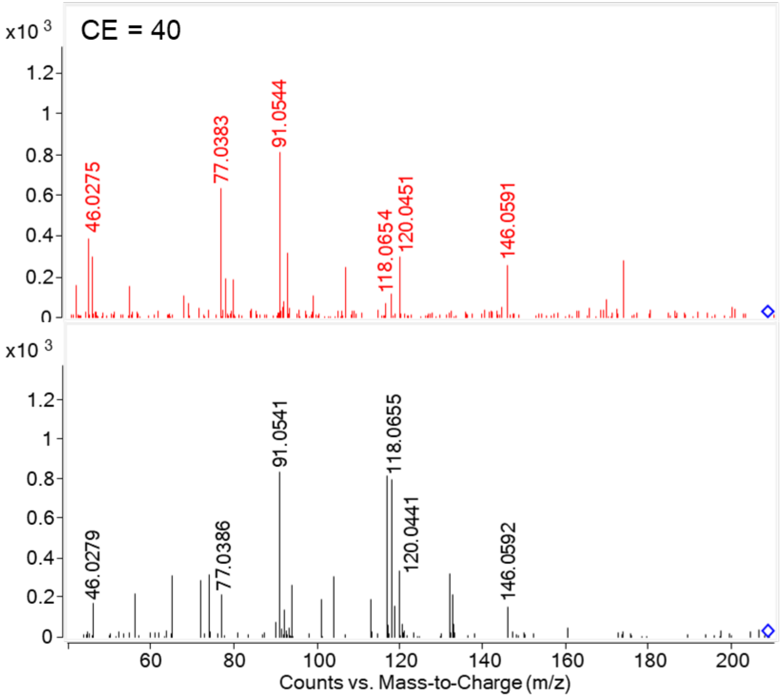

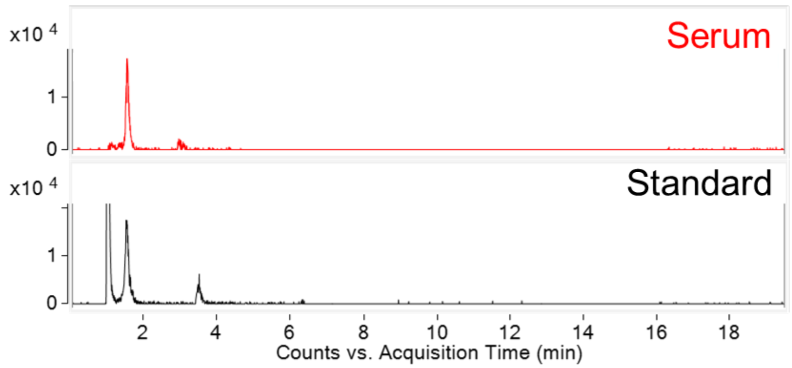


**Figure S3n**. Serotonin detected as $[M+H]^+$ at *m/z* 177.1022. **Top**) Chromatographic retention time was matched between synthetic standard and the endogenous molecule detected in serum. **Bottom**) Collision-induced dissociation product ion spectra comparison for synthetic and endogenous molecules at collision energy values of 10 and 40.

RT also matched serotonin-$d_4$ heavy-isotope labeled internal standard

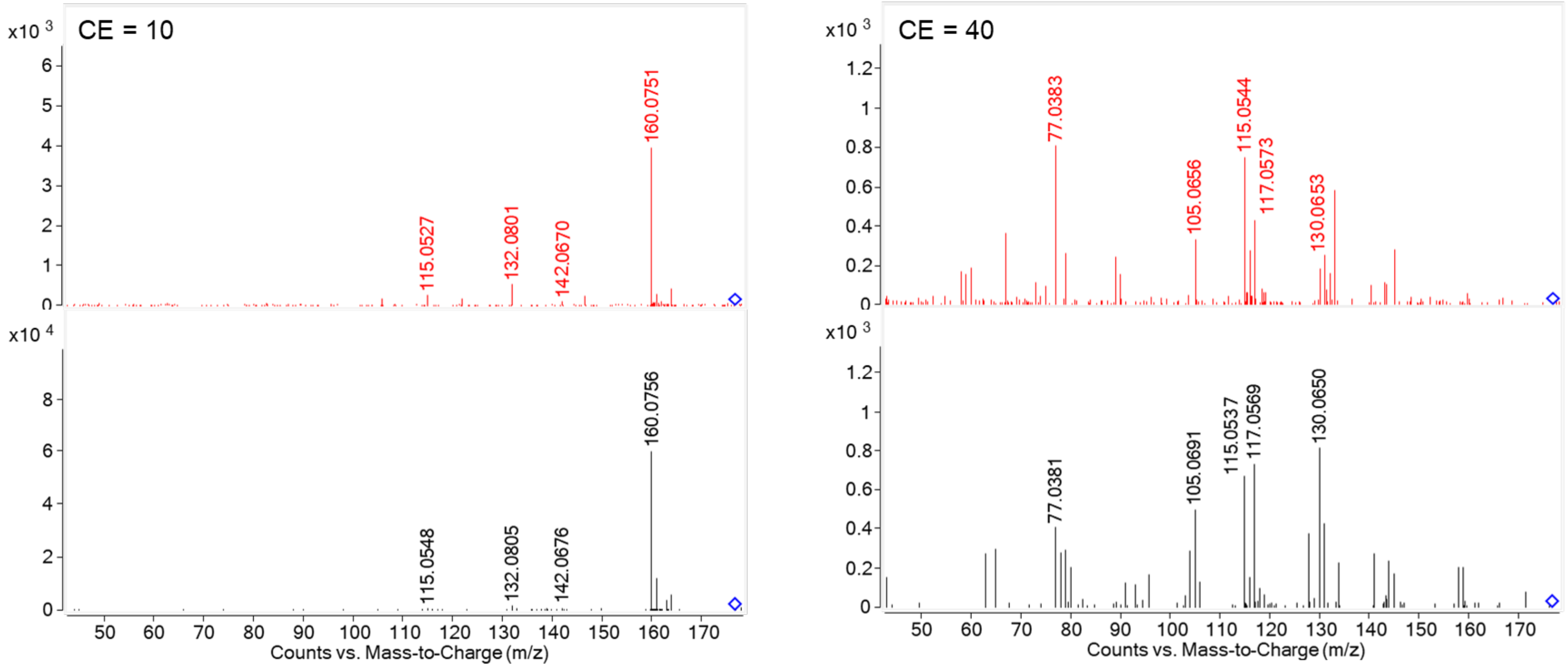

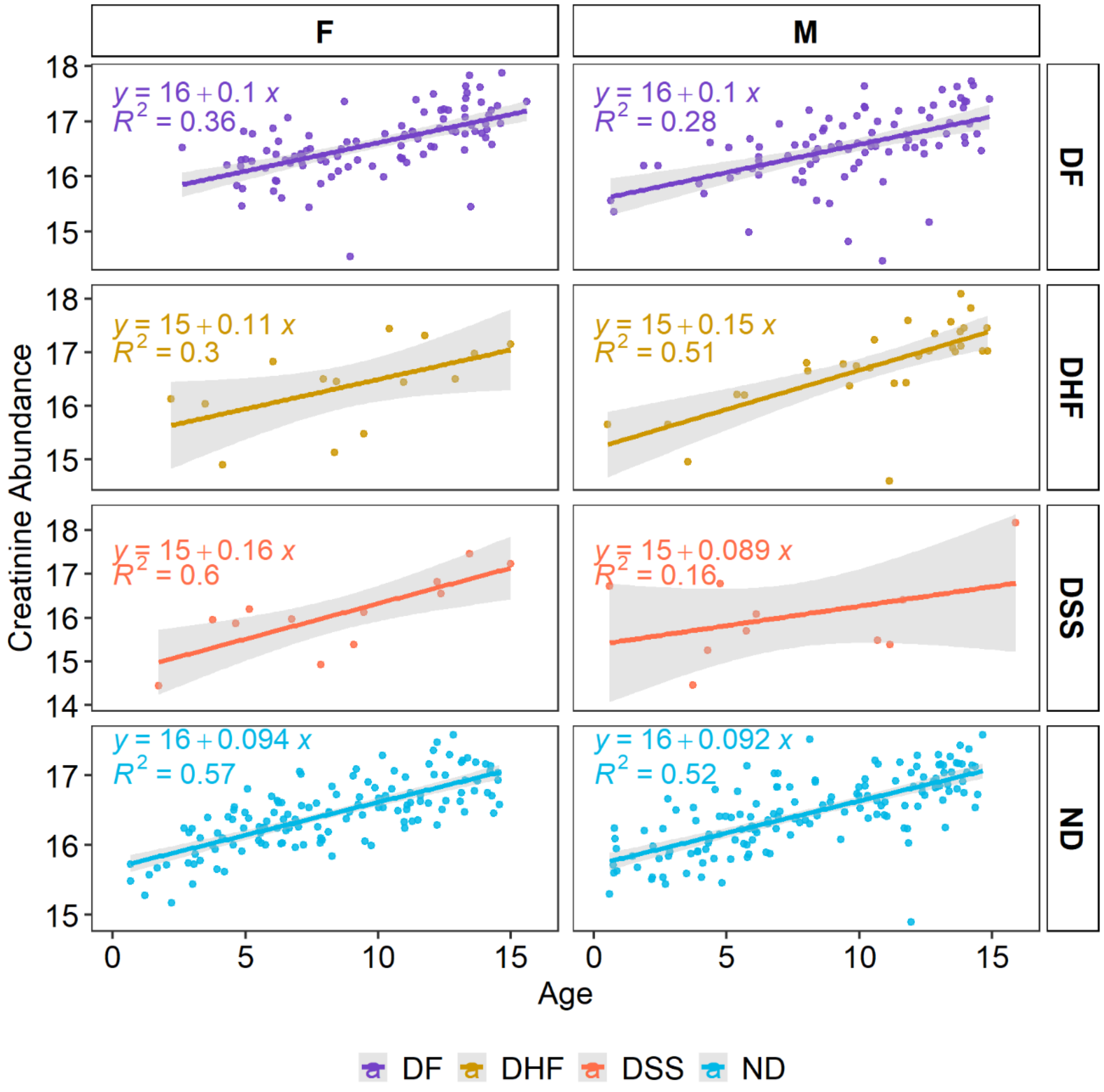


**Figure S4.** Linear regressions for creatinine abundance as a function of patient age for each combination of patient sex and disease state. The positive correlation between creatinine abundance and age is independent of patient sex and disease outcome.

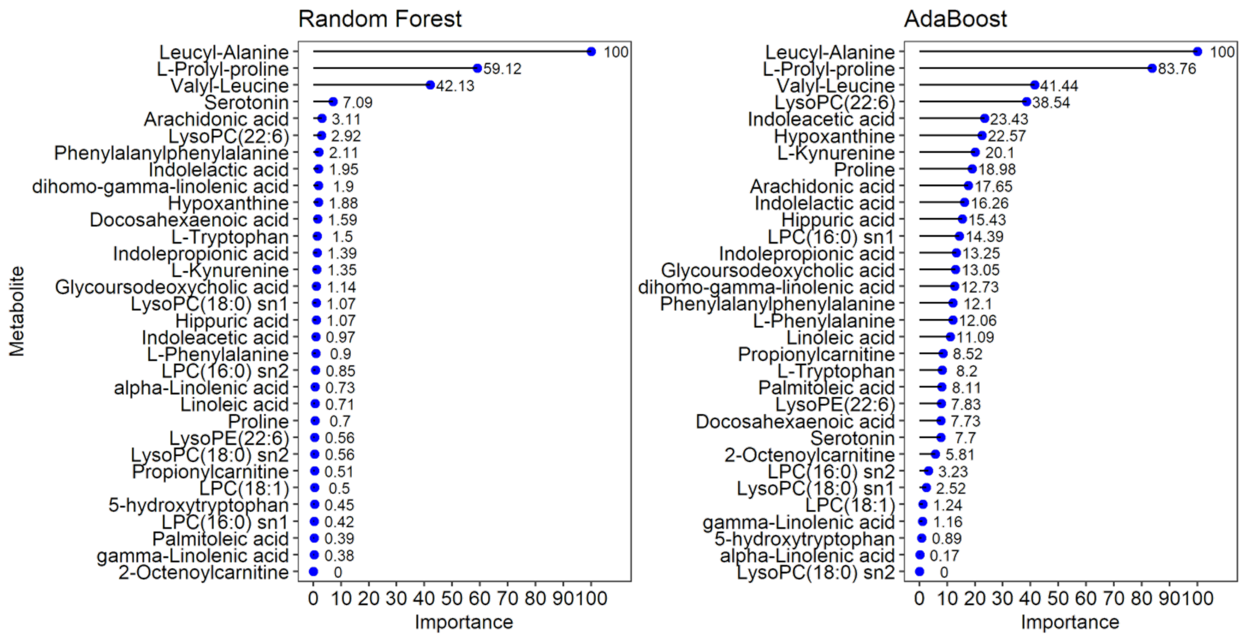


**Figure S5.** Metabolite importance plots for the random forest and adaBoost classification models.

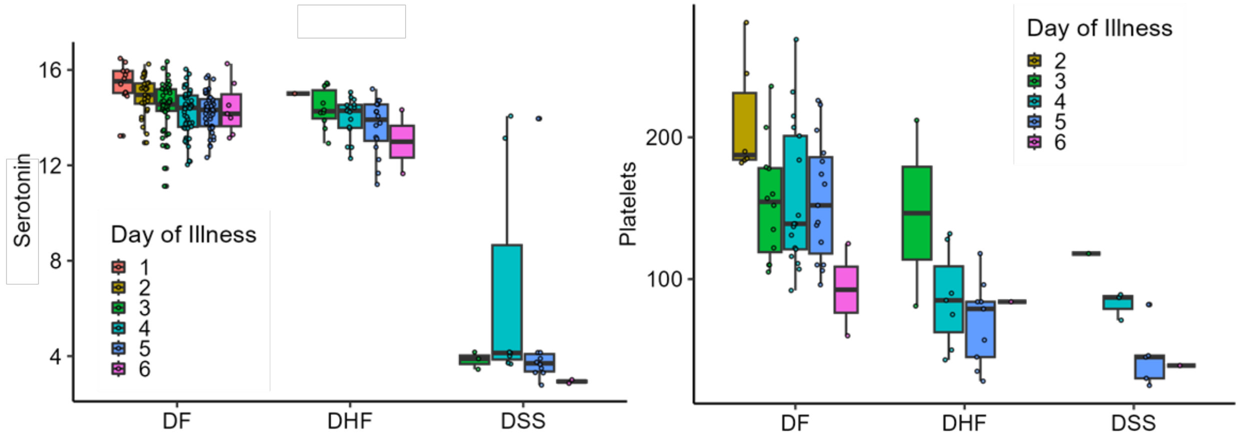


**Figure S6.** Left) Serum serotonin abundance as function of days of illness plotted for DF, DHF and DSS patients. Right) Serum platelets for a subset of 144 patients as a function of day of illness plotted for DF, DHF and DSS patents.

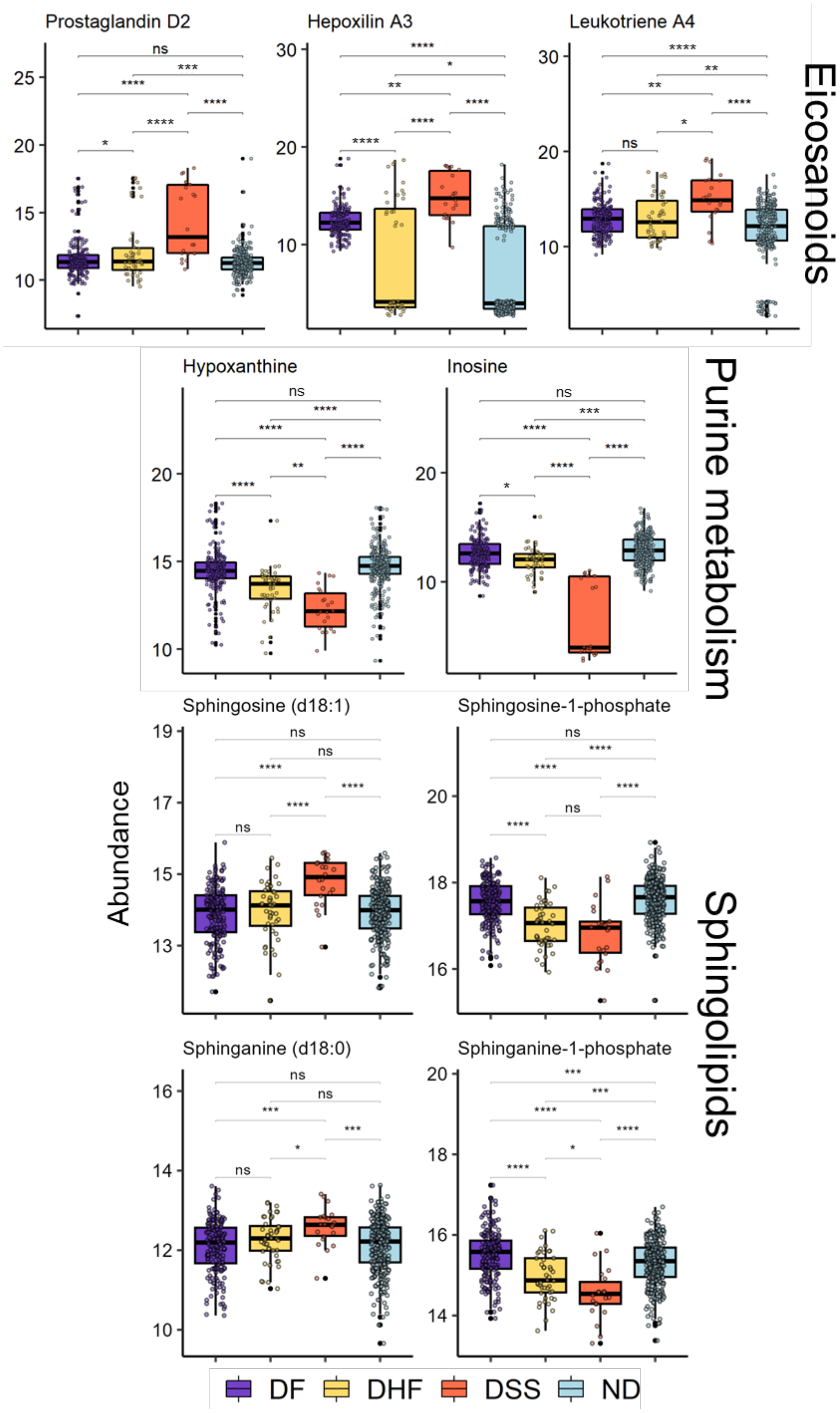
Prostaglandin D2
Hepoxilin A3
Leukotriene A4
Eicosanoids
Hypoxanthine
Inosine
Purine metabolism
Sphingosine (d18:1)
Sphingosine-1-phosphate
Sphinganine (d18:0)
Sphinganine-1-phosphate
Sphingolipids
Abundance
DF
DHF
DSS
ND

**Figure S7.** Abundance of n-6 derived eicosanoids, molecules related to purine metabolism and sphingolipids in sera of ND and dengue disease patients. Abundance is represented as the normalized and log2-transformed LC-MS peak area. Benjamini-Hochberg adjusted p-values generated using a moderated t-test were used to define statistical significance level; $p < 0.0001$, 0.001, 0.01, 0.05 were each represented by ****, ***, **, or *, respectively.

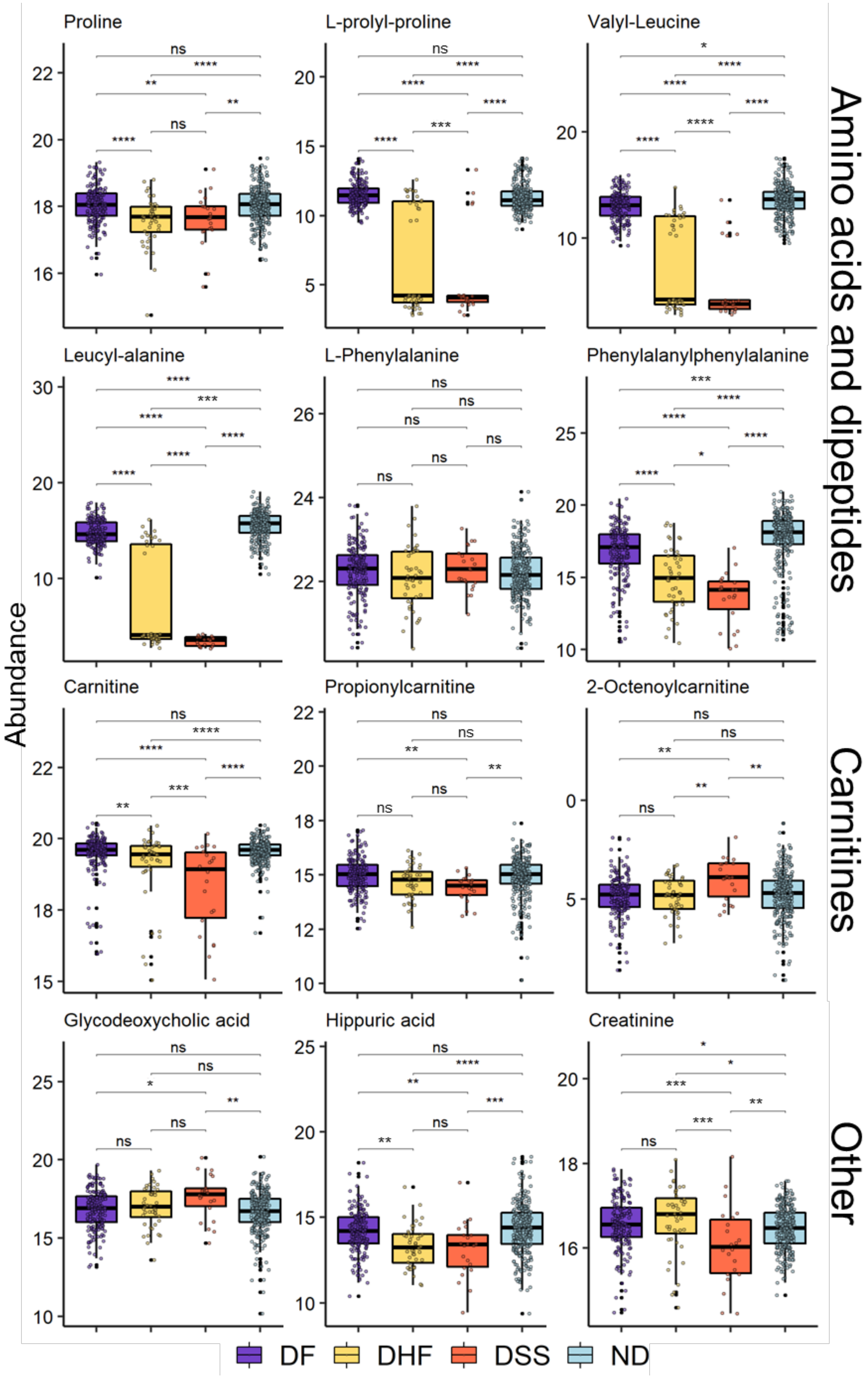
Proline
L-prolyl-proline
Valyl-Leucine
Leucyl-alanine
L-Phenylalanine
Phenylalanylphenylalanine
Carnitine
Propionylcarnitine
2-Octenoylcarnitine
Glycodeoxycholic acid
Hippuric acid
Creatinine
Abundance
Amino acids and dipeptides
Carnitines
Other
DF
DHF
DSS
ND

**Figure S8.** Abundance of amino acids, dipeptides, carnitines and other metabolites in sera of ND and dengue disease patients. Abundance is represented as the normalized and log2-transformed LC-MS peak area. Benjamini-Hochberg adjusted p-values generated using a moderated t-test were used to define statistical significance level; $p < 0.0001$, 0.001, 0.01, 0.05 were each represented by ****, ***, **, or *, respectively.

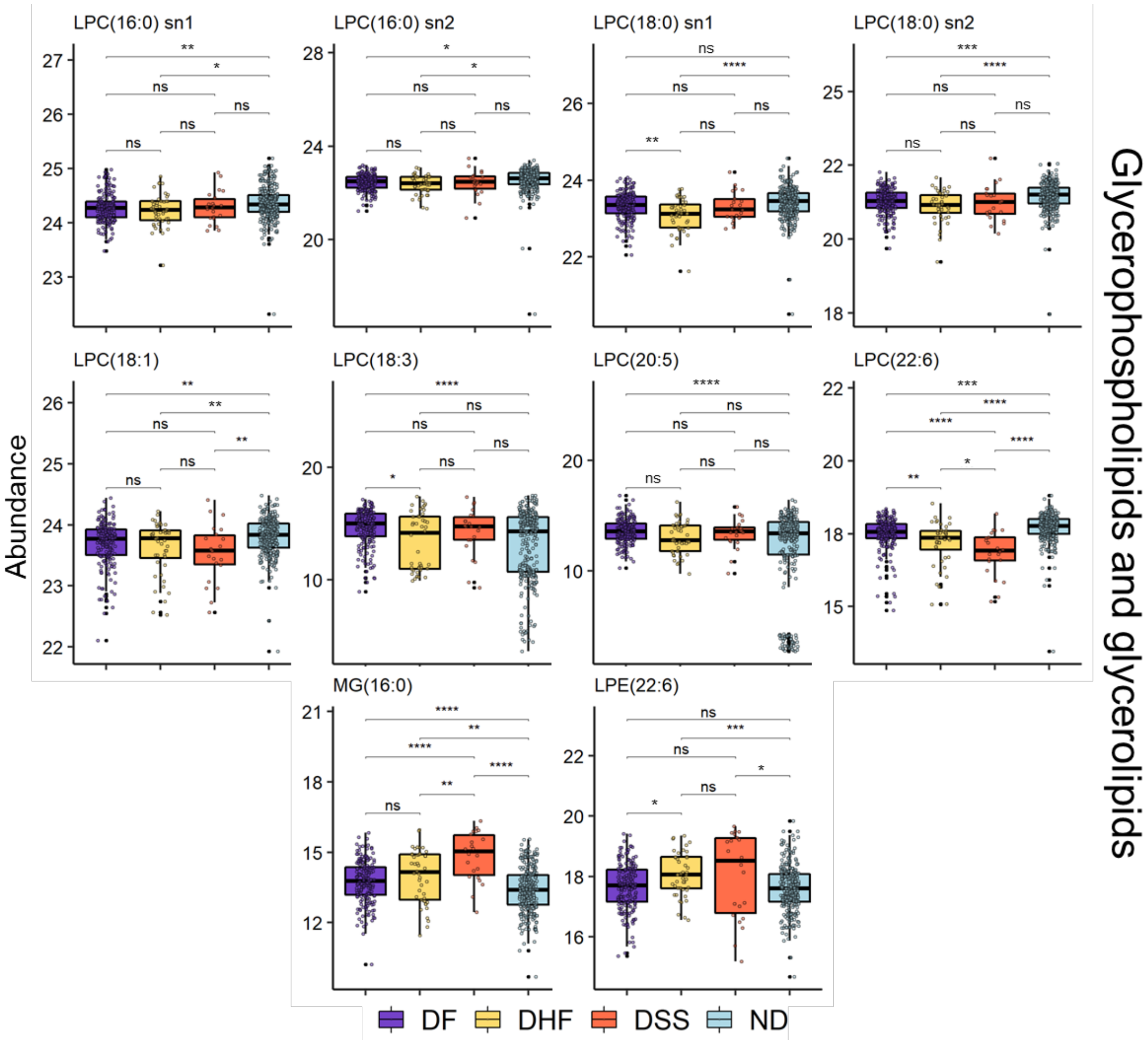


**Figure S9.** Abundance of glycerophospholipids and glycerolipids in sera of ND and dengue disease patients. Abundance is represented as the normalized and log2-transformed LC-MS peak area. Benjamini-Hochberg adjusted p-values generated using a moderated t-test were used to define statistical significance level; $p < 0.0001$, 0.001, 0.01, 0.05 were each represented by ****, ***, **, or *, respectively.

**Table S4**. Median t-score (medt) and Fischer's exact test p-value (FET) for all proposed metabolic pathways and dengue disease state comparisons (see next page)

| Pathway | medt_DS SvDF | FET_DS SvDF | medt_DSS vDHF | FET_DSS vDHF | medt_DH FvDF | FET_DH FvDF |
|---|---|---|---|---|---|---|
| TCA cycle | 6.2232 | 0.7661 | 2.7674 | 0.7083 | 0.3532 | 0.1063 |
| Omega-3 fatty acid metabolism | 3.7358 | 0.3439 | 3.5329 | 0.0115 | -5.7500 | 0.7851 |
| Alkaloid biosynthesis II | 3.5342 | 0.9015 | 2.0582 | 0.8726 | NA | NA |
| Di-unsaturated fatty acid beta-oxidation | 3.2167 | 0.2985 | 3.2271 | 0.5149 | -2.3852 | 0.9202 |
| Linoleate metabolism | 3.1860 | 0.1523 | 3.4191 | 0.0002 | -5.6332 | 0.6118 |
| Fatty acid activation | 3.1160 | 0.0329 | 3.4634 | 0.0556 | -4.0297 | 0.9988 |
| De novo fatty acid biosynthesis | 2.9148 | 0.0023 | 3.5077 | 0.0028 | -5.6738 | 0.9950 |
| Fatty Acid Metabolism | 2.9148 | 0.0805 | 3.1396 | 0.2756 | -2.3829 | 0.9695 |
| D4&E4-neuroprostanes formation | 2.7847 | 0.3217 | 4.3481 | 0.2646 | -4.9074 | 0.4618 |
| Bile acid biosynthesis | 2.7665 | 0.8304 | 2.3400 | 0.7839 | -2.4072 | 0.9817 |
| Vitamin A (retinol) metabolism | 2.6739 | 0.0631 | 3.8457 | 0.0265 | -7.0931 | 0.2477 |
| Xenobiotics metabolism | 2.6598 | 0.8424 | 2.6466 | 0.8222 | -2.1342 | 0.7687 |
| Arachidonic acid metabolism | 2.6572 | 0.0047 | 3.6075 | 0.0037 | -5.5926 | 0.3777 |
| Ascorbate (Vitamin C) and Aldarate Metabolism | 2.6431 | 0.1028 | 2.4803 | 0.0695 | 2.5761 | 0.2965 |
| Leukotriene metabolism | 2.6405 | 0.0647 | 4.4315 | 0.1494 | -5.6332 | 0.3967 |
| Prostaglandin formation from dihomo gama-linoleic acid | 2.6344 | 0.2201 | 2.8770 | 0.1696 | -5.7365 | 0.9244 |
| Prostaglandin formation from arachidonate | 2.6284 | 0.8746 | 5.2683 | 0.3139 | -6.2063 | 0.5001 |
| Vitamin E metabolism | 2.5989 | 0.9376 | 2.4803 | 0.7369 | -3.1690 | 0.9734 |
| Glycerophospholipid metabolism | 2.5965 | 0.2152 | 2.9495 | 0.2041 | -3.1358 | 0.9530 |
| C21-steroid hormone biosynthesis and metabolism | 2.5191 | 0.9996 | 3.8457 | 0.9948 | -4.1688 | 0.0874 |
| Vitamin D3 (cholecalciferol) metabolism | 2.5138 | 0.7249 | 2.1700 | 0.5566 | -3.5442 | 0.9976 |
| Glycosphingolipid metabolism | 2.4956 | 0.6196 | 2.1464 | 0.7501 | -3.4129 | 0.9643 |
| Omega-6 fatty acid metabolism | 0.3558 | 0.2201 | 5.5403 | 0.1696 | -5.8220 | 0.9244 |

| Pathway | medt_DSSvDF | FET_DSSvDF | medt_DSSvDHF | FET_DSSvDHF | medt_DHFvDF | FET_DHFvDF |
|---|---|---|---|---|---|---|
| Selenoamino acid metabolism | 0.3094 | 0.9640 | -2.1573 | 0.9424 | 2.4513 | 0.7801 |
| Putative anti-Inflammatory metabolites formation from EPA | -0.0671 | 0.2263 | -2.5885 | 0.8865 | -2.9021 | 0.3081 |
| Methionine and cysteine metabolism | -0.0671 | 0.9523 | -2.6819 | 0.9938 | -2.4072 | 0.5430 |
| Vitamin B6 (pyridoxine) metabolism | -2.3123 | 0.4985 | 0.2396 | 0.9424 | -2.5270 | 0.9605 |
| Lysine metabolism | -2.5737 | 0.2599 | -3.0751 | 0.8824 | -2.8685 | 0.3222 |
| Aspartate and asparagine metabolism | -2.6003 | 0.2215 | -3.0339 | 0.9667 | -2.7251 | 0.1984 |
| Tryptophan metabolism | -2.6659 | 0.5418 | -2.2170 | 0.9705 | -2.7495 | 0.1612 |
| Arginine and Proline Metabolism | -2.8137 | 0.2831 | -3.2003 | 0.9344 | -2.9526 | 0.0283 |
| Urea cycle_amino group metabolism | -2.8303 | 0.4892 | -3.3726 | 0.9979 | -2.9779 | 0.5070 |
| Glutathione Metabolism | -3.0271 | 0.9953 | -2.2840 | 0.9993 | -2.9087 | 0.1857 |
| Phosphatidylinositol phosphate metabolism | -3.0896 | 0.6126 | 0.0631 | 0.4637 | -2.5429 | 0.8558 |
| Carnitine shuttle | -3.2158 | 0.9295 | -0.1353 | 0.9247 | -2.4170 | 0.4009 |
| Butanoate metabolism | -3.5000 | 0.1702 | -3.2064 | 0.4769 | -2.4556 | 0.2278 |
| Tyrosine metabolism | -3.5000 | 0.7519 | -2.2840 | 0.9995 | -2.5983 | 0.0066 |
| Pyruvate Metabolism | -3.5656 | 0.4698 | -2.2840 | 0.4125 | -3.2262 | 0.7243 |
| Glycosylphosphatidylinositol(GPI)-anchor biosynthesis | -3.5904 | 0.6857 | -2.0865 | 0.6425 | NA | NA |
| Beta-Alanine metabolism | -3.7098 | 0.9063 | -3.3276 | 0.9978 | -2.9248 | 0.4309 |
| Vitamin B9 (folate) metabolism | -3.7342 | 0.9867 | -13.9727 | 0.9947 | -2.5728 | 0.7851 |
| Valine, leucine and isoleucine degradation | -3.7593 | 0.1320 | -3.2064 | 0.2008 | -2.4801 | 0.3212 |
| Benzoate degradation via CoA ligation | -3.7593 | 0.3217 | -2.1832 | 0.2646 | NA | NA |
| Glycine, serine, alanine and threonine metabolism | -3.7811 | 0.9215 | -3.2064 | 0.9835 | -2.5983 | 0.6303 |
| Drug metabolism - cytochrome P450 | -3.8136 | 0.0499 | -2.7713 | 0.0251 | -3.1888 | 0.4251 |
| Purine metabolism | -3.8477 | 0.7919 | -4.0365 | 0.9749 | -2.5696 | 0.0229 |
| Sialic acid metabolism | -4.0681 | 0.4478 | -6.8229 | 0.3248 | -2.4436 | 0.9013 |

| Pathway | medt_DSSvDF | FET_DSSvDF | medt_DSSvDHF | FET_DSSvDHF | medt_DHFvDF | FET_DHFvDF |
|---|---|---|---|---|---|---|
| Hexose phosphorylation | -4.0681 | 0.8182 | -6.8229 | 0.9424 | 0.1496 | 0.7801 |
| Vitamin B3 (nicotinate and nicotinamide) metabolism | -4.1657 | 0.8258 | -4.0748 | 0.9947 | -2.6159 | 0.0378 |
| C5-Branched dibasic acid metabolism | -4.2021 | 0.6857 | -2.7452 | 0.6425 | -2.9350 | 0.4746 |
| Dimethyl-branched-chain fatty acid mitochondrial beta-oxidation | -4.3519 | 0.3217 | -15.6776 | 0.9547 | -3.4059 | 0.4618 |
| Nitrogen metabolism | -4.3958 | 0.7661 | -4.0748 | 0.9547 | -2.5759 | 0.1063 |
| Vitamin B5 - CoA biosynthesis from pantothenate | -4.5662 | 0.6264 | -3.2387 | 0.5496 | -1.9931 | 0.9244 |
| CoA Catabolism | -4.5662 | 0.6264 | -3.2387 | 0.5496 | -1.9931 | 0.9244 |
| Propanoate metabolism | -4.8386 | 0.2201 | -2.7452 | 0.5496 | -3.2262 | 0.2747 |
| Squalene and cholesterol biosynthesis | -4.8386 | 0.7587 | -2.9371 | 0.4038 | -2.5474 | 0.4309 |
| Histidine metabolism | -4.9295 | 0.9691 | -4.7241 | 0.9878 | -2.5110 | 0.4373 |
| Saturated fatty acids beta-oxidation | -4.9781 | 0.2175 | -4.1698 | 0.3248 | -6.4372 | 0.9951 |
| Alanine and Aspartate Metabolism | -5.0181 | 0.8842 | -4.0748 | 0.9878 | -2.6518 | 0.2048 |
| Fructose and mannose metabolism | -5.0439 | 0.8182 | -6.8229 | 0.7538 | -2.6651 | 0.1564 |
| Glutamate metabolism | -5.0806 | 0.7803 | -3.3726 | 0.8865 | -2.5759 | 0.1122 |
| Pyrimidine metabolism | -5.0806 | 0.9523 | -3.3726 | 0.7724 | -2.5504 | 0.1462 |
| Caffeine metabolism | -5.4329 | 0.9057 | -2.3897 | 0.5496 | -2.6865 | 0.2747 |
| Mono-unsaturated fatty acid beta-oxidation | -5.5568 | 0.6857 | -4.4978 | 0.6425 | NA | NA |
| Pentose phosphate pathway | -5.9188 | 0.7661 | -6.8229 | 0.7083 | -2.6865 | 0.8556 |
| Ubiquinone Biosynthesis | -6.0115 | 0.7661 | -4.4713 | 0.7083 | -2.3797 | 0.8556 |
| N-Glycan biosynthesis | -6.0570 | 0.7661 | -3.5480 | 0.7083 | -2.6369 | 0.4618 |
| Biopterin metabolism | -6.1542 | 0.6146 | -3.0892 | 0.9403 | -3.5281 | 0.4443 |
| Porphyrin metabolism | -6.2854 | 0.3104 | -4.6317 | 0.6187 | -3.0575 | 0.7303 |
| Glycolysis and Gluconeogenesis | -6.6605 | 0.6146 | -8.4415 | 0.5149 | -2.6437 | 0.1857 |
| Aminosugars metabolism | -6.8516 | 0.8842 | -5.1690 | 0.9421 | -2.5535 | 0.2048 |

| Pathway | medt_DS SvDF | FET_DS SvDF | medt_DSS vDHF | FET_DSS vDHF | medt_DH FvDF | FET_DH FvDF |
|---|---|---|---|---|---|---|
| Hyaluronan Metabolism | -6.9527 | 0.6857 | -5.1690 | 0.6425 | NA | NA |
| Keratan sulfate degradation | -6.9527 | 0.8182 | -5.9960 | 0.7538 | -2.6865 | 0.4523 |
| N-Glycan Degradation | -6.9527 | 0.8182 | -5.9960 | 0.7538 | -2.6865 | 0.4523 |
| Glycosphingolipid biosynthesis - ganglioseries | -6.9527 | 0.8558 | -11.3972 | 0.5149 | -2.7752 | 0.7308 |
| Galactose metabolism | -7.0961 | 0.4478 | -4.9466 | 0.3248 | -2.6651 | 0.5430 |
| Vitamin B12 (cyanocobalamin) metabolism | -7.2395 | 0.6857 | -4.7241 | 0.6425 | -2.3797 | 0.4746 |
| Chondroitin sulfate degradation | -7.6149 | 0.7661 | -6.8229 | 0.7083 | -2.6865 | 0.8556 |
| Glycosphingolipid biosynthesis - globoseries | -7.6149 | 0.7661 | -6.8229 | 0.7083 | -2.6865 | 0.8556 |
| Heparan sulfate degradation | -7.6149 | 0.7661 | -6.8229 | 0.7083 | -2.6865 | 0.8556 |
| Starch and Sucrose Metabolism | -8.2770 | 0.8182 | -4.8244 | 0.7538 | -12.1658 | 0.7801 |
| Androgen and estrogen biosynthesis and metabolism | -8.7814 | 0.9971 | -3.7750 | 0.7185 | -2.8916 | 0.2501 |
| Dynorphin metabolism | -10.1708 | 0.4985 | -13.2464 | 0.7538 | -3.2208 | 0.9605 |
| Drug metabolism - other enzymes | -10.6025 | 0.1028 | -11.9780 | 0.3041 | -3.3373 | 0.2965 |
| Polyunsaturated fatty acid biosynthesis | -12.0431 | 0.4698 | -9.5961 | 0.4125 | -2.1047 | 0.7243 |
| Vitamin B1 (thiamin) metabolism | -12.1115 | 0.3217 | -8.9501 | 0.2646 | -2.1677 | 0.8556 |
| Lipoate metabolism | -14.4744 | 0.9015 | -3.8755 | 0.8726 | -13.4285 | 0.7243 |
| Phytanic acid peroxisomal oxidation | -15.7168 | 0.4698 | -3.3796 | 0.4125 | -5.3265 | 0.7243 |
| Fatty acid oxidation, peroxisome | -18.8380 | 0.4698 | -3.3796 | 0.4125 | -3.7156 | 0.2248 |
| Fatty acid oxidation | -23.8025 | 0.3217 | -20.5256 | 0.2646 | -3.9729 | 0.4618 |
| R Group Synthesis | -24.2646 | 0.4698 | -20.9081 | 0.4125 | -2.1047 | 0.7243 |

1. mzCloud – Advanced Mass Spectral Database (available at https://www.mzcloud.org/).

2. J. K. Pauling, M. Hermansson, J. Rgen Hartler, K. Christiansen, S. F. Gallego, B. Peng, R. Ahrends, C. S. Ejsing, Proposal for a common nomenclature for fragment ions in mass spectra of lipids. (2017), doi:10.1371/journal.pone.0188394.

3. UWPR (available at https://proteomicsresource.washington.edu/cgi-bin/fragment.cgi).

10. R. Wehrens, J. A. Hageman, F. Van Eeuwijk, R. Kooke, • Pádraic, J. Flood, E. Wijnker, J. J. B. Keurentjes, A. Lommen, • Henriëtte, D. L. M. Van Eekelen, • Robert, D. Hall, R. Mumm, • Ric, C. H. De Vos, Improved batch correction in untargeted MS-based metabolomics. *Metabolomics* **12**, doi:10.1007/s11306-016-1015-8.

11. D. J. Stekhoven, P. Bühlmann, MissForest—non-parametric missing value imputation for mixed-type data. *Bioinformatics* **28**, 112–118 (2012).

12. A. J. Hackstadt, A. M. Hess, Filtering for increased power for microarray data analysis. *BMC Bioinformatics* **10**, 11 (2009).

13. S. Li, Y. Park, S. Duraisingham, F. H. Strobel, N. Khan, Predicting Network Activity from High Throughput Metabolomics. *PLoS Comput Biol* **9**, 1003123 (2013).

1. mzCloud – Advanced Mass Spectral Database (available at https://www.mzcloud.org/).

2. J. K. Pauling, M. Hermansson, J. Rgen Hartler, K. Christiansen, S. F. Gallego, B. Peng, R. Ahrends, C. S. Ejsing, Proposal for a common nomenclature for fragment ions in mass spectra of lipids. (2017), doi:10.1371/journal.pone.0188394.

3. UWPR (available at https://proteomicsresource.washington.edu/cgi-bin/fragment.cgi).

10. R. Wehrens, J. A. Hageman, F. Van Eeuwijk, R. Kooke, • Pádraic, J. Flood, E. Wijnker, J. J. B. Keurentjes, A. Lommen, • Henriëtte, D. L. M. Van Eekelen, • Robert, D. Hall, R. Mumm, • Ric, C. H. De Vos, Improved batch correction in untargeted MS-based metabolomics. *Metabolomics* **12**, doi:10.1007/s11306-016-1015-8.

11. D. J. Stekhoven, P. Bühlmann, MissForest—non-parametric missing value imputation for mixed-type data. *Bioinformatics* **28**, 112–118 (2012).

12. A. J. Hackstadt, A. M. Hess, Filtering for increased power for microarray data analysis. *BMC Bioinformatics* **10**, 11 (2009).

13. S. Li, Y. Park, S. Duraisingham, F. H. Strobel, N. Khan, Predicting Network Activity from High Throughput Metabolomics. *PLoS Comput Biol* **9**, 1003123 (2013).
